# Fast and sensitive GCaMP calcium indicators for imaging neural populations

**DOI:** 10.1101/2021.11.08.467793

**Authors:** Yan Zhang, Márton Rózsa, Yajie Liang, Daniel Bushey, Ziqiang Wei, Jihong Zheng, Daniel Reep, Gerard Joey Broussard, Arthur Tsang, Getahun Tsegaye, Sujatha Narayan, Christopher J. Obara, Jing-Xuan Lim, Ronak Patel, Rongwei Zhang, Misha B. Ahrens, Glenn C. Turner, Samuel S.-H. Wang, Wyatt L. Korff, Eric R. Schreiter, Karel Svoboda, Jeremy P. Hasseman, Ilya Kolb, Loren L. Looger

## Abstract

Calcium imaging with protein-based indicators is widely used to follow neural activity in intact nervous systems. The popular GCaMP indicators are based on the calcium-binding protein calmodulin and the RS20 peptide. These sensors report neural activity at timescales much slower than electrical signaling, limited by their biophysical properties and trade-offs between sensitivity and speed. We used large-scale screening and structure-guided mutagenesis to develop and optimize several fast and sensitive GCaMP-type indicators. The resulting ‘jGCaMP8’ sensors, based on calmodulin and a fragment of endothelial nitric oxide synthase, have ultra-fast kinetics (rise times, 2 ms) and still feature the highest sensitivity for neural activity reported for any protein-based sensor. jGCaMP8 sensors will allow tracking of larger populations of neurons on timescales relevant to neural computation.

## Introduction

Measurement of Ca^2+^-dependent fluorescence using genetically encoded calcium indicators (GECIs) is one of the most widely used methods for tracking neural activity in defined neurons and neural networks ^1^. Recent advances have been driven in a virtuous cycle of new methods for *in vivo* microscopy ^2–4^ and engineered GECIs with higher response and sensitivity. In particular, the green fluorescent protein- (GFP-) based GCaMP sensors ^5–8^ have been iteratively engineered to enhance the signal-to-noise ratio (SNR) for detection of Ca^2+^ entering neurons through voltage-gated channels. The widely used GCaMP6 ^8^ and jGCaMP7 ^6^ sensors enable detection of single action potentials under favorable conditions. They can be used to monitor the activity of large groups of neurons using two-photon microscopy and wide-field fluorescence imaging ^2^. They have also been used to measure activity-induced calcium changes in small subcellular compartments such as dendritic spines ^8–11^ and axons ^12, 13^.

Electrical signals propagate through neural circuits over timescales of milliseconds. Determining how the activity of one set of neurons influences another and ultimately animal behavior requires tracking activity on concomitant time scales. Action potentials produce essentially delta function-like calcium currents, yielding large, rapid (<1 ms) increases in cytoplasmic calcium ^14^. Free calcium ions typically bind fluorescent calcium indicators very rapidly. For example, millisecond timescale detection of action potentials has been demonstrated with synthetic fluorescence calcium indicators *in vivo* ^15–17^. However, the kinetics of GECI fluorescence change is limited by sensor biophysics; for instance, in response to single action potentials in pyramidal neurons GCaMPs have fluorescent rise-times (50%) on the order of 100 ms ^6–8, 18–20^. Consequently, GCaMPs are often used to map relatively static representations of neural information, rather than tracking the rich dynamics in neural circuits ^21, 22^.

Previous attempts to improve GCaMP kinetics have been only partially successful. Among the GCaMP6 and jGCaMP7 indicators, the “f” (fast) variants were optimized for kinetics. They have risetimes of ∼50 ms, but with reduced sensitivity compared to their slower siblings (“s”, sensitive variants). Generally, attempts to improve ΔF/F_0_ are associated with a slowing of kinetics ^6, 8, 19, 20^. The mechanisms underlying this trade-off are not simply due to binding affinity for Ca^2+^ and in general are not well-understood. For example, the kinetics are sensitive to mutation of the RS20-CaM interface, far from the Ca^2+^-binding EF hands ^8, 19, 20^. In addition to these point mutants, the RS20 peptide has been swapped for that from CaM-dependent kinase kinase CaMKK-α/β (ckkap peptide) in the “XCaMP” sensors ^23^ – as well as the red GECIs R-CaMP2 ^24^ (actually an RGECO variant) and K-GECO1 ^25^ – with mixed effects on affinity, kinetics, and ΔF/F_0_.

Here we present bright, sensitive GCaMP sensors with dramatically improved kinetics. jGCaMP8 sensors include: jGCaMP8s (fast rise, slow decay, sensitive), jGCaMP8f (fast rise, fast decay), jGCaMP8m (medium decay). All jGCaMP8 sensors have nearly 10x faster fluorescence rise-times than previous GCaMPs and can track individual spikes in neurons with spike rates of ∼50 Hz. jGCaMP8 sensors are also more linear than previous GCaMPs, allowing robust deconvolution for spike extraction. The jGCaMP8 sensors were tested *in vivo* in mice, flies and fish, and were found to provide better performance across all metrics relevant to imaging neural populations *in vivo*.

## Results

### Sensor design and optimization

Various calmodulin-binding peptides (**Supp. Table 1**) were chosen from the Protein Data Bank and cloned into GCaMP6s in place of the RS20 peptide. Based on fast kinetics, saturating ΔF/F_0_, apparent *K_d_*, Hill coefficient, and apparent brightness, we prioritized variants based on peptides from endothelial nitric oxide synthase (PDB 1NIW; peptide “ENOSP”) and death-associated protein kinase 1 (1YR5; peptide “DAPKP”) for optimization (**Methods**). The two linkers ^26^ were systematically mutated, and sensors were screened for high signal change and retained fast kinetics. ∼35 promising sensors were then tested in response to action potentials (APs) elicited in cultured neurons in 96-well plates (**Methods**). Action potentials produce essentially instantaneous increases in calcium ^14^ and are therefore ideal to screen for GECIs with fast kinetics ^27^. Fluorescence changes were extracted from multiple single neurons per well. Sensors were evaluated according to several properties (**Supp. Table 2**): sensitivity (response to 1 AP), dynamic range (response to a high-frequency train of 160 APs), kinetics (rise and decay times), and baseline brightness. Sensors based on DAPKP showed fast decay time and good sensitivity compared to jGCaMP7f – but with slow rise times (**Supp. Table 2**). Sensors with ENOSP had similar sensitivity and significantly faster rise and decay times than jGCaMP7f.

We prioritized ENOSP-based sensors for further optimization. ENOSP variant jGCaMP8.410.80 (linker 1 Leu-Lys-Ile) showed 1.8-fold faster half-rise time and 4.4-fold faster half-decay time than jGCaMP7f, with similar resting brightness and dynamic range, but 35% lower 1-AP response. We solved the crystal structure of jGCaMP8.410.80 (**Fig. 1A, Supp. Fig. 1A, Supp. Table 3**). The structure is similar to previous GCaMPs; the major differences are at the 3-way interface between cpGFP, CaM, and the new ENOSP peptide; the slight twist of the ENOSP peptide relative to the RS20 peptide; and at the first helix of EF-hand 1 (**Supp. Fig. 1A**). In jGCaMP8.410.80, Ile 32 (occupying the same space as GCaMP5G-Glu60) packs closely (both hydrophobic and C-H/*π* interactions) with Tyr352 (Tyr380 in GCaMP5G), allowing the tyrosine to penetrate more deeply into the cpGFP-CaM interface, and forming a water-mediated hydrogen bond network with the chromophore (**Supp. Fig. 1B**). This tyrosine was the core improvement in GCaMP5 ^26^ – increasing brightness and ΔF/F_0_ – the ENOSP peptide appears to further facilitate this interaction. Guided by the structure, we targeted interface sites (**Supp. Fig. 1C**) for site-saturation mutagenesis and tested the variants in cultured neurons for higher sensitivity and retained fast kinetics in detecting spikes. Several single mutations improved properties (**Supp. Table 2**), particularly residues near the ENOSP C-terminus and the cpGFP-CaM interface. Beneficial point mutations were combined in subsequent rounds of screening.^28, 29^

**Figure 1.**
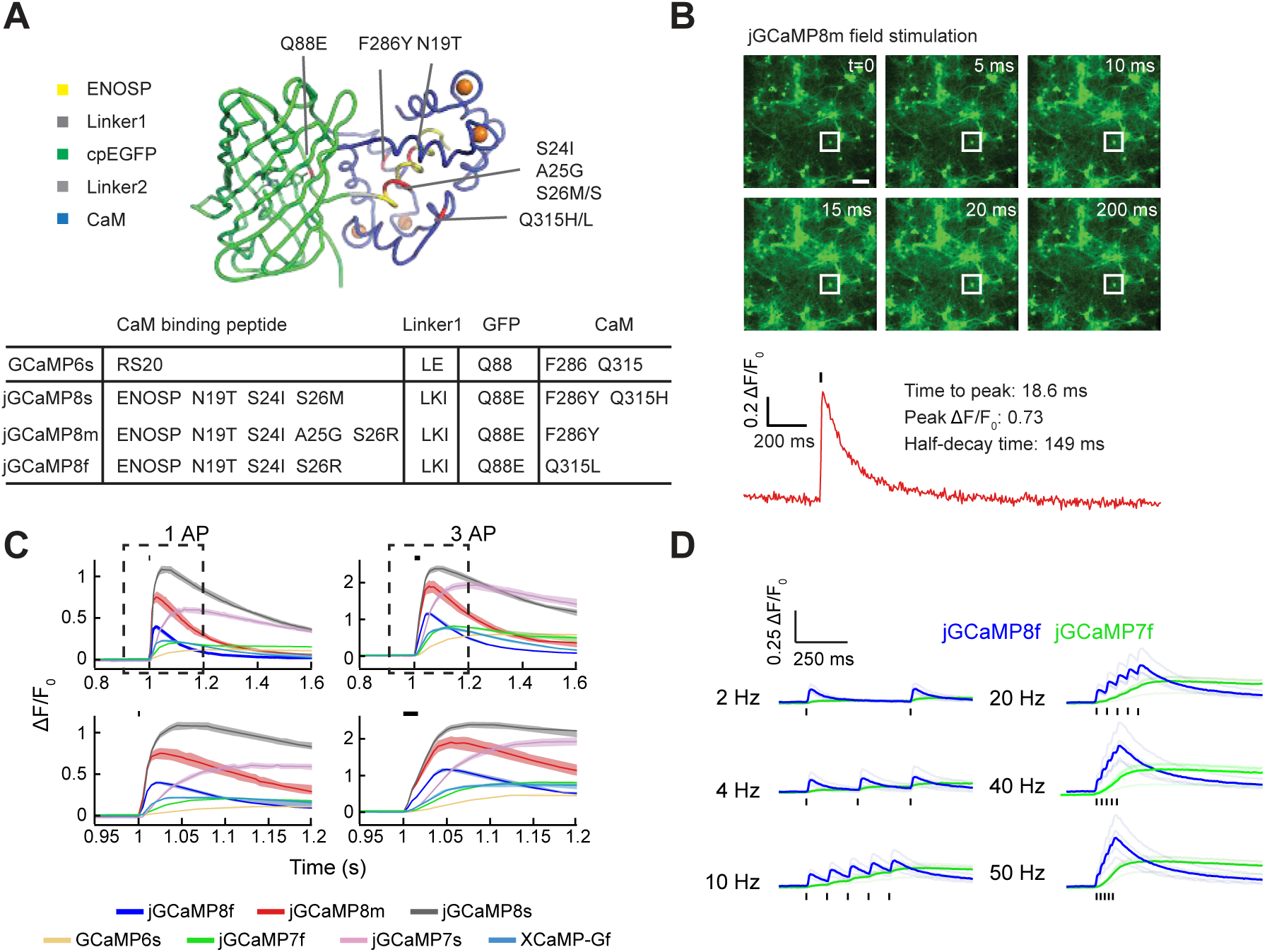
GCaMP mutagenesis and screening. **A.** jGCaMP8.410.80 structure and mutations in different jGCaMP8 variants relative to 6s (top). ENOSP (yellow), linker 1 (ENOSP-cpGFP; dark grey), linker 2 (cpGFP-CaM; light grey), cpGFP (green), CaM (blue), mutated sites (red), Ca^2+^ ions (orange). Bottom table, mutations for each jGCaMP8 variant. **B.** Top: representative frames of an 8m FOV in the field stimulation assay. Bottom: single-trial fluorescence trace and sensor response to 1-AP stimulation of the neuron outlined in the top panel. Scale bar: 100 µm. **C.** Top: responses to 1 and 3 field stimuli (black bars). Bottom: zoomed-in insets from the top panel (dashed boxes) to highlight rise and decay kinetics. Solid lines: mean, shaded area: s.e.m. (8f: 48 wells, 1,696 neurons; 8m: 11 wells, 496 neurons; 8s: 24 wells, 1,183 neurons; 6s: 859 wells, 24,998 neurons; 7f: 950 wells, 26,679 neurons; 7s: 22 wells, 514 neurons; XCaMP-Gf: 69 wells, 1,305 neurons; overall statistics: 23 independent transfections, 130 96-well plates). **D.** 8f and 7f responses to field stimulation pulses of increasing frequencies imaged in neuronal culture under two-photon illumination. Light traces: individual neurons (n=4 for each sensor); dark traces: means. Black bars: stimulation pulses.

Mutagenesis and screening in neurons covered 776 total sensor variants, of which 683 (88%) produced detectable responses to 1 AP (**Supp. Fig. 2**, **Supp. Table 2**). Kinetics were improved relative to the previous fast sensor jGCaMP7f. Specifically, compared to jGCaMP7f, the half-rise time (*t_rise_*_,1/2_) was significantly shorter in 48% of screened variants, the time-to-peak fluorescence (*t_peak_*) was significantly shorter in 47%, and the half-decay time (*t_decay_*_,1/2_) was significantly shorter in 40%. Sensitivity (1-AP ΔF/F_0_) was higher than jGCaMP7f in 19%, and only 2% of variants had increased saturation response (160-AP ΔF/F_0_). Together, the mutagenesis produced a large set of variants with significant improvement in kinetics and sensitivity (**Supp. Table 2**).

### jGCaMP8 characterization

Three high-performing “jGCaMP8” variants were selected for additional characterization (**Fig. 1B-D, Supp. Fig. 3**). jGCaMP8f (“fast”) exhibited 1-AP *t_rise_*_,1/2_ of 7.0±0.7 ms, and 1-AP *t_peak_* of 24.9±6.0 ms, more than 3- and 5-fold shorter than jGCaMP7f, respectively. We note that the rise-time measurements in cultured neurons were limited by the frame rate of the camera (200 Hz) and thus constitute an overestimate. jGCaMP8s (“sensitive”) exhibited 1-AP ΔF/F_0_ of 1.1±0.2, and 1-AP signal-to-noise ratio (SNR) of 41.3±10.4, approximately twice that of the most sensitive GECI to date, jGCaMP7s. jGCaMP8m (“medium”) is a useful compromise between sensitivity and kinetics: it exhibits 1-AP ΔF/F_0_ and 1-AP SNR comparable to jGCaMP7s, and kinetics comparable to jGCaMP8f, with the exception of a slower half-decay time (*t_decay_*_,1/2_, 134±14 *vs*. 92±22 ms; **Fig. 1C**). The fast kinetics and high sensitivity of the jGCaMP8 indicators allowed resolution of electrically evoked spikes at frequencies of up to 40 Hz (**Fig. 1D**). When stimulated with short bursts consisting of 3 and 10 APs, the jGCaMP8 sensors retained fast kinetics and high sensitivity. Overall, the jGCaMP8 series exhibited significant, multi-fold improvements across several parameters over previous GECIs.

We compared the jGCaMP8 sensors to the XCaMP series (green XCaMP variants XCaMP-G, XCaMP-Gf, and XCaMP-Gf_0_ ^23^, side-by-side in cultured neurons. The 1-AP ΔF/F_0_ was significantly higher for all jGCaMP8 sensors; the 1-AP SNR was significantly higher for jGCaMP8m and jGCaMP8s, 1-AP *t_rise_*_,1/2_ was significantly shorter for all jGCaMP8 sensors, 1- AP *t_peak_* was significantly shorter for jGCaMP8f and jGCaMP8m, and *t_decay_*_,1/2_ was significantly shorter for jGCaMP8f, when evaluated against all XCaMP sensors (**Supp. Fig. 3, Supp. Table 4**). The baseline fluorescence of the jGCaMP8 series was similar to jGCaMP7f, and significantly higher than the XCaMP sensors (**Supp. Fig. 4**). Photobleaching was also similar between jGCaMP7f and the jGCaMP8 sensors (**Supp. Fig. 5**).

GECIs with linear (*i.e.*, Hill coefficient ∼1) fluorescence responses to AP trains provide a larger effective dynamic range for quantifying spike rates and facilitate applications such as counting spikes within trains. We tested GCaMP sensors with bursts (83 Hz) containing different numbers (1-40) of action potentials. Given their higher sensitivity, 8m and 8s showed saturation behavior for smaller numbers of spikes compared to jGCaMP7. However, they behaved nearly linearly up to 10 spikes (**Supp. Fig. 6**). Finally, fluorescence recovery after photobleaching (FRAP) revealed that the jGCaMP8 variants showed similar diffusion in neurons compared to previous GECIs ^27^ (**Supp. Fig. 7A-C**) and independent of calcium (**Supp. Fig. 7D**), suggesting that they do not have altered cellular interactions.

### Imaging in Drosophila adult visual system and larva

GCaMP responses to visual stimulation were compared in *Drosophila* laminar monopolar L2 neurons (**Fig. 2A**), part of the OFF-motion visual system ^28–30^. Imaging was performed where L2 dendrites connect to columns in medulla layer 2. These non-spiking neurons depolarize during light decrease and hyperpolarize during increase. Fluorescence responses to visual stimulation were measured in multiple single neurons in individual animals (**Fig. 2B, Supp. Fig. 8A**). XCaMP-Gf was too dim to image (**Supp. Fig. 8B-C**) and was excluded from further study. At light-dark and dark-light transitions, all jGCaMP8 variants showed significantly faster rise, and 8m showed faster decay, respectively, than 7f (**Fig. 2C,D**; half-rise times: 7f, 128±11 ms; 8f, 76±8; 8m, 58±6; 8s, 80±8 ms; decay times: 7f, 277±29 ms; 8m, 137±21). 8m and 8f also showed markedly larger ΔF/F_0_ than 7f following light-on (**Fig. 2B-C**). All three jGCaMP8 indicators revealed a negative off-response after light-off (*i.e.*, hyperpolarization below baseline), whereas 7f was too slow (**Fig. 2C**). Flies were next subjected to light on-off stimulation at frequencies from 0.5-30 Hz. 8m and 8f showed much higher spectral density than 8s across all frequencies, and higher than 7f above 2 Hz (**Supp. Fig. 8D**). Visual stimuli of progressively shorter lengths were shown, from 25 ms dark flash down to 4 ms. 8m and 8f showed higher ΔF/F_0_ at all stimulus lengths (**Supp. Fig. 8E-top**). Quantification of stimulus detection by the discriminability index d’ ^31^ showed that 8m and 8f provide markedly superior stimulus detection above noise than 7f and 8s across all stimulus frequencies (**Supp. Fig. 8E-bottom**). The jGCaMP8 variants were somewhat dimmer than 7f because of lower expression (**Supp. Fig. 8B-C, Supp. Fig. 9**) but were sufficiently bright to provide high-SNR imaging.

**Figure 2.**
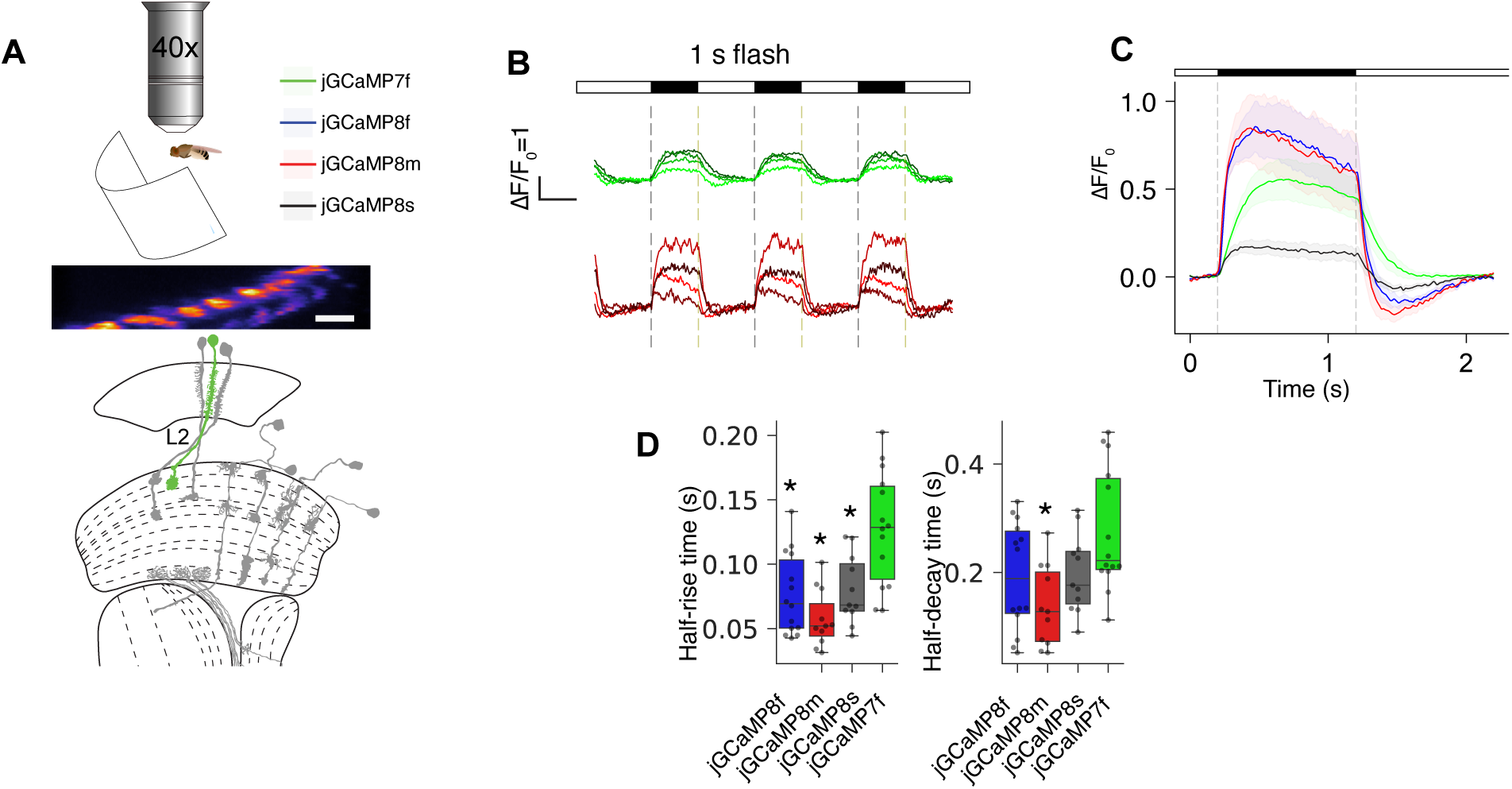
jGCaMP8 performance in *Drosophila*. **A.** Design of adult fly experiments. Top, fly receiving visual stimuli. Middle, fluorescence micrograph of L2 dendrites in medullar layer 2 (5 µm scale bar). Bottom, schematic of the *Drosophila* visual system (modified from ^57^). **B.** ΔF/F_0_ response to a 0.5 Hz visual stimulation frequency from variants 7f (top) and 8m (bottom). Individual traces show 4 representative individual animals per GECI (colors arbitrary), total *n* for each variant: 7f: 14 flies, 8f: 14, 8m: 11, 8s: 11. **C.** Mean ΔF/F_0_ response to 0.5 Hz stimulation. Dark line: mean, shaded area: s.e.m. Darkened visual stimulus between dashed lines. Total *n* for each variant: 10 trials per fly, mean of trials 2-10 recorded for each fly. **D.** Half-rise and half-decay times (box plots) for responses in C. Half-rise: 7f, 128±11 ms; 8f, 76±8; 8m, 58±6; 8s, 80±8 ms (Kruskal-Wallis multiple-comparison test, *P*=2.9E-4; pairwise Dunn’s comparison test with 7f, 8f: *P*=3.1e-3, 8m: 2.9e-5, 8s: 1.3e-2). Half-decay times: 7f, 277±29 ms; 8f, 192±26; 8m, 137±21; 8s, 198±21 ms (Kruskal-Wallis multiple-comparison test, *P*=2.4E-2; pairwise Dunn’s comparison test, 8f: *P*=1.1e-1, 8m: 2.2e-3, 8s: 1.8e-1). *: *P*<0.05.

Next, we imaged the GECIs at presynaptic boutons of the larval neuromuscular junction in response to precise electrical stimulation of motor axons (**Supp. Fig. 10**). jGCaMP8 variants showed large responses, with faster rise and decay times than 7f (**Supp. Fig. 10B, D, E**). The jGCaMP8 series detect individual stimuli much better than 7f at low frequencies and easily resolve spikes in 20 Hz trains (**Supp. Fig. 10H**), whereas previous GECIs cannot.

### Imaging in zebrafish optic tectum

We also imaged 8f, 7f, and 6f in zebrafish as histone H2B fusions (which improves cell segmentation ^32^; **Methods**). Transgenic Tg(*elavl3*:H2B-8f), Tg(*elavl3*:H2B-7f) and Tg(*elavl3*:H2B-6f) larval zebrafish were shown flashing visual stimuli and GCaMP transients were imaged in the optic tectum. 8f showed markedly faster rise (**Supp. Fig. 11E**) and decay (**Supp. Fig. 11F**) times than its predecessors.

### Imaging neural populations in mouse primary visual cortex

We next tested the jGCaMP8 sensors in L2/3 pyramidal neurons of mouse primary visual cortex (V1). We made a craniotomy over V1 and infected neurons with adeno-associated virus (AAV2/1-*hSynapsin-1*) ^33^(**Methods**) encoding jGCaMP8 variants, 7f ^6^, or XCaMP-Gf ^23^. After three weeks of expression, mice were lightly anesthetized and mounted under a custom two-photon microscope. Full-field, high-contrast drifting gratings were presented in each of eight directions to the contralateral eye (**Fig. 3A**). Two-photon imaging (30 Hz) was performed of L2/3 somata and neuropil (**Methods**).

**Figure 3.**
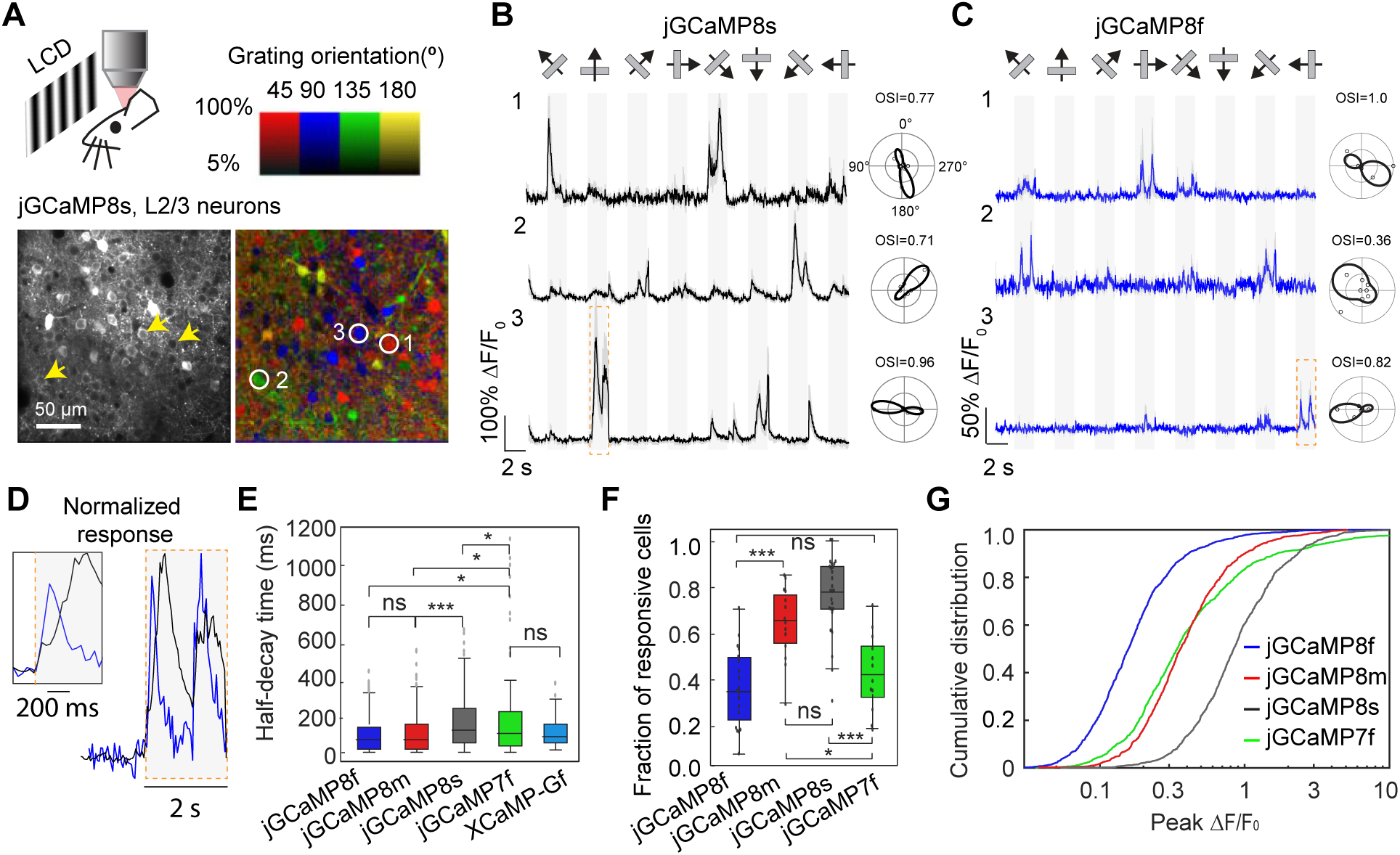
Imaging neural population in the mouse primary visual cortex (V1) *in vivo*. A. Top, schematic of the experiment. Bottom, example image of V1 L2/3 cells expressing 8s (left), and the same field of view color-coded based on the neurons’ preferred orientation (hue) and response amplitude (brightness). Color scale bar at top. B,C. Example traces from three L2/3 neurons expressing 8s (B, same cells as indicated in A) or 8f (C). Averages of five trials with shaded s.e.m. shown. Stimuli shown above traces. The preferred stimulus is the direction evoking the largest response. Polar plots indicate the preferred direction of cells. The orientation selectivity index (OSI) is displayed above each polar plot. D. Zoomed fluorescence traces corresponding to orange boxes in B (black, 8s) and C (blue, 8f), normalized to the peak of the response. Inset shows a further zoomed view of the first transient. E. Half-decay time of the fluorescence response after the end of the visual stimulus (7f, 320 cells, n = 3 mice; XCaMP-Gf, 124 cells, n = 3 mice; 8f, 317 cells, n = 5 mice; 8m, 365 cells, n = 3 mice; 8s, 655 cells, n = 6 mice). Kruskal-Wallis multiple-comparison test, *P* < 0.001. Dunn’s comparison test shown: \**P* < 0.05; \*\*\**P* < 0.001; ns, not significant. Full statistics in **Methods**. F. Proportion of cells responding to visual stimuli. 7f, 12 FOVs from n = 3 mice; 8f, 19 FOVs, n = 5 mice; 8m, 14 FOVs, n = 3 mice; 8s, 26 FOVs, n = 6 mice. Tukey’s multiple comparison test, *P* < 0.001. One-way ANOVA test shown: \**P* < 0.05; \*\*\**P* < 0.001; ns, not significant. Full statistics in **Methods**. G. Distribution of response amplitude (ΔF/_0_) for preferred stimulus. The 75th percentile ΔF/F_0_ (%) values for each construct: 71 (7f), 24 (8f), 59 (8m), 138 (8s). 7f, 1,053 cells from n = 3 mice; 8f, 1,253 cells, n = 5 mice; 8m, 848 cells, n = 3 mice; 8s, 1026 cells, n = 6 mice. Full statistics in **Methods**.

With the jGaMP8 indicators, visual stimulus-evoked fluorescence transients were observed in many cells (**Fig. 3B,C**; three representative cells shown for 8s and 8f) and were stable across trials (**Supp. Fig. 12**). All sensors produced transients with rapid rise and decay (**Fig. 3B-E**). Nearly identical responses were measured after long-term expression of jGCaMP8 (five additional weeks; **Supp. Fig. 13**). XCaMP-Gf was much dimmer (10x) than the others (**Supp. Fig. 14A-B**) with few responsive cells, precluding sensitivity analysis. As protein expression levels were similar across indicators (**Supp. Fig. 14C-D**), XCaMP-Gf is deficient in maturation or brightness *in vivo* and was not studied further.

The contrast changes in visual stimuli were tracked faithfully by sensor responses (**Fig. 3B,C**). Consistent with *in vitro* characterization, jGCaMP8f showed significantly shorter half-decay time (median[1^st^-3^rd^ quartiles] = 84[32-153] ms) than 7f (110[41-223] ms, *P* < 0.05) and comparable to 8m (84[32-165] ms) and XCaMP-Gf (91[48-155] ms) (**Fig. 3E**). On the other hand, 8s decay was significantly slower than the other indicators.

We quantified indicator sensitivity both as the proportion of labeled neurons responsive ^6, 8^ to visual stimuli (**Fig. 3F**) and as the cumulative distribution of peak ΔF/F_0_ across cells (**Fig. 3G**). Significantly more responsive cells were seen for 8s and 8m than for 8f and 7f (**Fig. 3F;** *P* < 0.001). Furthermore, 8s was dramatically right-shifted relative to the other indicators (**Fig. 3G**), reflective of its high sensitivity and saturating ΔF/F_0_. SNR of visually evoked fluorescence transients was significantly higher for 8s than for other sensors, followed by 8m and 7f, then by 8f (**Fig. 3G**).

Orientation tuning was similar for all sensors, except that 8m and 8s revealed a larger proportion of neurons with low orientation selectivity (**Supp. Fig. 15**). A plausible explanation for this is that the high-sensitivity indicators detect activity of GABAergic interneurons that is missed by the other sensors. (Interneurons yield smaller fluorescence responses ^8^, and have much less sharp tuning than excitatory neurons ^34^.) This hypothesis is supported by experiments with simultaneous imaging and electrophysiology (see below).

### Simultaneous imaging and electrophysiology

To quantify GECI responses to neural activity, we combined two-photon imaging (122 Hz) and loose-seal, cell-attached electrophysiological recordings (**Fig. 4A**). We compared fluorescence changes and spiking across sensors (*n*=40 cells from 8 mice, 8f; *n*=47 cells from 7 mice, 8m; *n*=49 cells from 7 mice, 8s; *n*=23 cells from 5 mice, 7f; **Supp. Fig. 16, Supp. Fig. 17, Supp. Table 5**). Fluorescent signals for cell body regions of interest (ROIs) were corrected for neuropil signal ^6, 8^ (**Supp. Fig. 18**). All jGCaMP8 variants produced large fluorescence transients even in response to single action potentials (APs) (**Fig. 4B-D**).

**Figure 4.**
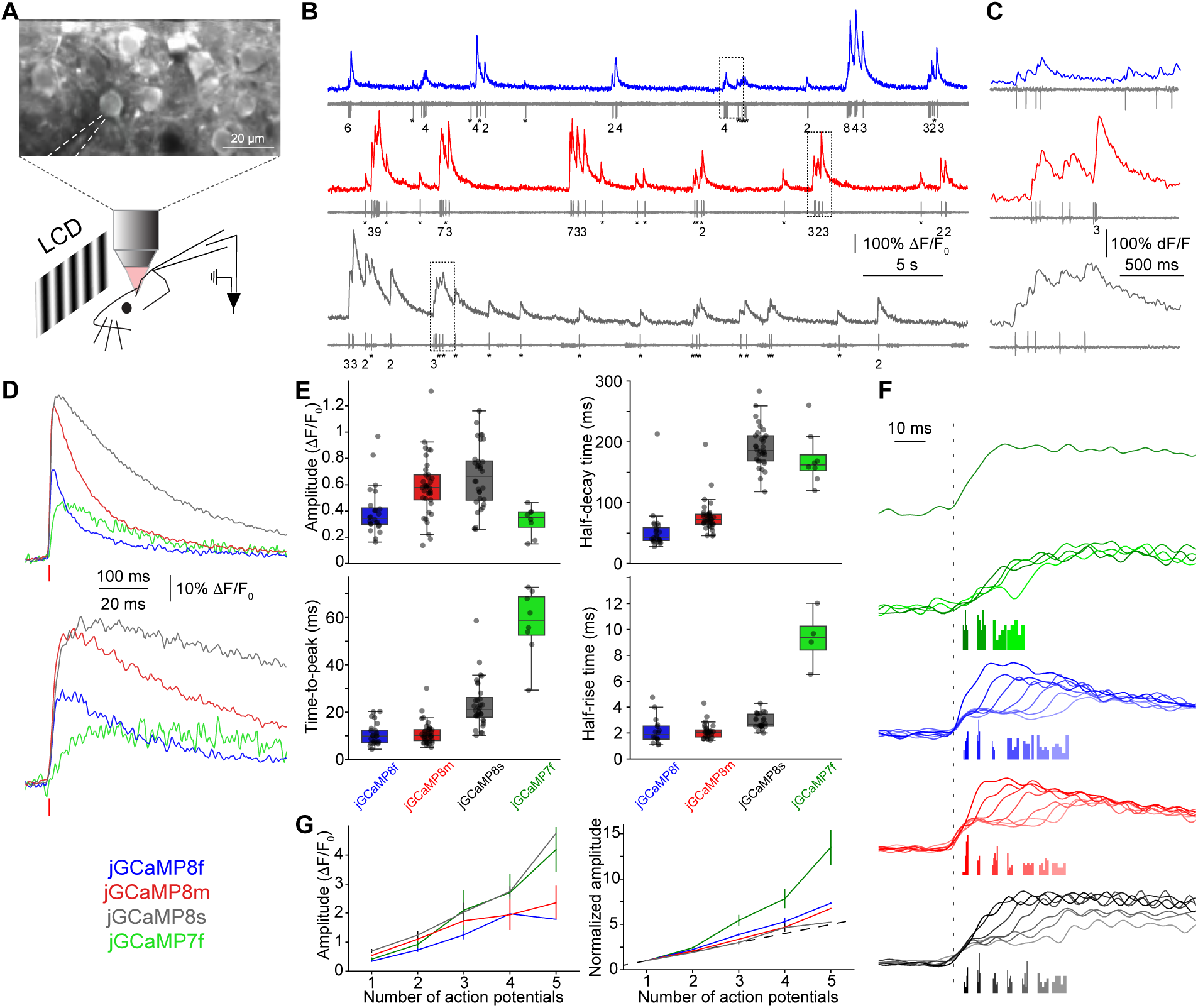
Simultaneous electrophysiology and imaging. A. Schematic of the experiment. Top, representative field of view. The recording pipette is indicated by dashed lines. B. Simultaneous fluorescence dynamics and spikes in example neurons expressing 8f (top), 8m (middle) and 8s (bottom). Number of spikes for each burst indicated below trace (single spikes, asterisks). C. Zoomed-in view of traces corresponding to dashed boxes in B. D. Grand average of fluorescence response elicited by single action potentials, aligned to action potential peak (red vertical bar), effectively sampled at 500 Hz (see text and **Supp. Fig. 19** for details). E. Fluorescence responses elicited by single action potentials. Box-whisker plot; black dots, single cells. F. Top, 7f response to a single action potential (from panel D). Bottom, response to action potential doublets, binned based on inter-spike intervals. Transients are aligned to the first action potential of the doublet (dotted line). The timing of the second action potential is denoted by the histograms below the transients. The inter-spike intervals are selected to be approximately 5, 10, 15, 20, 25, 30 and 35 ms. Responses for 7f, 8f, 8m, and 8s shown. G. Response linearity. Left, peak response as a function of number of action potentials within 20 ms window. Mean and s.e.m. shown. Right, same but normalized to 1-AP response.

Our experiments allowed us to resolve fluorescence transients with much higher effective temporal resolution than the nominal 122 Hz frame rate. Specifically, fields of view were chosen so that the patched neuron occupied <20% of the frame’s scan lines (**Supp. Fig. 19**). Since neurons were scanned at random phases with respect to AP fluorescence transients, neurons of interest were sampled with an effective temporal resolution of 500-600 Hz. All three jGCaMP8 variants showed rise (0 - 50%) times <5 ms, approximately 5x faster than 7f (**Fig. 4C-E**). Peak responses for all jGCaMP8 indicators were also larger than for 7f (**Fig. 4D-E**). To study spike-time estimation, we first binned action potential doublets with respect to their inter-spike interval length. The jGCaMP8 indicators resolved individual action potentials from doublets at spike rates of up to 100 Hz (**Fig. 4F**). We subsequently grouped spike bursts based on the number of APs (from 1 to 5) in a 20 ms integration window. All sensors show monotonic increases in fluorescence response with AP count, with the jGCaMP8 sensors responding more linearly than jGCaMP7f (**Fig. 4G**). This greater linearity is consistent with neuronal culture results and lower Hill coefficients in purified protein (**Supp. Table 6**).

The *synapsin-1* promoter yields expression in many neurons, including both pyramidal cells and fast-spiking (FS, presumably parvalbumin expressing) interneurons ^35^ – which are interspersed in our imaged ROIs. Out of our recorded neurons, we identified the subset of FS spiking interneurons by their high spike rates and short spike durations (**Supp. Fig. 20A**)^36^. All three jGCaMP8 sensors produced robust responses (**Supp. Fig. 20B;** ∼3% ΔF/F_0_ on average, with responses up to 5%) to single APs in FS interneurons, much larger than GCaMP6s (∼1% ΔF/F_0_)^8^.

Taken together, in mouse cortex *in vivo*, the jGCaMP8 sensors show excellent single-spike detection and spike train deconvolution, spike time estimation, good expression, strong performance in fast-spiking interneurons, and no evidence of adverse effects of long-term expression.

We also tested the jGCaMP8 variants alongside 6f and 7f in mouse cerebellar Purkinje cell dendritic arbors with the *Pcp2*-Cre mouse line (**Supp. Fig. 21A-B**). All jGCaMP8 variants showed faster rise-time than 6f and 7f (**Supp. Fig. 21C-D**), and decay-time was markedly faster for 8m and 8f (**Supp. Fig. 21E**).

### Spike train modeling with jGCaMP8

Calcium-evoked fluorescence is an indirect measure of neural activity ^8, 37^. A large body of work has been devoted to estimating spike trains from calcium imaging data. The performance of these deconvolution algorithms is largely limited by linearity, sensitivity and kinetics of the calcium-dependent sensors ^18, 38–40^. We tested the effects of the faster kinetics, superior linearity, and high SNR of the jGCaMP8 indicators on state-of-the-art spike-fitting models ^37^ (**Methods**), using our simultaneous imaging and electrophysiology data (**Fig. 4**; **Fig. 5A**). We compared the variance explained across linear and non-linear (“sigmoid”) models and measured to what extent nonlinearities are required to fit fluorescence dynamics for different indicators (**Fig. 5B**).

**Figure 5.**
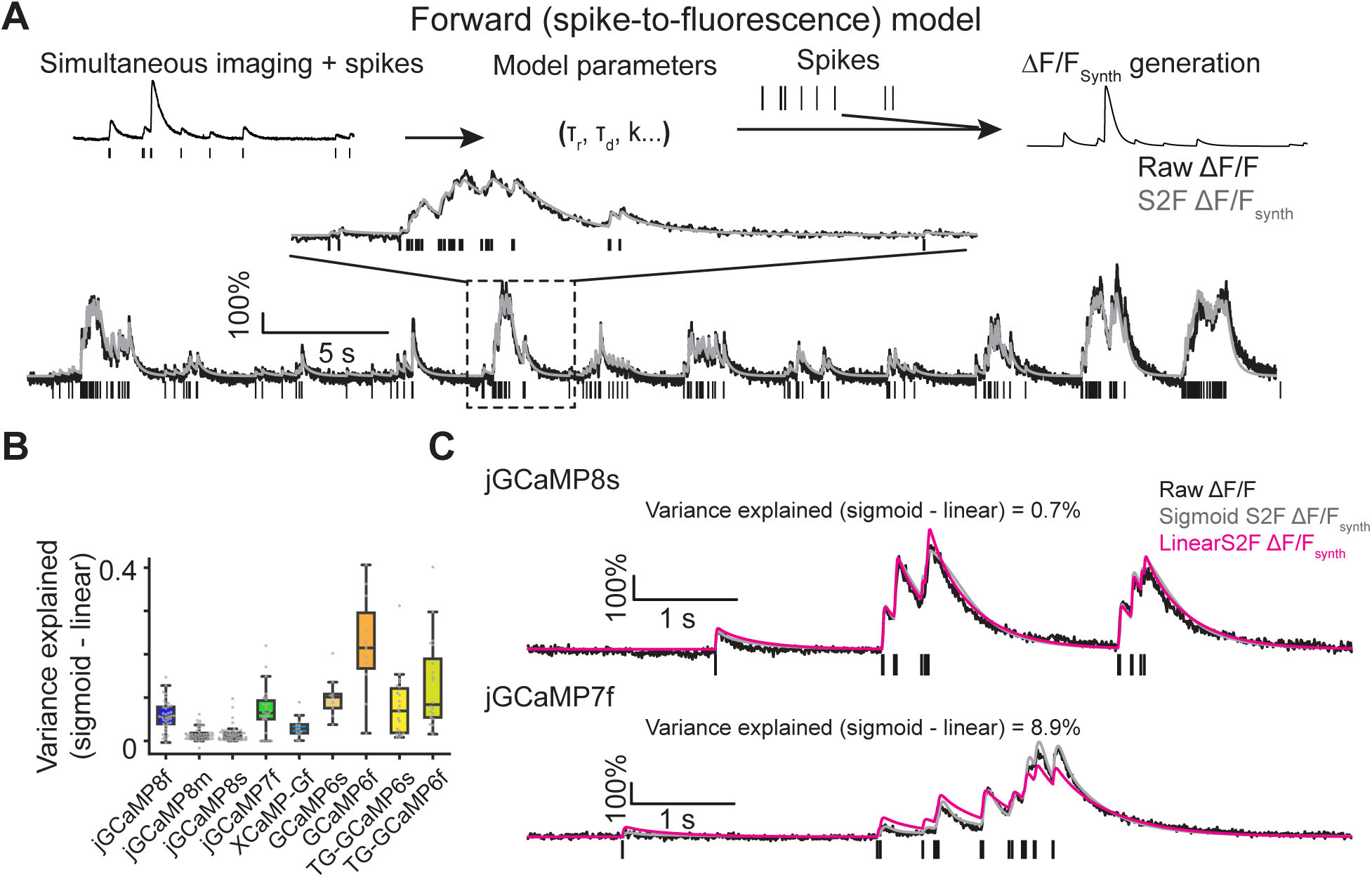
Spike fitting. A. Spike-to-fluorescence model. Top: schematic plot of the spike-to-fluorescence (S2F) forward model that generates a synthetic fluorescence trace (ΔF/F_Synth_) from an input spike train. Bottom: example fit and data of a cell. Experimental, measured ΔF/F_0_ (black) is overlaid with the simulated ΔF/F_Synth_ (gray) from the S2F model. The input to the model, the simultaneously recorded spikes (black), is shown below the traces. B. The degree of nonlinearity (measured by the difference of variance explained using a sigmoid fit from that using a linear fit). Nonlinearity is low in jGCaMP8 sensors (see **Supp. Table 6** for more details) but high in GCaMP6 sensors. C. Exemplary cell dynamics with different degrees of nonlinearities. Black lines, measured ΔF/ F_0_; gray lines, simulated ΔF/F_Synth_ from the S2F sigmoid model; magenta lines, simulated ΔF/ F_Synth_ from the S2F linear model.

Linear models performed much better for jGCaMP8 than GCaMP6 or jGCaMP7f in fitting fluorescence traces (8m and 8s had the best variance explained, at 85.0 ± 1.8% and 78.3 ± 2.8%, respectively; mean ± s.e.m.; **Supp. Table 7**), reflecting their improved linearity (**Fig. 5B-C; Supp. Table 7**), SNR, and kinetics (**Supp. Fig. 22A-G; Supp. Table 8**). Model estimates of rise- and decay-time constants are consistent with direct measurement (**Supp. Fig. 22C, F**). Moreover, the model showed that the jGCaMP8 indicators maintain linearity over a wide range of activity regimes, whereas jGCaMP7f showed both sub-linearity at low and supra-linearity at high spike rates (**Fig. 5C; Supp. Fig. 22H**). Consistent with their linearity and high sensitivity, 8m and 8s outperformed other indicators in recovering spike timing in widely used deconvolution algorithms ^38^ (**Supp. Fig. 22I).** Altogether, the linear model showed excellent performance at fitting activity profiles generated from the three jGCaMP8 indicators.

## Discussion

jGCaMP8s is the most sensitive GECI available, and jGCaMP8f is the fastest. jGCaMP8m shows intermediate sensitivity and speed. The rapid kinetics lead to high SNR for small transients and excellent deconvolution of spikes in trains.

The jGCaMP8 indicators feature a different calmodulin (CaM)-binding peptide (from endothelial nitric oxide synthase) than previous GECIs (mostly the RS20 peptide from myosin light-chain kinase). The XCaMP sensors as based on the ckkap peptide from CaM-dependent kinase kinase CaMKK-α/β. In our hands, the XCaMP sensors were quite dim in all preparations – even with high-level expression – precluding side-by-side comparison. ^43–45^

In addition to the fast kinetics and large fluorescence response, the jGCaMP8 indicators are more linear with respect to [Ca^2+^] and spike number. This linearity provides more interpretable data. The new sensors will provide better estimates of the number and timing of transients, both for action potentials as demonstrated here, and for synaptic inputs – which will provide a clearer picture of the size and relative timing of inputs into spines and dendrites in single neurons and populations.

Previous generations of “fast GCaMPs” ^19, 20^ suffered from low signal-to-noise, as ΔF/F_0_ was compromised in the pursuit of kinetics. The combination of large-scale peptide grafting and massive screening of linker and interface mutants was able to overcome the intrinsic barriers to simultaneously maintaining fast kinetics and high SNR. The jGCaMP8 indicators showed similarly good performance after expression for several months as previous GCaMPs. Long-term, high-level GCaMP expression can result in “cytomorbid” cells with altered trafficking ^7^ and even epileptiform activity in mice ^41^. The sensor GCaMP-X ^42^ incorporates a second calmodulin-binding peptide as a “protection motif,” which decreases calcium channel disruption and nuclear accumulation. A similar strategy could be employed with the new sensors, although it is possible that incorporation of this second peptide would disrupt their fast kinetics and high SNR.

The jGCaMP8 indicators have improved kinetics across neuron culture, flies, fish, and mice. Furthermore, 8m and 8f also performed remarkably well in fast-spiking (likely parvalbumin) cortical interneurons, with large SNR and rapid kinetics, despite the small free calcium increases per action potential in these neurons ^44^. The jGCaMP8 indicators will be the reagents of choice for most calcium imaging applications.

## Methods

All surgical and experimental procedures were conducted in accordance with protocols approved by the Institutional Animal Care and Use Committee and Institutional Biosafety Committees of Howard Hughes Medical Institute (HHMI) Janelia Research Campus, and of the corresponding committees at the other institutions.

### Sensor design

We surveyed the Protein Data Bank for unique structures of calmodulin (CaM) in complex with a single peptide. Twenty-nine peptides were sufficiently different from the RS20 peptide sequence used in previous GCaMPs to warrant testing (**Supp. Table 1**). The structures of these complexes were superimposed on the GCaMP2 structure (PDB 3EK4) in PyMOL, and amino acids were added or removed to bring all peptides to a length estimated to work well in the GCaMP topology. Synthetic DNA encoding each of the 29 peptides replaced the RS20 peptide in the bacterial expression vector pRSET-A-GCaMP6s. Of the initial sensors, 20/29 sensed calcium. All 20 had lower saturating ΔF/F_0_ than GCaMP6s, all but three had weaker Ca^2+^ affinity (apparent *K_d_*) than GCaMP6s, all but one had lower cooperativity (Hill coefficient, *n*), and many were dimmer (**Supp. Table 1**). Several sensor variants showed much faster Ca^2+^ decay kinetics, as determined by stopped-flow fluorescence on purified protein (**Supp. Table 1**). Based on fast kinetics, saturating ΔF/F_0_, apparent *K_d_*, Hill coefficient, and apparent brightness, we prioritized variants based on the peptides from endothelial nitric oxide synthase (PDB 1NIW; peptide “ENOSP”) and death-associated protein kinase 1 (1YR5; peptide “DAPKP”) for optimization (**Supp. Table 1**).

### Sensor optimization

These two sensor scaffolds were optimized in protein purified from *Escherichia coli* expression. Libraries were constructed to mutate the linker connecting the peptide to cpGFP (linker 1) ^26^ and screened for high signal change and retained fast kinetics. The linker connecting cpGFP and CaM (linker 2) was similarly mutated on top of variants from the optimization of linker 1. Out of 4000 ENOSP-based variants and 1600 DAPKP-based variants, 23 and 10 respectively had fast kinetics and high saturating ΔF/F_0_ in purified protein (**data not shown**).

Guided by the structure of jGCaMP8.410.80, we targeted 16 interface positions for site-saturation mutagenesis: 7 in ENOSP, 4 on cpGFP, and 5 on CaM (**Supp. Fig. 1B**). Sensor variants were tested in cultured neurons for higher sensitivity in detecting neural activity while maintaining fast kinetics. Several single mutations improved properties (**Supp. Table 2**), particularly residues near the ENOSP C-terminus and the cpGFP-CaM interface. Beneficial point mutations were combined in subsequent rounds of screening. Ten additional CaM positions (**Supp. Fig. 1B**) surrounding ENOSP were next subjected to site-saturation mutagenesis. Finally, mutations (**Supp. Fig. 1B**) from the FGCaMP sensor (developed using CaM and RS20-like peptide sequences from the fungus *Aspergillus niger* and the yeast *Komagataella pastoris*) ^43, 44^ were introduced to improve biorthogonality and/or kinetics.

### Sensor screen and characterization in solution

Cloning, expression, and purification of sensor variants in *Escherichia coli*, calcium titrations, pH titrations, kinetic assay, and photophysical analysis were performed essentially as described before ^26^.

In this study, the RSET tag (His_6_ tag-Xpress epitope-enterokinase cleavage site), which had been carried over from the pRSET-A cloning vector in earlier work, was removed from all sensors: constructs simply encode a hexa-histidine affinity tag: Met-His_6_ tag-peptide-linker 1- cpGFP-linker 2-CaM. For the screen of linkers replacing RS20 (previously mistakenly referred to as “M13”), libraries of sensors in the pRSET-A bacterial expression vector were generated using primers containing degenerate codons (NNS) with Q5 site-directed mutagenesis (New England BioLabs) and transformed into T7 Express competent cells (New England BioLabs). A sequence encoding six repeats of the Gly-Gly-Ser tripeptide was designed as a highly flexible, presumably non-CaM-binding negative control. We expressed the new variants, as well as the presumptive Gly-Gly-Ser negative control and GCaMP6s as a positive control, in *Escherichia coli* T7 Express. Single colonies were picked and grown in 800 µL ZYM-5052 autoinduction medium containing 100 µg/mL ampicillin in 96 deep-well blocks for 48 hours at 30°C. Cells were collected by centrifugation, frozen, thawed, and lysed. Clarified lysate was used to estimate the dynamic range by measuring fluorescence in the presence of 1 mM Ca^2+^ or 1 mM EGTA.

For protein purification, T7 Express cells containing sensors were grown at 30°C for 48 hours in ZYM-5052 autoinduction medium with 100 µg/mL ampicillin. Collected cells were lysed in 1/50 volume of B-PER (Thermo Fisher) with 1 mg/mL lysozyme and 20 U/mL Pierce Universal Nuclease (Thermo Fisher) and subsequently centrifuged. Supernatants were applied to HisPur Cobalt Resin (Thermo Fisher). The resin was washed with 20 column volumes of 20 mM Tris, pH 8.0, 300 mM NaCl, 1 mM imidazole, followed by 10 column volumes of 20 mM Tris, pH 8.0, 500 mM NaCl, 5 mM imidazole. Proteins were eluted into 20 mM Tris, pH 8.0, 100 mM NaCl, 100 mM imidazole.

For calcium titrations, sensors were diluted 1:100 in duplicate into 30 mM MOPS, pH 7.2, 100 mM KCl containing either 10 mM CaEGTA (39 µM free calcium) or 10 mM EGTA (0 µM free calcium). As before, these two solutions were mixed in different amounts to give 11 different free calcium concentrations. GCaMP fluorescence (485 nm excitation, 5 nm bandpass; 510 nm emission, 5 nm bandpass) was measured in a Tecan Safire2 plate reader (Tecan). The data was fit with a sigmoidal function using KaleidaGraph (Synergy Software) to extract the apparent *K_d_* for Ca^2+^, the Hill coefficient, and dynamic range.

*k_off_* was determined at room temperature using a stopped-flow device coupled to a fluorimeter (Applied Photophysics). Each sensor variant in 1 µM Ca^2+^ in 30 mM MOPS, pH 7.2, 100 mM KCl was rapidly mixed with 10 mM EGTA in 30 mM MOPS, pH 7.2, 100 mM KCl. Fluorescence decay data was fit with a single or double exponential decay function.

For pH titrations, purified proteins were diluted into pH buffers containing 50 mM citrate, 50 mM Tris, 50 mM glycine, 100 mM NaCl and either 2 mM CaCl_2_ or 2 mM EGTA, which were pre-adjusted to 24 different pH values between 4.5 and 10.5 with NaOH. A sigmoidal function was used to fit fluorescence versus pH, and the *pK_a_* value was determined from the midpoint.

### Sequence & structural analysis of variants

Linker1 encodes Leu-Glu in GCaMP6s (and indeed, in all previous RS20-based GCaMP sensors – this linker was extensively mutated in the GCaMP5 screen ^26^, but the best variant, GCaMP5G, retained Leu-Glu); we first mutated Leu-Glu to fully degenerate 2-amino acid (aa) sequences and screened for variants with both high signal change and retained fast kinetics. Following selection of the best 2-aa linkers, these variants were expanded to libraries of 3-aa linkers by addition of fully degenerate residues. After optimization of Linker 1, Linker 2 was mutated from Leu-Pro, to which it had been selected in GCaMP5G ^26^, the parent of GCaMP6 and GCaMP7. Linker 2 mutagenesis was similar to that for Linker 1, but alternative Linker 2 sequences either slowed kinetics or decreased ΔF/F_0_, and Linker 2 was thus retained as Leu-Pro.

In addition to 8f/m/s, several other variants may be of interest, including 455, 543, 640, 707, and 712 (**Supp. Table 2**). All promising variants contain, in addition to the Leu-Lys-Ile linker 1, additional mutations to the ENOSP peptide: Asn19Thr and Ser24Ile appear in every variant except 712, Ser26Arg appears in every variant but jGCaMP8s (with Ser26Met), jGCaMP8m has Ala25Gly, and 712 has Met28Ser. Every variant contains the Gln88Glu mutation at the CaM-GFP interface. Further mutations include Phe286Tyr (8s, 8m, and 707); Glu288Gln (707); Gln315Leu (8f), Gln315His (8s, 707), Gln315Lys (455); Met346Gln (543); and Met419Ser (640). Of these, Phe286Tyr comes from the FGCaMP sensor; all others are unique to this work. Importantly, GCaMP6s data from both purified protein and cultured neurons are essentially identical between this work (lacking the RSET tag) and previous work (with it) (**data not shown**) – implying that the RSET tag does not noticeably modulate GCaMP function in protein and neuronal culture and that observed jGCaMP8 improvements stem from the peptide substitution and other mutations.

### Photophysical measurements

All the measurements were performed in 39 *μ*M free Calcium(+Ca) buffer (30 mM MOPS, 10 mM CaEGTA in 100 mM KCl, pH 7.2) or 0 *μ*M free Calcium(-Ca) buffer (30 mM MOPS, 10 mM EGTA in 100 mM KCl, pH 7.2). Absorbance measurements were performed using UV-Vis spectrometer (Cary 100, Agilent technologies), and fluorescence excitation-emission spectra were measured using a fluorimeter (Cary Eclipse, Varian Inc.). ΔF/F_0_ between proteins in +Ca and -Ca buffer was calculated from the fluorescence emission spectra. Quantum yield measurements were performed using an integrating sphere spectrometer (Quantaurus, Hamamatsu) for proteins in +Ca buffer. Extinction coefficients were determined *via* the alkali denaturation method, using extinction coefficient of denatured GFP as a reference (ε = 44000 M^-1^cm^-1^ at 447 nm).

### Two-photon spectroscopy

The two-photon excitation spectra were performed as previously described ^26^. Protein solutions of 1-5 *μ*M concentration in +Ca or -Ca buffer were prepared and measured using an inverted microscope (IX81, Olympus) equipped with a 60X, 1.2NA water immersion objective (Olympus). Two-photon excitation was obtained using an 80 MHz Ti-Sapphire laser (Chameleon Ultra II, Coherent) with sufficient power from 710 nm to 1080 nm. Fluorescence collected by the objective was passed through a short-pass filter (720SP, Semrock) and a bandpass filter (550BP200, Semrock) and detected by a fiber-coupled Avalanche Photodiode (APD) (SPCM_AQRH-14, Perkin Elmer). The obtained two-photon excitation spectra were normalized to 1 *μ*M concentration and subsequently used to obtain the action cross-section spectra (AXS) with fluorescein as a reference (Average AXS from ^45, 46^).

Fluorescence correlation spectroscopy (FCS) was used to obtain the 2P molecular brightness of the protein molecule. The peak molecular brightness was defined by the rate of fluorescence obtained per total number of emitting molecules. 50-100 nM protein solutions were prepared in +Ca buffer and excited with 930 nm wavelength at various power ranging from 2-30 mW for 200 sec. Emission fluorescence was collected by an APD and fed to an autocorrelator (Flex03LQ, Correlator.com). The obtained autocorrelation curve was fit to a diffusion model ^47^ to determine the number of molecules <N> present in the focal volume. The 2-photon molecular brightness *(ε)* at each laser power was calculated as the average rate of fluorescence *<F>* per emitting molecule *<N>*, defined as *ε* = *<F>/<N>* in kilocounts per second per molecule (kcpsm). As a function of laser power, the molecular brightness initially increases with increasing laser power, then levels off and decreases due to photobleaching or saturation of the protein chromophore in the excitation volume. The maximum or peak brightness achieved, *<ε_max_>*, represents a proxy for the photostability of a fluorophore.

### Screening in neuronal cell culture

GCaMP variants were cloned into an *hSyn1*-GCaMP-NLS-mCherry-WPRE expression vector, and XCaMP variants (XCaMP-G, XCaMP-Gf, XCaMP-Gf0) were cloned into an AAV-*hSyn1*-XCaMP-NES vector. We used the nuclear export sequence (NES) for the XCaMP sensors as this was how they were characterized in the original publication^23^. As this excludes the XCaMP sensors from the nucleus, where Ca^2+^ signals are slower ^48^, whereas the variants developed here were not explicitly excluded (although GCaMPs without an explicit NES are nevertheless fairly nuclear-excluded), this will make the XCaMPs appear faster than they really are compared to the GCaMP indicators.

The primary rat culture procedure was performed as described^6^. Briefly, neonatal rat pups (Charles River Laboratory) were euthanized, and neocortices were dissociated and processed to form a cell pellet. Cells were resuspended and transfected by combining 5×10^5^ viable cells with 400 ng plasmid DNA and nucleofection solution in a 25 µL electroporation cuvette (Lonza). Electroporation of GCaMP mutants was performed according to the manufacturer’s protocol.

Neurons were plated onto poly-D-lysine (PDL) coated, 96-well, glass-bottom plates (MatTek) at ∼1×10^5^ cells per well in 100 µL of a 1:2 mixture of NbActiv4 (BrainBits) and plating medium (28 mM glucose, 2.4 mM NaHCO_3_, 100 µg/mL transferrin, 25 µg/mL insulin, 2 mM L-glutamine, 100 U/mL penicillin, 10 µg/mL streptomycin, 10% FBS in MEM). Typically, each plate included GCaMP6s (8 wells), GCaMP6f (8 wells), and jGCaMP7f (8 wells). Other wells were electroporated with mutated variants (4 wells per variant), for a total of 80 wells (the first and last columns in the plate were not used). Plates were left in the incubator at 37°C and 5% CO_2_.

On DIV 14-19, neurons underwent field stimulation and imaging^6, 49^. Fluorescence time-lapse images (200 Hz; total of 7 seconds) were collected on an Olympus IX81 microscope using a 10x, 0.4 NA objective (UPlanSApo, Olympus) and an ET-GFP filter cube (Chroma #49002). A 470 nm LED (Cairn Research) was used for excitation (intensity at the image plane, 0.34 mW/ mm^2^). Images were collected using an EMCCD camera (Ixon Ultra DU897, Andor) with 4×4 binning, corresponding to a 0.8 mm x 0.8 mm FOV. Reference images (100 ms exposure) were used to perform segmentation. Red illumination for variants co-expressing mCherry was performed with a 590 nm LED (Cairn Research) through an ET-mCherry filter cube (Chroma #49008) with an intensity of 0.03 mW/mm^2^. Trains of 1, 3, 40, and 160 field stimuli were delivered with a custom stimulation electrode. For sensor linearity measurements, 1, 2, 3, 5, 10, and 40 field stimuli were delivered. For frequency response experiments, two-photon imaging was performed (objective: 16x, 0.8 numerical aperture (Nikon), wavelength 940 nm, laser power: 56 mW, acquisition rate: 155 Hz; filter set: 525/50 nm (for signal collection) and a 565 nm dichroic mirror).

The responses of individual variants were analyzed as described ^6, 8^. The Ilastik toolkit ^50^ was used to segment cell bodies in the reference images. Wells with fewer than five detected neurons, and wells with poor neuronal proliferation, were discarded. Plates with more than four discarded control (GCaMP6s) wells were discarded and re-screened. The ΔF/F_0_, SNR, and kinetics (half-rise, half-decay, time-to-peak) metrics were computed for each cell. Median values from each well are reported to quantify performance. Each observation was normalized to the median GCaMP6s value from the same experimental batch. Baseline brightness for constructs co-expressing mCherry (in a [Binder notebook](https://mybinder.org/v2/gh/ilyakolb/interactive-multiparam-plot/master?filepath=interactive-multiparameter-screening-plot.ipynb) and **Supp. Table 2**) was calculated by dividing the GFP cellular fluorescence in the beginning of the 3-AP stimulation epoch by the mCherry cellular fluorescence (for a ratiometric measurement). For comparison with XCaMP variants (in **Supplementary Figure 4**), no mCherry normalization was performed, but all baseline brightness values were still normalized to GCaMP6s in the same transfection week. To determine significant differences in observations between constructs, a two-tailed Mann-Whitney U-test was performed between constructs and controls (GCaMP6s or jGCaMP7f). A median ΔF/F_0_ trace was computed across all detected cell bodies in a well for each stimulus. Photobleaching was corrected in the 1-AP recordings by fitting a double exponential to the beginning and end segments of the fluorescence trace.

Finally, variants were filtered according to four criteria to remove noisy, non-responding clones. 1) Variants with half-rise time > 4x slower than GCaMP6s, as these represented poor fits or noise. 2) Similarly, variants with time-to-peak >3x that of 6s. 3) Variants with half-rise time < 0.1x that of 6s, as these represented sensors with poor ΔF/F_0_. 4) Similarly, variants with half-decay time <0.01x that of 6s. This 683 (out of 776 tested) jGCaMP8 variants, along with the controls, that showed detectable response to 1 AP in cultured neurons (**Supp. Fig. 2**). All the parameters measured in our screen can be examined as an interactive scatterplot in a [Binder notebook](https://mybinder.org/v2/gh/ilyakolb/interactive-multiparam-plot/master?filepath=interactive-multiparameter-screening-plot.ipynb).

### Baseline fluorescence measurements

In a separate round of measurements from those measuring ΔF/F_0_, SNR, and kinetics, the baseline fluorescence of jGCaMP8 series was compared to jGCaMP7f and the XCaMP series. Due to significant week-to-week variability in baseline fluorescence, all constructs for this experiment were transfected side-by-side (2 consecutive transfection weeks, five 96-well plates). To minimize possible plate-to-plate variability within each transfected batch, the baseline fluorescence of each construct was normalized to in-plate GCaMP6s.

### Fluorescence recovery after photobleaching

FRAP experiments were carried out on a Nikon Ti-E inverted microscope outfitted with a Yokogowa CSU-X1 spinning disk and an Andor DU-897 EMCCD camera. Fluorescence excitation was carried out using a solid-state laser line at 488nm, and emission was collected with 100x 1.49NA objective (Nikon Instruments) through a standard GFP filter set. Photobleaching was performed using a Bruker Mini-Scanner by focusing a 405 nm laser to a single, diffraction-limited spot for 100 ms. Cultured neurons plated in 35 mm glass-bottom dishes (MatTek) were immersed in regular imaging buffer with the addition of synaptic blockers (same as used for neuronal culture field stimulation) and 1 µM TTX to block AP generation. In a subset of experiments, the buffer was supplemented with 5 µM ionomycin. Bleaching spots were chosen to be on the soma of the neuron but distant from the nucleus. A spot was photobleached 10 times (0.1 Hz) as the cell was concurrently imaged at 25 or 50 frames per second.

For analysis, pixels within a 1.5 µm radius around the bleach spot were averaged in each frame. The resulting fluorescence trace was normalized to the mean fluorescence of an identically sized spot on the opposite side of the soma, outside the nucleus. The trace was then split into 10 epochs (each corresponding to a bleaching event) and the fluorescence *f_i_*(*t*) of each epoch was normalized by dividing by the fluorescence value immediately preceding the bleaching pulse (*f_i_*_(_*t_pre_*_)_) as follows:

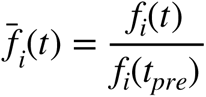

The resistant fraction was calculated as follows:

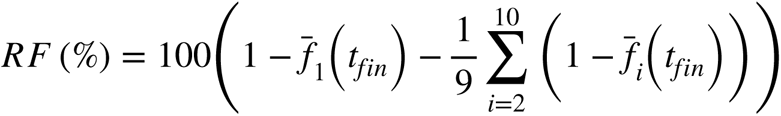

where 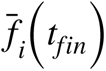 is the final fluorescence value at the end of epoch *i*, and the final term in the equation is the averaged fluorescence loss of all epochs after the first. This term is subtracted to account for the overall fluorescence loss with each bleaching pulse.

### Crystal structure determination

All GCaMP samples for crystallization were kept in 20 mM Tris, 150 mM NaCl, pH 8.0, 2 mM CaCl_2_. All crystallization trials were carried out at 22°C with the hanging-drop vapor diffusion method. Commercial sparse-matrix screening solutions (Hampton Research) were used in initial screens. 1 µL of protein solution was mixed with 1 µL of reservoir solution and equilibrated against 300 µL of reservoir solution. Diffraction data were collected at beamline 8.2.1 at the Berkeley Center for Structural Biology and processed with XDS ^51^. The phase was determined by molecular replacement using MOLREP , and the structure of GCaMP2 (PDB 3EK4) ^52, 53^ without the RS20 peptide as the starting model. Refinement was performed using REFMAC ^54^, followed by manual remodeling with Coot ^55^. Details of the crystallographic analysis and statistics are presented in **Supp. Table 3**.

### Adult Drosophila L2 Assay

GECIs were tested by crossing males carrying the variant to a w+ ; *53G02*-Gal4^AD^ (in attP40); *29G11*-Gal4^DBD^ (in attP2) females ^56^. Heterozygous flies were used in our experiments. Flies were raised at 21°C on standard cornmeal molasses media.

Females 3-5 days after eclosure were anesthetized on ice. After transferring to a thermoelectric plate (4°C), legs were removed, and then facing down, the head was glued into a custom-made pyramid using UV-cured glue. The proboscis was pressed in and fixed using UV-cured glue. After adding saline (103 mM NaCl, 3 mM KCl, 1 mM NaH_2_PO_4_, 5 mM TES, 26 mM NaHCO_3_, 4 mM MgCl_2_, 2.5 mM CaCl_2_, 10 mM trehalose and 10 mM glucose, pH 7.4, 270–275 mOsm) to the posterior side of the head, the cuticle was cut away above the right side, creating a window above the target neurons. Tracheae and fat were removed, and muscles M1 and M6 were cut to minimize head movement.

Two-photon imaging took place under a 40x 0.8NA water-immersion objective (Olympus) on a laser-scanning microscope (BrukerNano, Middleton, WI) with GaAsP photomultiplier tubes (PMTs). Laser power at 920 nm was kept constant at 8 mW using a Pockels cell. No bleaching was evident at this laser intensity. The emission dichroic was 580 nm and emission filters 511/20-25 nm. Images were 32×128 pixels with a frame rate at 372 Hz.

A MATLAB script drove the visual stimulation *via* a digital micromirror device (DMD, LightCrafter) at 0.125 Hz onto a screen covering the visual field in front of the right eye. A blue LED (Thorlabs M470L3) emitting through a 474/23-25 nm bandpass filter (to keep blue light from contaminating the green imaging channel) provided illumination.

Light dimming produced a stereotypical calcium increase in L2 neurons^28–30^. Intensity measurements were taken in medulla layer 2 (**Fig. 2A**; modified from ^57^). A target region image was chosen by testing each focal layer with 0.5 Hz full-field visual stimulation until a layer with maximum ΔF/F_0_ was identified. Then 2-3 columns producing a maximum response were identified within this layer. In addition to the ROI containing these L2 columns, a background ROI was selected where no fluorescence was evident. The mean background intensity was subtracted from the mean L2 ROI. Imaging then targeted this region over a protocol involving multiple tests, as such:

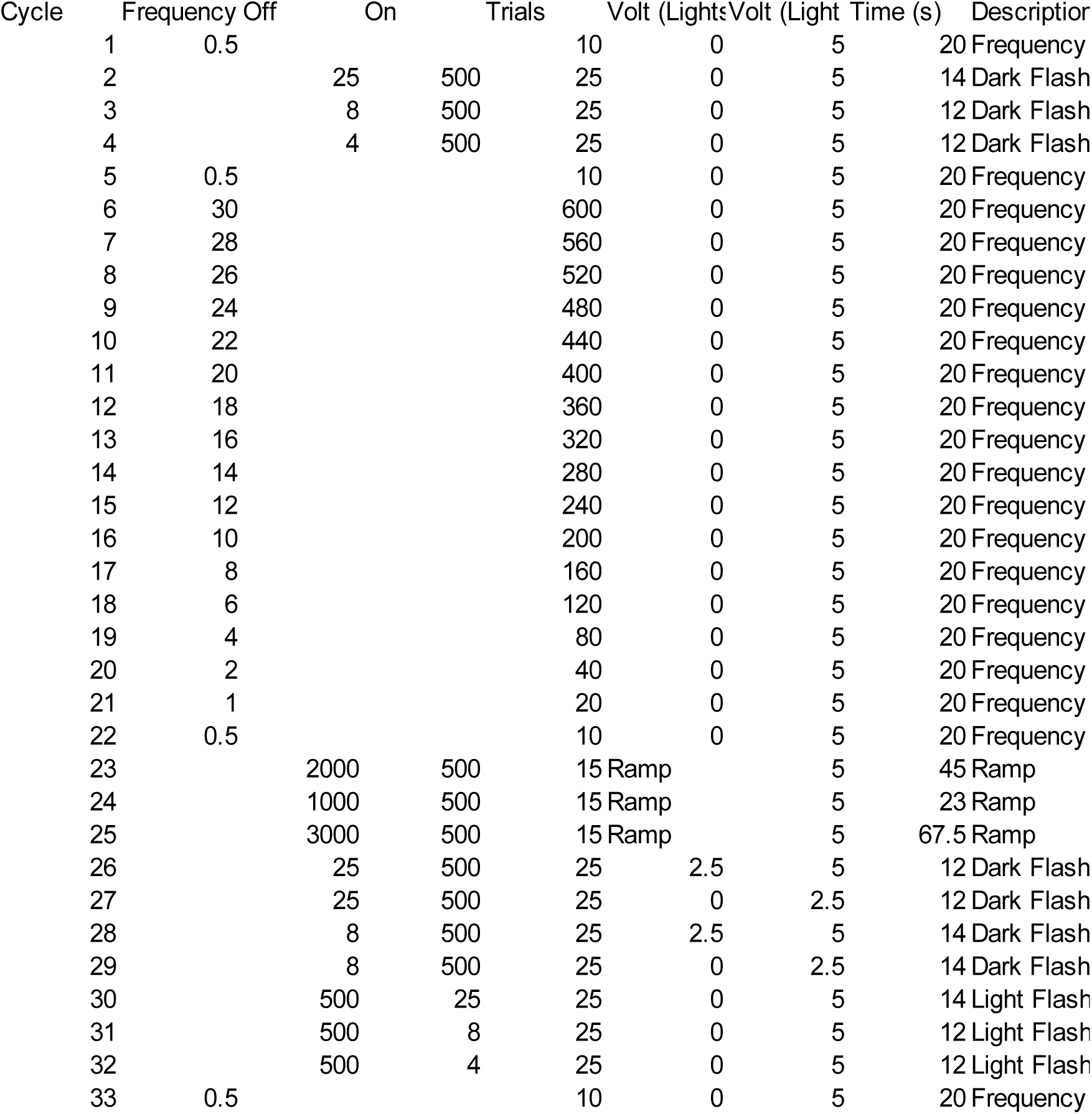

Image analysis was performed using custom Python scripts. In the ΔF/F_0_ calculation, baseline F_0_ included the last 20% of images taken at the end of the light period. Stimulus onset is the light-to-dark transition. Change in fluorescence ΔF is the intensity minus baseline. ΔF/F_0_ is ΔF divided by baseline. The final signal is processed through a Gaussian filter (σ=3).

### Imaging in the *Drosophila* larval neuromuscular junction

We made 20XUAS-IVS-Syn21-op1-GECI-p10 in VK00005 transgenic flies ^58^ and crossed them with 10XUAS-IVS-myr::tdTomato in su(Hw)attP8 x *R57C10*-Gal4 at VK00020; *R57C10*-Gal4 at VK00040 double-insertion pan-neuronal driver line. Heterozygous flies were used in our experiments. Sensor cDNAs were codon-optimized for *Drosophila*. The NMJ assay is as in our previous study^1^. Briefly, female 3^rd^ instar larvae were dissected in chilled (4°C) Schneider’s Insect Medium (Sigma) to fully expose the body wall muscles. Segment nerves were severed in proximity to the ventral nerve cord (VNC). Dissection medium was then replaced with room temperature HL-6 saline in which 2 mM CaCl_2_ and 7 mM of L-glutamate were added to induce tetany – freezing the muscles in place. A mercury lamp (X-CITE exacte) light source was used for excitation, and out-of-objective power was kept less than 5 mW to reduce bleaching. Type Ib boutons on muscle 13 from segment A3-A5 were imaged while the corresponding hemi-segment nerve was stimulated with square voltage pulses (4 V, 0.3 ms pulse width, 2 s duration, 1-160 Hz frequency) through a suction electrode driven by a customized stimulator. Bath temperature and pH were continuously monitored with a thermometer and pH meter, respectively, and recorded throughout the experiment. The filters for imaging were as follows: excitation: 472/30 nm; dichroic: 495 nm; emission: 520/35 nm. Images were captured with an EMCCD (Andor iXon 897) at 128.5 frames per second and acquired with Metamorph software. ROIs around boutons were manually drawn, and data were analyzed with a custom Python script.

#### NMJ immunofluorescence

Variants were crossed to a pan-neuronal driver line, also containing tdTomato, (pJFRC22-10XUAS-IVS-myr::tdTomato in su(Hw)attP8 ;; *R57C10* at VK00020, *R57C10* at VK00040). 3^rd^ instar larvae were filleted and fixed following standard techniques ^59^. Primary chicken anti-GFP (Thermo Fisher A10262, 1:1000) and secondary goat anti-chicken AlexaFluor 488 plus (Thermo Fisher A32931, 1:800) were used to stain GECIs. Primary rabbit anti-RFP (Clontech 632496, 1:1000) and secondary goat anti-rabbit Cy3 (Jackson 111-165-144, 1:1000) labeled tdTomato.

#### MBON-γ2α’1 immunofluorescence

Variants were co-expressed with membrane-localized myr::tdTomato using the *MB077B* driver. Adults 3-6 days old were harvested, brains dissected, and fixed using standard techniques. GCaMP variants were directly labeled with anti-GFP (AlexaFluor 488, Molecular Probes A-21311, 1:500). Primary Rat anti-RFP (mAb 5F8 Chromotek, 1:500) and secondary goat anti-rat Cy3 (Jackson 112-165-167, 1:1000) labeled tdTomato.

#### Immunofluorescence quantification

ROIs were drawn on targeted regions using custom Python scripts. Within each ROI, otsu-thresholding was used to identify regions expressing myr::tdTomato. Intensity measurements were then taken for both the variant and tdTomato within these regions. The ratio is the intensity from the green channel (variant staining) divided by the intensity from the red channel (myr::tdTomato staining).

#### Western blot

Protein was extracted from female brains with the same genotype used in the NMJ immunostaining. Western blots were performed following standard techniques. Each variant was stained using primary rabbit anti-GFP (Millipore Sigma) and secondary goat anti-rabbit IgG conjugated to horseradish peroxidase (HRP; Thermo Fisher). Actin was stained using mouse IgM anti-α-actin (Thermo Fisher, 1:5000) and goat anti-mouse IgG and IgM-HRP (Thermo Fisher, 1:5000). Signal was formed using SuperSignal West Dura luminescence and was imaged on a BioRad Gel imager. Band intensity was measured using FIJI. Band intensity from the variant was divided by band intensity from the actin band to determine the ratio.

### Light sheet imaging of larval zebrafish

#### Presentation of flashes during fictive swimming

Zebrafish larvae of 6-8 dpf expressing 8f, 7f and 6f as nuclearly-localized fusions to histone H2B under the pan-neuronal *elavl3* promoter were reared in conditions described in ^60^ and prepared for fictive behavior recording and light sheet imaging according to previously detailed methods ^32^. A LED torch, placed diagonally from the fish, emitted short pulses of light (100 ms) at 20- second intervals (**Supp. Fig. 11A).** The fish experienced no structured visual stimulus besides the constant 488 nm light sheet at all other times. To image the optic tectum, where visual responses were observed, we captured five horizontal planes at an interval of 20 µm spanning its dorsoventral axis in a single volume. We imaged at 30 Hz, a rate sufficient to capture the fast kinetics of the calcium sensors.

#### Identification of flash-responsive neurons

Fluorescence images were motion-corrected and segmented into cell segments by a custom preprocessing pipeline (https://github.com/zqwei/fish_processing).

To find flash-responsive neurons, we first computed a trial- and neuron-averaged response triggered on the onset of the flash stimulus for each fish for trials where the fish did not swim around the presentation of the stimulus. We then fit each of those responses to a difference between two exponentials to derive two time constants

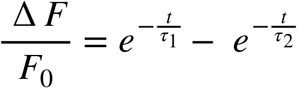

We then used the result of those fits to generate a calcium kernel with which regressors for the visual stimulus and motor output were generated. We found visual-responsive cells by looking for cells with a high visual coefficient (>75^th^ percentile) and a low motor coefficient (<25^th^ percentile). For each fish, we then computed another trial-averaged response triggered on the onset of the flash stimulus, this time only for visual-responsive cells and for all trials. These traces were averaged across fish to generate a variant-mean.

The variant-means were again fit to the function above to generate a calcium kernel for each variant. Finally, we once again performed regression with these calcium kernels to derive our final set of visual-responsive cells for each fish. Further analyses, such as the re-computation of the variant-mean **(Supp. Fig. 11C**), as well as single-cell characterization of response amplitude (**Supp. Fig. 11D**), half-rise time (**Supp. Fig. 11E**) and half-decay time (**Suppl. Fig. 11F**) were performed on this set of cells.

### Cortex: mouse surgeries

Young adult (postnatal day 50-214) male C57BL/6J (Jackson Labs) mice were anesthetized using isoflurane (2.5% for induction, 1.5% during surgery). A circular craniotomy (3 mm diameter) was made above V1 (centered 2.5 mm left and 0.5 mm anterior to the Lambda suture). Viral suspension (30 nL) was injected in 4-5 locations on a 500 µm grid, 300-400 µm deep. Constructs included: AAV2/1-*hSynapsin-1-*jGCaMP8 constructs (pGP-AAV-*syn1*-jGCaMP8f-WPRE, Addgene plasmid #162376, 4e12 GC/mL titer; pGP-AAV-*syn1*-jGCaMP8m-WPRE, Addgene plasmid #162375, 2.2e12 GC/mL titer; pGP-AAV-*syn1*-jGCaMP8s-WPRE, Addgene plasmid #162374, 2.1e12 GC/mL titer). A 3 mm diameter circular coverslip glued to a donut-shaped 3.5 mm diameter coverslip (no. 1 thickness, Warner Instruments) was cemented to the craniotomy using black dental cement (Contemporary Ortho-Jet). A custom titanium head post was cemented to the skull. An additional surgery was performed for loose-seal recordings. 18-80 days after the virus injection, the mouse was anesthetized with a mixture of ketamine-xylazine (0.1 mg ketamine & 0.008 mg xylazine per gram body weight), and we surgically removed the cranial window and performed durotomy ^61^. The craniotomy was filled with 10-15 µL of 1.5% agarose, then a D-shaped coverslip was secured on top to suppress brain motion and leave access to the brain on the lateral side of the craniotomy.

### Cortex: two-photon population imaging

Mice were kept on a warm blanket (37°C) and anesthetized using 0.5% isoflurane and sedated with chlorprothixene (20–30 µL at 0.33 mg/mL, intramuscular). Imaging was performed with a custom-built two-photon microscope with a resonant scanner. The light source was an Insight femtosecond-pulse laser (Spectra-Physics) running at 940 nm. The objective was a 16× water immersion lens with 0.8 numerical aperture (Nikon). The detection path consisted of a custom filter set (525/50 nm (functional channel), 600/60 nm (cell targeting channel) and a 565 nm dichroic mirror) ending in a pair of GaAsP photomultiplier tubes (Hamamatsu). Images were acquired using ScanImage (vidriotechnologies.com)^62^. Functional images (512 × 512 pixels, 215 × 215 µm^2^; or 512 × 128 pixels, 215 × 55 µm^2^) of L2/3 cells (50–250 µm under the pia mater) were collected at 30 Hz or 122 Hz. Laser power was up to 50 mW at the front aperture of the objective unless stated otherwise for the XCaMP-Gf experiments.

### Cortex: loose-seal recordings

Micropipettes (3–9 MΩ) were filled with sterile saline containing 20 µM AlexaFluor 594. Somatic cell attached recordings were obtained from upper layer 2 neurons (50-200 µm depth from brain surface) visualized with the shadow patching technique ^63^. Spikes were recorded either in current clamp or voltage clamp mode. Signals were filtered at 20 kHz (Multiclamp 700B, Axon Instruments) and digitized at 50 kHz using Wavesurfer *(wavesurfer.janelia.org/)*. The frame trigger pulses of ScanImage were also recorded and used offline to synchronize individual frames to electrophysiological recordings. After establishment of a low-resistance seal (15–50 MOhm), randomized visual stimulation was delivered to increase the activity of the cells in the field of view. In a small subset of recordings, we microstimulated the recorded neuron in voltage-clamp recording mode by applying DC current to increase its firing probability ^64^.

### Cortex: visual stimulation

Visual stimuli were moving gratings generated using the Psychophysics Toolbox in MATLAB (Mathworks), presented using an LCD monitor (30 × 40 cm^2^), placed 25 cm in front of the center of the right eye of the mouse. Each stimulus trial consisted of a 2 s blank period (uniform gray display at mean luminance) followed by a 2 s drifting sinusoidal grating (0.05 cycles per degree, 1 Hz temporal frequency, eight randomized different directions). The stimuli were synchronized to individual image frames using frame-start pulses provided by ScanImage.

### Cortex: post hoc *anatomy*

After the loose-seal recording sessions, mice were anesthetized with a mixture of ketamine-xylazine (0.1 mg ketamine & 0.008 mg xylazine per gram body weight) and were trans-cardially perfused with 4% PFA in 1X DPBS. The brains were extracted and post-fixed overnight in the perfusing solution. The brains were sectioned at 50 µm thickness, blocked with 2% BSA+ 0.4 Triton-100 (in PBS) for 1 h at room temperature, incubated with primary antibody (Rb-anti-GFP, 1:500, Invitrogen, #G10362) for 2 days at 4℃, and secondary antibody (AlexaFluor 594 conjugated goat anti-Rb, 1:500, Invitrogen, #A-11012) overnight at 4℃. The sections were mounted on microscope slides in Vectashield hard-set antifade mounting medium with DAPI (H-1500, Vector). Samples were imaged using a TissueFAXS 200 slide scanner (TissueGnostics, Vienna, Austria) comprising an X-Light V2 spinning disk confocal imaging system (CrestOptics, Rome, Italy) built on an Axio Imager.Z2 microscope (Carl Zeiss Microscopy, White Plains, NY) equipped with a Plan-Apochromat 20x/0.8 M27 objective lens.

### Cortex: analysis of *in vivo* two-photon imaging

The acquired data was analyzed using MATLAB (population imaging) or Python (imaging during loose-seal recordings). In the MATLAB pipeline, for every recorded FOV, we selected ROIs covering all identifiable cell bodies using a semi-automated algorithm, and the fluorescence time course was measured by averaging all pixels within individual ROIs, after correction for neuropil contamination (r = 0.7), as described in detail in ^8^. We used one-way ANOVA tests (*P*< 0.01) for identifying cells with significant increase in their fluorescence signal during the stimulus presentation (responsive cells). We calculated ΔF/F_0_ = (F − F_0_)/F_0_, where F is the instantaneous fluorescence signal and F_0_ is the average fluorescence in the interval 0.7 s before the start of the visual stimulus. For each responsive cell we defined the preferred stimulus as the stimulus that evoked the maximal ΔF/F_0_ amplitude (averaging the top 25% of ΔF/F_0_ values during the 2 s of stimulus presentation). The half-decay time was calculated as follows: for each responsive cell we averaged its ΔF/F_0_ response to the preferred stimulus over five trials. We also calculated the standard deviation of the averaged baseline signal during 0.7 s before the start of the stimulus. Only cells where maximal ΔF/F_0_ amplitude was higher than four standard deviations above the baseline signal were included in the analysis. The time required for each trace to reach half of its peak value (baseline fluorescence subtracted) was calculated by linear interpolation. The fraction of cells detected as responsive was calculated as the number of significantly responsive cells over all the cells analyzed. The cumulative distribution of peak ΔF/ F_0_ responses included the maximal response amplitude from all analyzed cells, calculated as described above for each cell’s preferred stimulus. The orientation sensitivity index (OSI) was calculated as before ^6, 8^, by fitting the fluorescence response from individual cells to the eight drifting grating stimuli with two Gaussians, centered at the preferred response angle (R_pref_ ) and the opposite angle (R_opp_). The OSI was calculated as

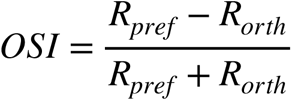

where R_orth_ is the orthogonal angle to the preferred angle.

The movies recorded during loose-seal recordings were motion-corrected and segmented with the python implementation of Suite2p (github.com/MouseLand/suite2p) ^65^. The ROI corresponding to the loose-seal recorded cell was then manually selected from the automatically segmented ROIs. For this dataset, we could calculate the neuropil contamination for most of the movies and got a distribution with a median of r_neu ∼ 0.8 (**Supp. Fig. 18**), so we used this value uniformly for neuropil correction. Calcium events were defined by grouping action potentials with a 20 ms inclusion window. Then we calculated ΔF/F_0_ = (F − F_0_)/F_0_, where F is the instantaneous fluorescence signal and F_0_ was defined separately for all calcium events as the mean fluorescence value of the last 200 ms before the first action potential in the group. We also calculated a global ΔF/F_0_ trace (ΔF/F_0_)_global_, where we used the 20^th^ percentile of the fluorescence trace in a 60 s long running window as the F_0,global_. In the analyses we only included calcium events where this (ΔF/F_0_)_global_ value was less than 0.5 right before the action potential, to include only events starting near baseline fluorescence values, in order to exclude non-linear summation and saturation. Traces in Fig. 4 were filtered with a Gaussian kernel (σ = 5 ms).

### Cerebellum: mouse surgeries

Young adult (postnatal day 42-98) male C57BL/6J (Jackson Labs) mice were anesthetized using isoflurane (2.5% for induction, 1.5% during surgery). A circular craniotomy (3 mm diameter) above medial crus I (2 mm left and 1 mm posterior to the midline junction of the interparietal and occipital bones). Viral suspension (200 nL) was injected in 2 locations near the center point at a depth of 300-400 µm. Constructs injected included: AAV2/1-*CAG*-FLEx-jGCaMP8 constructs (pGP-AAV-*CAG*-FLEx-jGCaMP8f-WPRE, Addgene plasmid #162382; pGP-AAV-*CAG*-FLEx-jGCaMP8m-WPRE, Addgene plasmid #162381; pGP-AAV-*CAG*-FLEx-jGCaMP8s-WPRE, Addgene plasmid #162380; pGP-AAV-*CAG*-FLEx-jGCaMP7f-WPRE, Addgene plasmid #104496; and pGP-AAV-*CAG*-FLEx-jGCaMP6f-WPRE, Addgene plasmid #100835; all viruses diluted to 4e12 GC/mL titer).

Purkinje cell-specific expression was induced by co-injection of virus expressing Cre under control of a promoter fragment from the Purkinje cell protein 2 (*Pcp2*; *a.k.a. L7*) gene (AAV2/1- *sL7*-Cre, 5.3e10 GC/mL titer) ^66^. A 3 mm diameter circular coverslip glued to a donut-shaped 3.5 mm diameter coverslip (no. 1 thickness, Warner Instruments) was cemented to the craniotomy using dental cement (C&B Metabond, Parkell Inc.). A custom titanium head post was cemented to the skull.

### Cerebellum: two-photon imaging

Head-restrained mice were allowed to freely locomote on a wheel. Imaging was performed with a custom-built two-photon microscope with a resonant scanner. The light source was a Mai Tai sapphire laser (Spectra Physics) running at 920 nm. The objective was a 16× CFI LWD Plan fluorite objective water immersion lens with 0.8 numerical aperture (Nikon). The detection path consisted of a bandpass filter (525/50 nm) and a 565 nm dichroic mirror directed towards a photomultiplier tube ( Hamamatsu). Images were acquired using Scan Image (vidriotechnologies.com) ^62^. Functional images (512 x 32 pixels, 215 x 27 µm^2^) of Purkinje cell dendrites (50–250 µm below the pia mater) were collected at 283 Hz. Laser power was up to 50 mW at the front aperture of the objective.

### Cerebellum: analysis of *in vivo* two-photon imaging

Purkinje cell dendrite movies were captured during free locomotion without applied stimulation. Movies were motion-corrected and converted to ΔF/F_0_ traces using the Python implementation of CaImAn ^67^. Individual events within the traces were identified by finding adjacent local maxima in ΔF/F_0_ variance that had one local maximum in the ΔF/F_0_ trace between them. Statistics for individual events were calculated by fitting the equation:

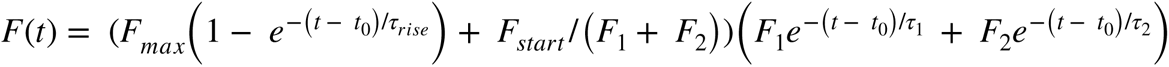

where *t_0_* is the start of the peak, *τ_rise_* is the rise time constant shaped by α; *τ_1_* and *τ_2_* are the decay time constants, *F_max_* is the maximum amplitude of the trace above the starting point *F_start_*, and *F_1_* and *F_2_* are component amplitudes.

### Spike-to-fluorescence model

We developed a phenomenological model that converts spike times to a synthetic fluorescence time series ^37^. This ‘spike-to-fluorescence’ (S2F) model consists of two steps. First, spikes at times {*t_k_*} are converted to a latent variable, c(t), by convolution with two double-exponential kernels:

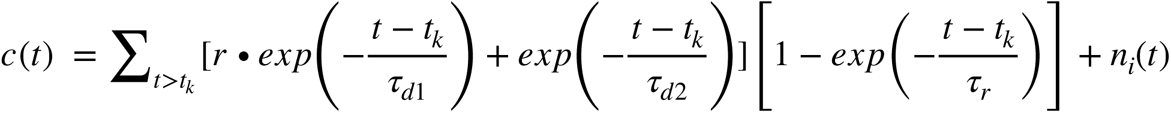

**τ**_r_ and **τ**_d1_, **τ**_d2_ are the rise time and decay times, respectively. In our model, we required **τ**_d1_< **τ**_d2_ (*i.e.*, fast and slow components), with r representing the ratio of the weight for the fast component to that of the slow one. *n_i_*(*t*)∼*N*_(_0,*σ* ^2^) is Gaussian-distributed ‘internal’ noise. c(t) was truncated at zero if noise drove it to negative values. We tested the performances of models with various choices of rise and decay times: (1) one rise and one decay time, (2) one rise and two decay times, and (3) two rise and two decay times. Using cross-validation, we found that one rise and two decay times models fit pyramidal cells (as described above), whereas interneurons were fit well by one rise and a single decay time (as described above with r = 0). Subsequently, c(t) was converted to a synthetic fluorescence signal through a sigmoidal function:

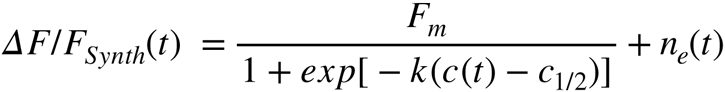

k is a non-linearity sharpness parameter, c_1/2_ is a half-activation parameter, and F_m_ is the maximum possible fluorescence change. *n_e_*(*t*)∼*N*(0,*σ* ^2^) is Gaussian-distributed external noise. For comparison, we also generated a S2F linear model with *ΔF*/*F_Synth_*(*t*) = *F_max_c*(*t*) + *F*_0_, where F_max_ is a scaling parameter (we kept the naming as max to clarify the relationship to other models); F_0_ is the baseline.

### Variance explained

Variance explained measures the goodness-of-fit of an S2F model, as 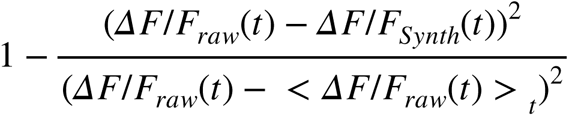. Here we used only the period t with spike rate >0 Hz after spikes in the calculation, where the instantaneous spike rate at time *t* is estimated by a boxcar-rolling average over a 600 ms time window.

### Statistics

Exact statistical tests used for each comparison, as well as *n*, are listed in the main text and figure legends. Box-whisker plots throughout the manuscript indicate the median and 25^th^–75^th^ percentile range; whiskers indicate the shorter of 1.5 times the 25^th^–75^th^ range or the extreme data point. For Fig. 3E, full statistics are: jGCaMP7f vs XCaMP-Gf: P = 1.0; jGCaMP7f vs jGCaMP8f: P=0.013; jGCaMP7f vs jGCaMP8m: P = 0.029; jGCaMP7f vs jGCaMP8m: P = 0.010; jGCaMP7f vs jGCaMP8s: P = 0.010; XCaMP-Gf vs jGCaMP8f: P = 1.0; XCaMP-Gf vs jGCaMP8m: P = 1.0; XCaMP-Gf vs jGCaMP8s: P = 0.0027; jGCaMP8f vs jGCaMP8m: P = 1.0; jGCaMP8f vs jGCaMP8s: P < 0.001; jGCaMP8m vs jGCaMP8s: P < 0.001. \**P* < 0.05; \*\*\**P* < 0.001; ns, not significant. For Fig. 3F, data passed Shapiro-Wilk normality test (α = 0.05 level). For Fig. 3F, full statistics are: jGCaMP7f vs jGCaMP8f: P = 0.83; jGCaMP7f vs jGCaMP8m; P = 0.0184; jGCaMP7f vs jGCaMP8s: P < 0.001; jGCaMP8f vs jGCaMP8m: P < 0.001; jGCaMP8m vs jGCaMP8s: P = 0.23. For Fig. 3G, full statistics are: jGCaMP7f vs jGCaMP8f: P < 0.001; jGCaMP7f vs jGCaMP8m; P = 1.0; jGCaMP7f vs jGCaMP8s: P < 0.001; jGCaMP8f vs jGCaMP8m: P < 0.001; jGCaMP8m vs jGCaMP8s: P < 0.001 (Kruskal-Wallis test with Dunn’s multiple comparison test was used to compare the magnitude of response across groups).

### *In situ* stability of the jGCaMP8 sensors

The jGCaMP8 indicators were about half as bright as 7f in *Drosophila* adult and larval NMJ (and had lower levels of protein expression), whereas in mouse cortex they had similar brightness and expression levels. The mechanisms underlying the decreased expression levels in *Drosophila* are unclear but may involve species-specific variation in the “N-end rule” ^68^ and ubiquitin exoligation at the N-terminus, which was shortened by truncation of the RSET affinity tag in the jGCaMP8 variants.

### Reagent distribution and data availability

DNA constructs and AAV particles of jGCaMP8s, jGCaMP8m, and jGCaMP8f (pCMV, pAAV- *synapsin-1*, pAAV-*synapsin-1*-FLEX, and pAAV-*CAG*-FLEX) have been deposited at Addgene (#162371-162382). Sequences have been deposited in GenBank (#OK646318-OK646320). *Drosophila* stocks were deposited at the Bloomington Drosophila Stock Center (http://flystocks.bio.indiana.edu); *Drosophila* UAS and lexAOp plasmids are at Addgene (#162383-#162388). Fish lines are available on request. Most datasets generated for characterizing the new sensors are included in the published article (and its Supplementary Information files). Additional datasets are available from the corresponding authors on reasonable request.

## Acknowledgements

This work is part of the GENIE Project at the Howard Hughes Medical Institute, Janelia Research Campus. The GENIE Project is led by JPH, WLK, and IK; the Steering Committee includes GCT, KS, ERS, and LLL. This work was supported by the Howard Hughes Medical Institute, and NIH U19 NS104648 and NIH R01 NS045193 to SS-HW and NIH F32 MH120887 to GJB. We would like to thank the Berkeley Center for Structural Biology for use of beamline 8.2.1. We are grateful to the Viral Tools, Cell and Tissue Culture, Molecular Biology, Media Prep, Vivarium/Aquarium, Anatomy and Histology, and Fly Facility Shared Resources at Janelia for technical assistance with numerous parts of the project.

## Author contributions

Designed project: YZ, ERS, LLL. Led the project: YZ, KS, IK, JPH, LLL. Optimized peptides for grafting: YZ, ERS. Mutagenesis: YZ, GT, JPH. Fly: DB, JZ, GCT. Fish: JXL, SN, MBA, ZW. Cerebellum: GJB, SS-HW. Photophysics: YZ, RP, IK. Crystallography: YZ. Purified protein experiments: YZ. Cortex: MR, YL, KS. Simultaneous imaging & electrophysiology: MR. Spike to fluorescence: ZW, MR, KS. Cultured neuron screen design: IK, JPH, KS. Cultured neuron experiments: YL, DR, AT, CJO, RZ, JPH, IK. Histology: DR, AT, IK. Analysis methodologies: YZ, MR, YL, KS, IK, DB, JZ, ZW. Coordination of shipments & deposits: YZ, JPH. Wrote paper: YZ, MR, YL, KS, DB, LLL, with contributions from all the authors.

## Supplementary Tables

**Supp. Table 1.** Biophysical properties of initial sensors with different calmodulin-binding peptides used in this study.

**Supp. Table 2.** Sensor variants run through cultured neuron assay. ΔF/F_0_, half-rise time, time-to-peak, half-decay time, signal-to-noise ratio, and normalized F_0_ given for each of 1, 3, 10, 160 AP. *P*-values computed by the Mann-Whitney *U* test for each variant, compared to 6s, given. Nucleic acid and protein sequence of each variant, as well as detailed name (if any) shown.

**Supp. Table 3.** Crystal structure determination of jGCaMP8.410.80.

**Supp. Table 4.** Comparison of sensitivity and kinetics of jGCaMP8 to XCaMP-G, -Gf, and -Gf0 sensors. Colors in each cell indicate whether the value was significantly higher for jGCaMP8 (yellow), XCaMP (blue), or not statistically different (no color), as evaluated with Dunn’s multiple comparisons test (*P*-values in cells).

**Supp. Table 5.** Data on recorded cells, including GECI, mouse, neuron # per each mouse, putative cell type (pyramidal, interneuron), recording length, recording mode, total # of recorded spikes, and median spike SNR.

**Supp. Table 6.** Biophysical characterization in purified protein. Apparent *K_d_*, Hill coefficient (cooperativity), saturating ΔF/F_0_, off and on-rates (8f is modeled as having a 2-component decay curve), pK_a_ in both Ca^2+^-free and Ca^2+^-saturated states, excitation, emission, and absorption wavelength maxima, and extinction coefficient of Ca^2+^-free and Ca^2+^-saturated states. Values are *n* = 3, mean *±* std. err.

**Supp. Table 7.** Statistics of the degree of nonlinearity of sensors measured by the difference of variance explained by S2F sigmoid from linear model (mean ± std.dev.).

**Supp. Table 8.** Statistics of S2F parameter fits (mean ± std.dev.).

## Supplementary Figures

**Supp. Fig. 1.**
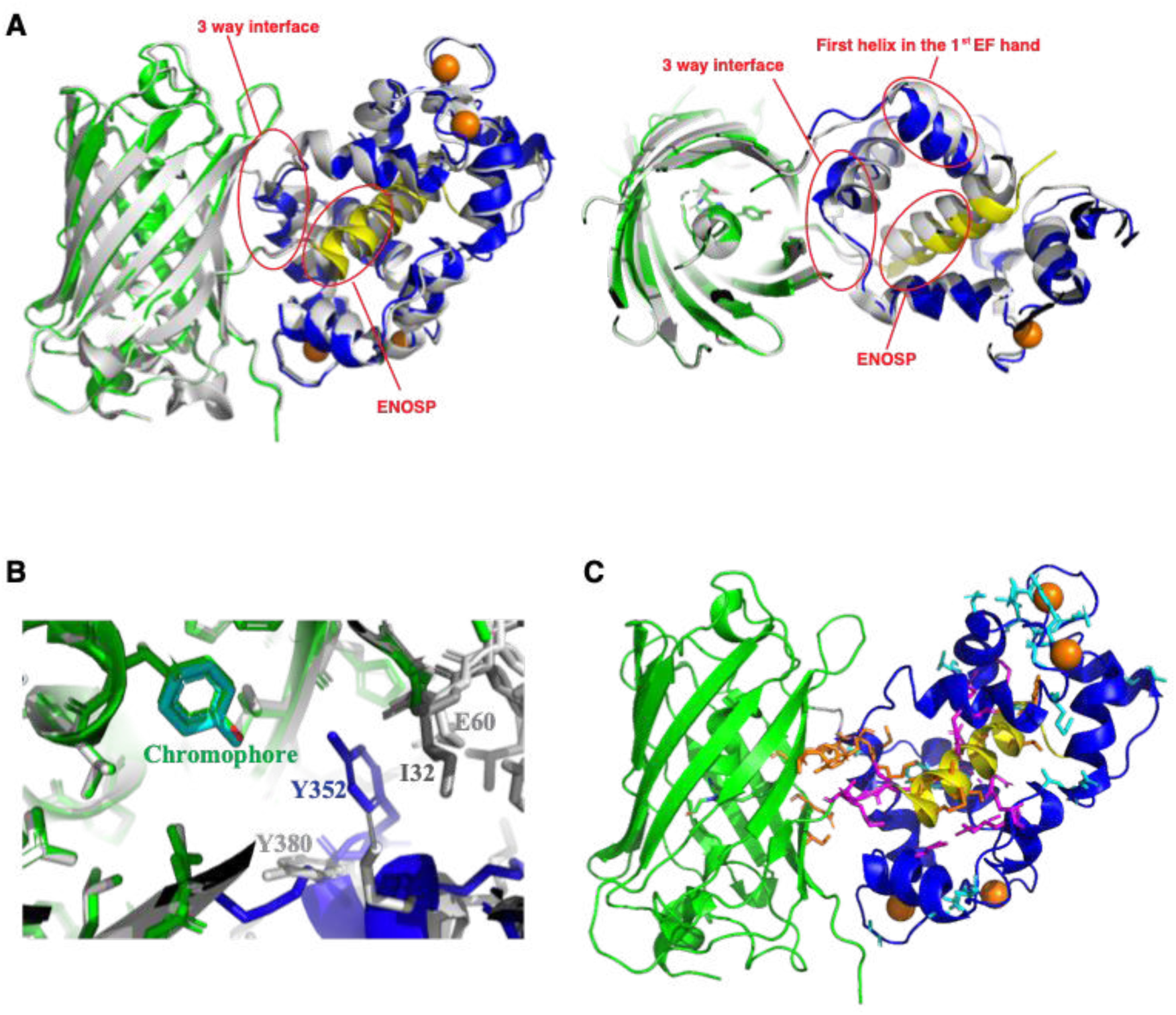
Crystal structure of jGCaMP8.410.80. ENOSP (yellow), linker 1 (ENOSP-cpGFP, grey), linker 2 (cpGFP-CaM, grey), cpGFP (green), CaM (blue), Ca^2+^ ions (orange). **A.** Overlay of the structures of jGCaMP8.410.80 and GCaMP5G (light grey). Left: side view. Right: top view. **B.** A close up of the chromophore area in the structures of jGCaMP8.410.80 and GCaMP5G. Ile32 (dark gray) in Linker 1 of jGCaMP8.410.80 facilitates closer interaction of Tyr352 (blue) with the GFP chromophore. The corresponding residues in GCaMP5G, Glu60 and Tyr380, are depicted in gray. **C.** Individual residue mutations screened in this study, shown on the structure of jGCaMP8.410.80. Sixteen initial interface positions are in orange. Ten subsequently mutated CaM positions are in magenta. Mutations based on the FGCaMP sensor are in cyan.

**Supp. Fig. 2.**
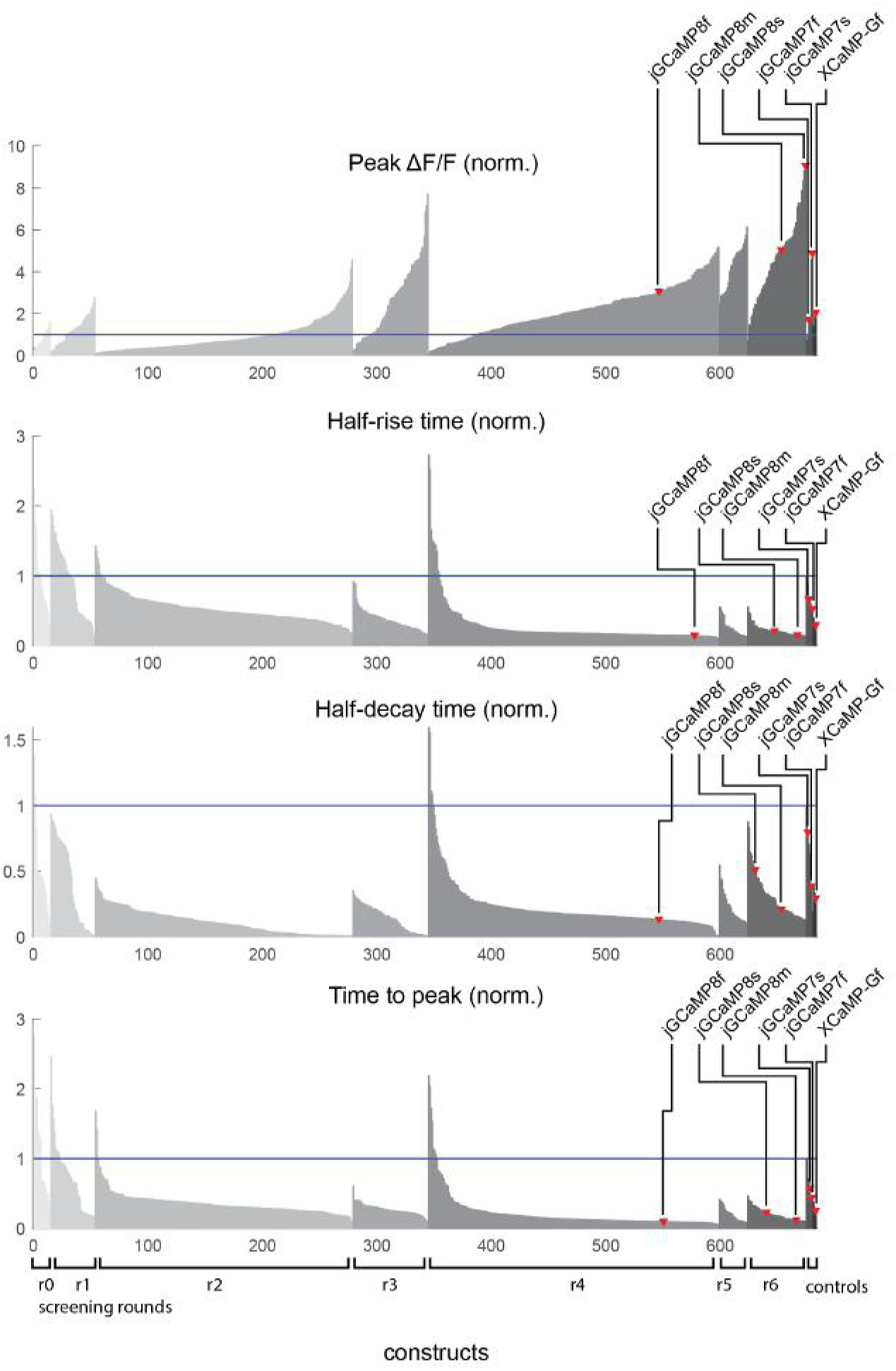
Results of cultured neuron 1-AP field stimulation screen (*n*=683 constructs with detectable 1-AP responses; **Methods**). All results are normalized to GCaMP6s controls (blue line) and listed in ranked order (increasing for peak ΔF/F_0_, decreasing for all others) from each screening round. Sensor engineering took place over seven rounds: Round 0: Graft peptides, screen linkers. Round 1: Site-saturation mutagenesis of 16 interface positions: 7 in ENOSP, 4 on cpGFP, and 5 on CaM. Round 2: Combination of beneficial mutations to date. Round 3: Site-saturation mutagenesis of 10 additional CaM positions surrounding ENOSP. Round 4: Graft mutations from FGCaMP. Round 5 and 6: Two additional rounds of combination of beneficial mutations.

**Supp. Fig. 3.**
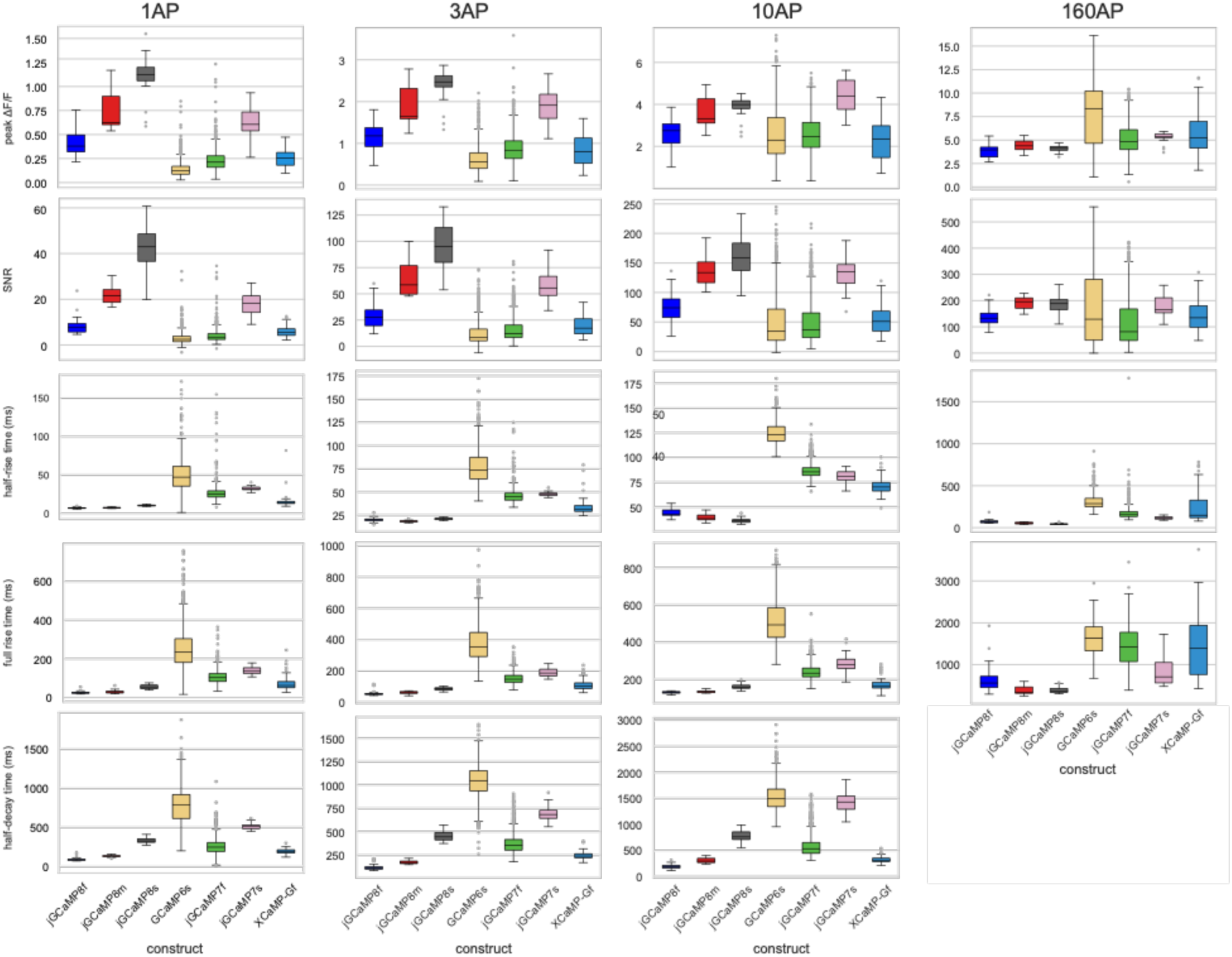
Response characteristics of jGCaMP8 indicators to 1, 3, 10, and 160 field stimulation pulses (45 V, 83 Hz). Half-decay at 160 pulses is not reported because cell fluorescence typically does not decay to baseline during our imaging time (6 s after stimulus onset).

**Supp. Fig. 4.**
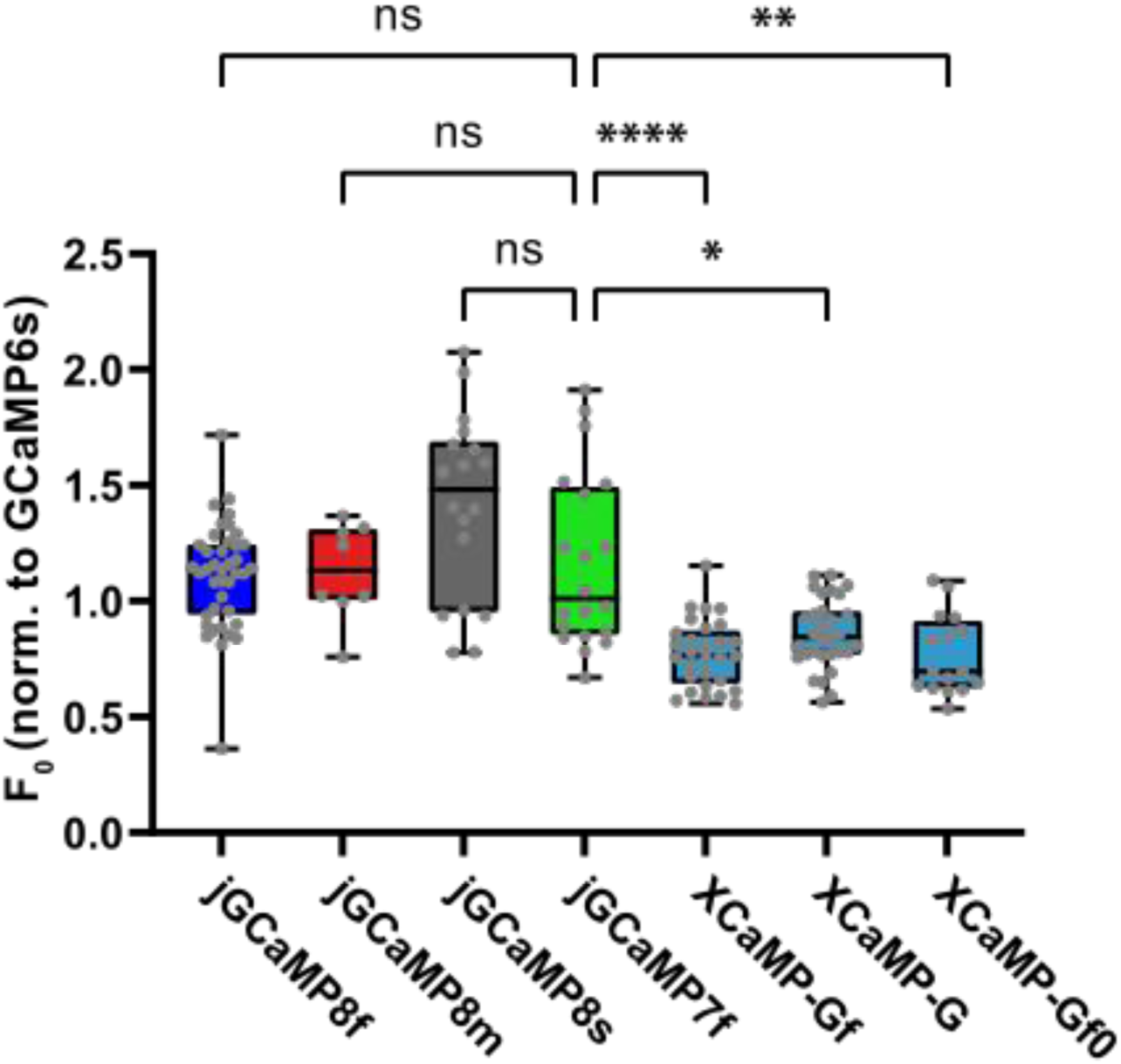
Baseline sensor brightness. The jGCaMP8 series exhibited similar baseline fluorescence compared to 7f, but XCaMP sensors were significantly dimmer (*H(6) = 71.77*, *P*<0.0001, Kruskal-Wallis test; Dunn’s multiple comparisons test with 7f as control). n.s.: not significant (*P*>0.99). \**P*=0.012; \*\**P*=0.0012; \*\*\*\**P*<0.0001. Each point represents a single well. Data from five 96-well plates over 2 weeks of separate transfections.

**Supp. Fig. 5.**
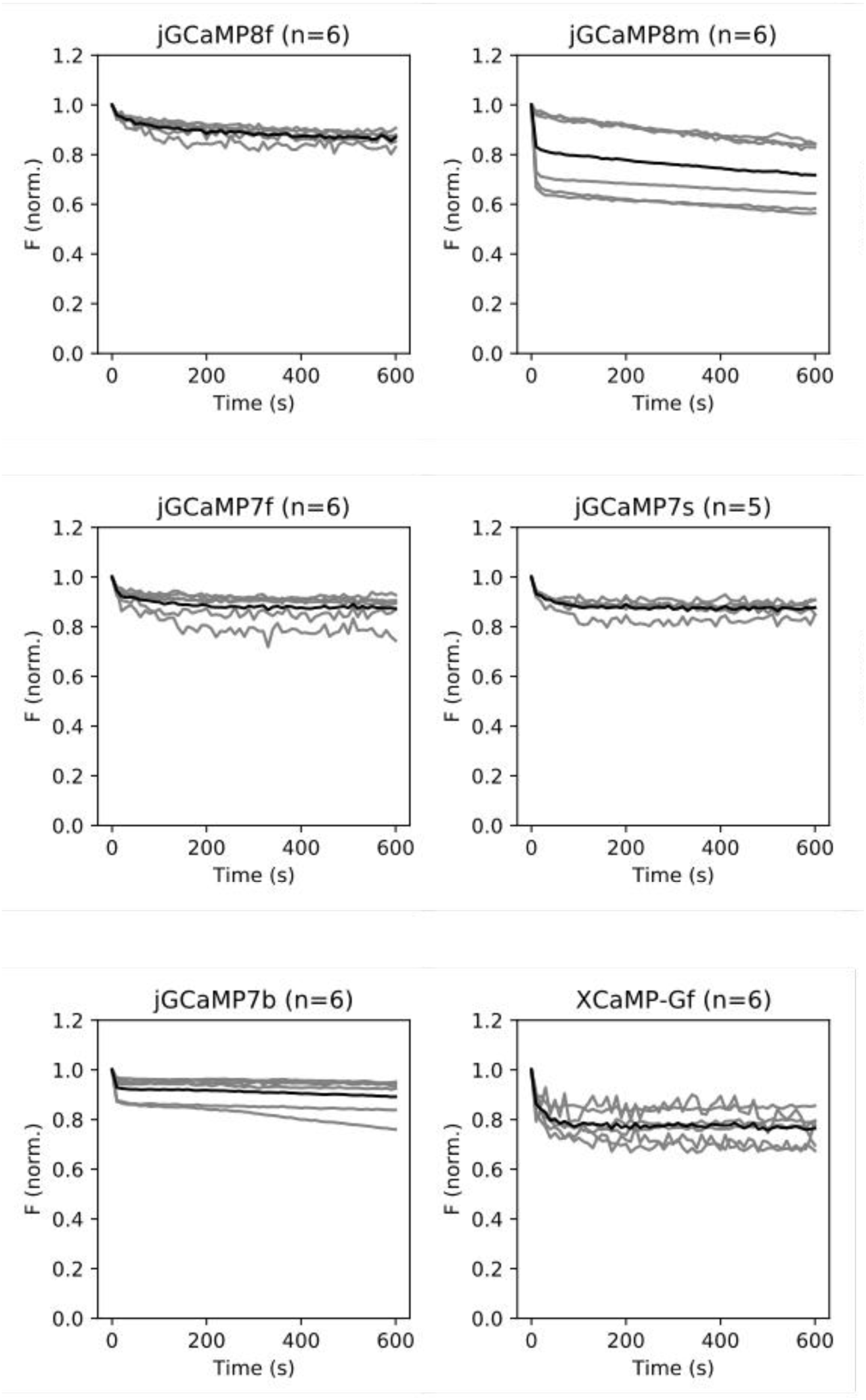
Photobleaching of jGCaMP8, jGCaMP7, and XCaMP variants in neuron cell culture. Grey lines: individual cells, black lines: mean. Each cell’s fluorescence trace was normalized to the initial value. *N* values indicate number of cells (*n*=1 well per variant, *n*=1 transfection day). After continuous illumination for 10 minutes, neurons transfected with jGCaMP8 variants lost on average 13-28% of their initial fluorescence. 8m exhibited biphasic bleaching: a rapid phase consisting of ∼15% fluorescence loss within 10 s followed by a slower phase (10% within 10 minutes). Of the other variants, 7c also exhibited this property. We noticed considerable variability in the photobleaching rates within individual neurons, possibly stemming from expression level and differences in baseline brightness in each neuron as a function of intracellular resting [Ca^2+^].

**Supp. Fig. 6.**
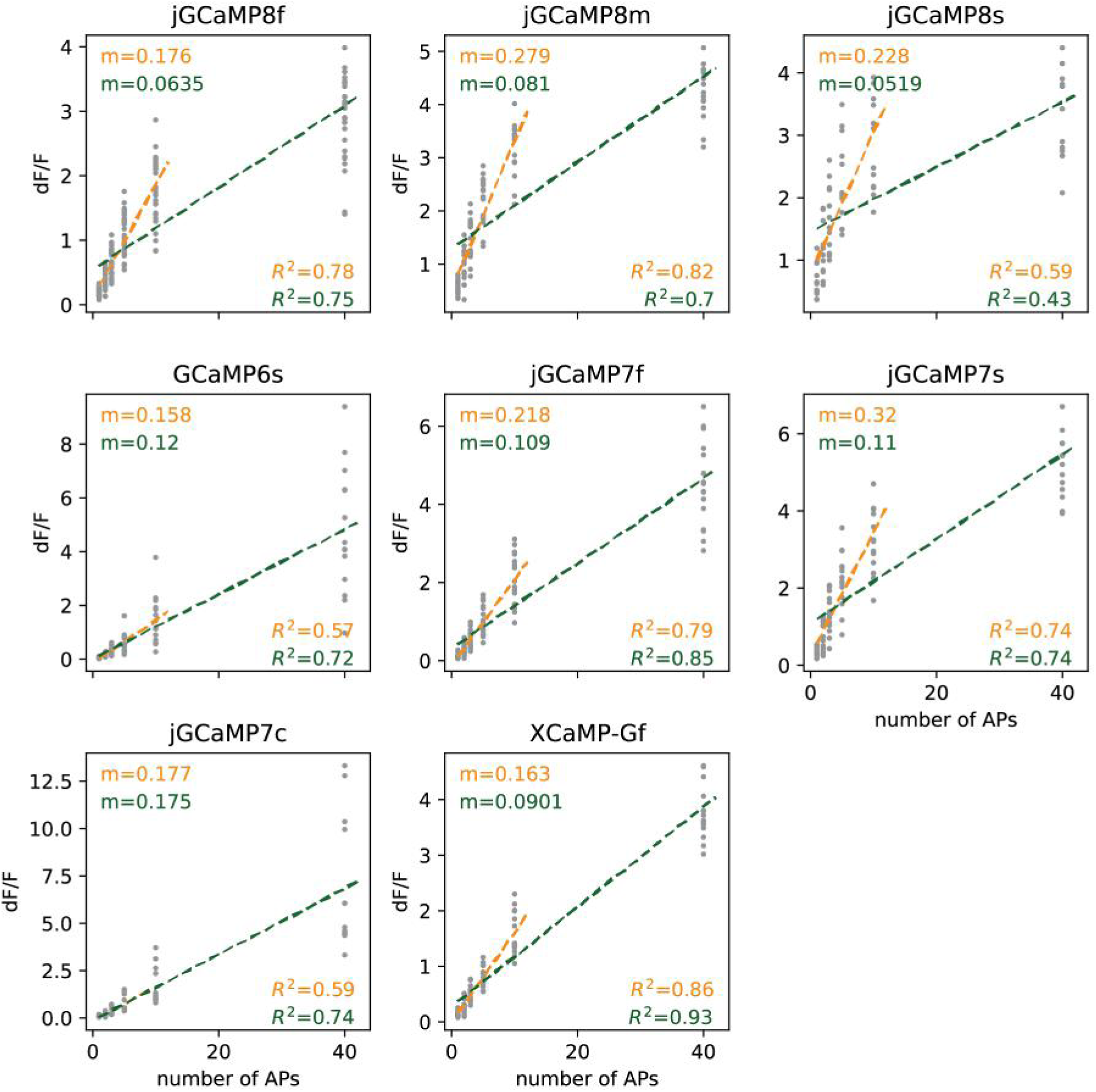
Linearity of ΔF/F_0_ of jGCaMP8, jGCaMP7, and XCaMP variants in cultured neurons. Each gray dot represents a single well. ΔF/F_0_ values in the 1-10 and 1-40 pulse range were fit to a linear model (orange and green, respectively). The slopes (*m*) and R^2^ values are reported for each fit. 8f: 29 wells, 594 neurons; 8m: 16 wells, 408 neurons; 8s: 12 wells, 121 neurons; 6s: 14 wells, 187 neurons; 7s: 14 wells, 177 neurons; 7c: 13 wells, 117 neurons; XCaMP-Gf: 14 wells, 194 neurons; 2 independent transfections, four 96-well plates. The jGCaMP8 sensors were moderately linear and exhibited a large slope in the 1-10 AP range (0.59 ≤ R^2^ ≤ 0.82; 0.18 ≤ m ≤ 0.28), but less linear and exhibited a lower slope in the 1-40 AP range (0.43 ≤ R^2^ ≤ 0.75; 0.052 ≤ m ≤ 0.081). On the other hand, while 6s, 7c, and XCaMP-Gf maintained their linearity throughout the 1-40 AP range, they had generally lower slopes in the 1-10 AP range (0.16 ≤ m ≤ 0.18).

**Supp. Fig. 7.**
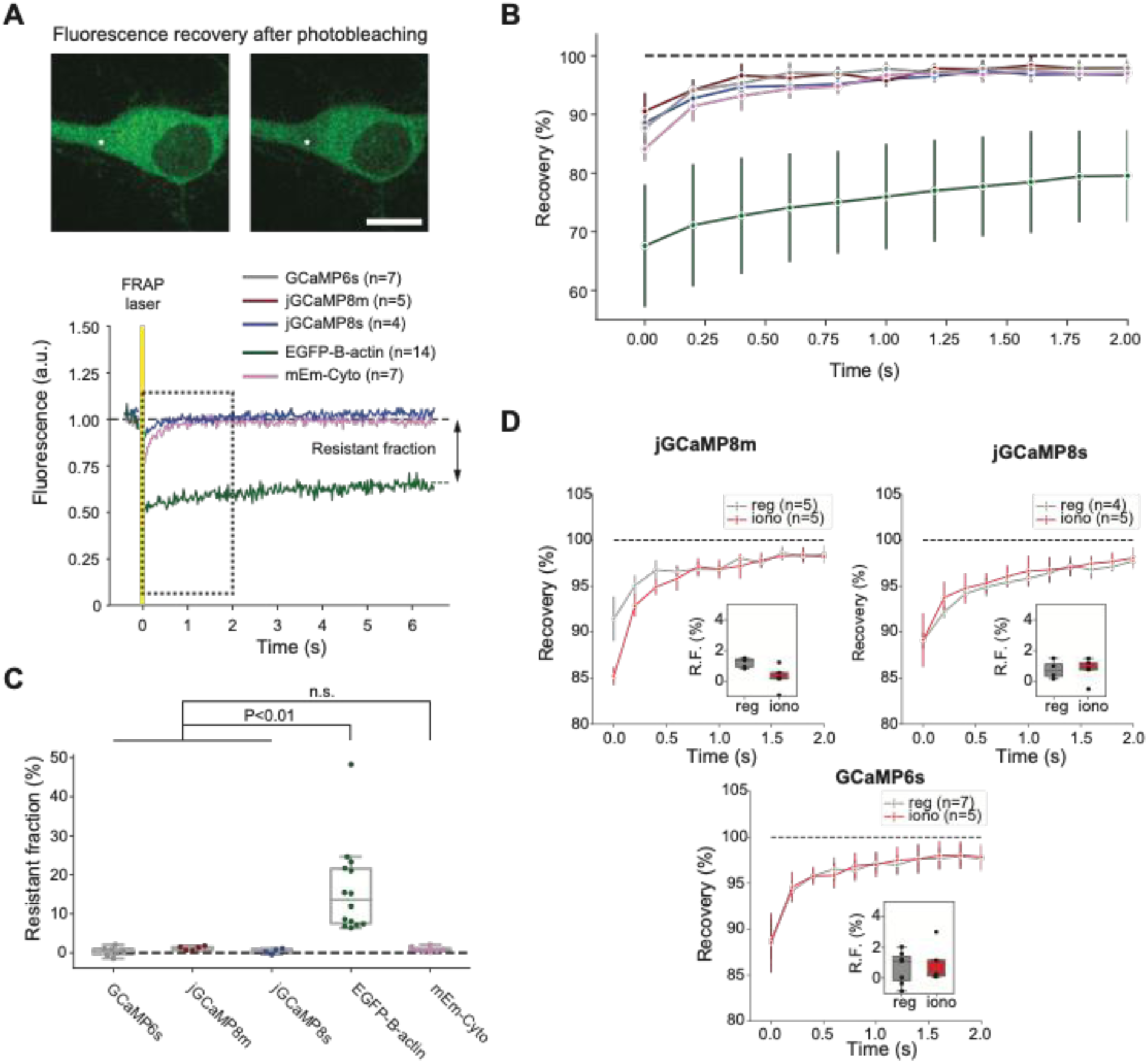
Sensor diffusion in cultured neurons studied with fluorescence recovery after photobleaching (FRAP). A. Top: Images of a representative cultured neuron expressing 8m before (left) and immediately after (right) laser illumination. Asterisk indicates bleached region. Bottom: Representative single-trial FRAP curves for 8s (blue), cytoplasmic mEmerald (mEm-Cyto; pink) and EGFP-β-actin (green), normalized to pre-stimulation fluorescence values and aligned to the FRAP laser pulse (yellow). Boxed area denotes zoomed-in region shown in B. *N* values indicate number of neurons tested in each condition for subsequent panels. Scale bar: 10 µm. B. Recovery curves of all tested variants (mean ± std.dev.). For clarity, only every 10^th^ point in the trace is plotted. The color scheme is the same as in A – this panel also shows 6s (grey) and 8m (dark red). C. Resistant fractions. The resistant fractions of 6s (0.3 ± 1.2%), 8m (1.3 ± 0.5%), and 8s (0.4 ± 0.7%) were not significantly different from a cytosolic GFP marker (mEm-Cyto, 0.9 ± 0.7%), but were significantly different from actin-bound GFP (EGFP-β-actin, 16.1 ± 11.4%; Welch’s ANOVA with Dunnett’s T3 multiple comparisons test; n.s.: *P*>0.45). D. Recovery curves of 8m, 8s and 6s, without (“reg”) or with (“iono”) added ionomycin to saturate sensor with Ca^2+^ (**Methods**).

**Supp. Fig. 8.**
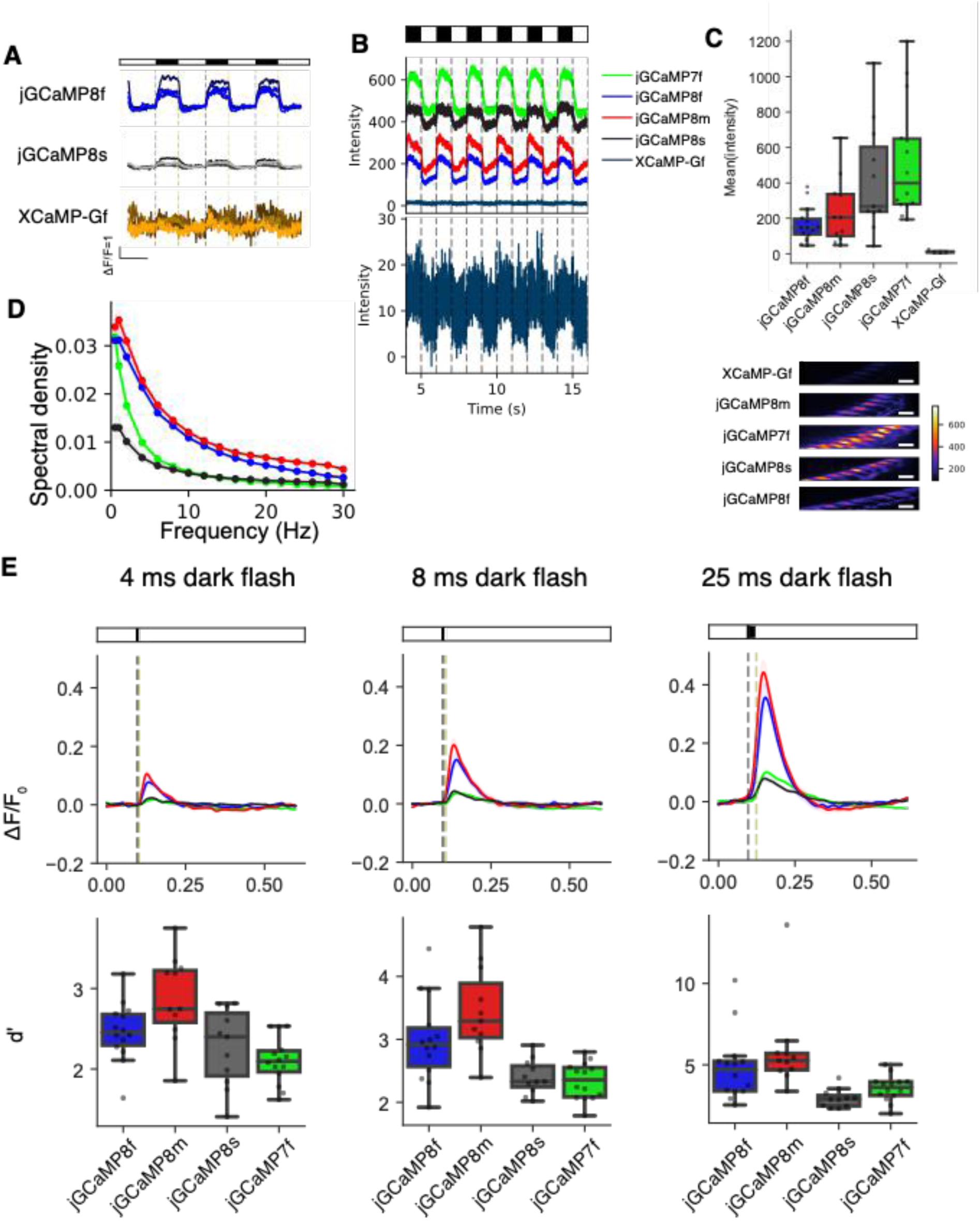
jGCaMP8 sensor characterization in adult *Drosophila* L2 visual system assay. **A**. Responses of 8f, 8s, and XCaMP-Gf to the visual stimulus, as in Fig. 2B. B. Raw fluorescence intensity counts for the five sensors tested. Inset below: XCaMP-Gf shown with y-axis ∼30x smaller. C. Top: quantification of data from panel B; total *n* for each variant: 7f: 14 flies, 8f: 14, 8m: 11, 8s: 11, XCaMP-Gf: 4. Bottom: images of mean intensity over the stimulation period, with color scale constant between variants. Scale bar, 5 *μ*m. The jGCaMP8 indicators were somewhat dimmer than 7f. D. Spectral power density measured from L2 responses at stimulation frequencies ranging from 0.5 to 30 Hz. E. ΔF/F_0_ responses to dark flashes 4, 8, or 25 ms in duration. Top: fluorescence traces show the mean and s.e.m. Bottom: box plots showing the discriminability index d’. At 4 ms duration, Kruskal-Wallis test finds *P*=2.3E-3 and pairwise Dunn’s multiple comparison test to 7f as follows: 8f=0.03, 8m=2.0E-4, and 8s=0.24. At 8 ms duration, Kruskal-Wallis test find *P*=3.5E-5 and pairwise Dunn’s multiple comparison test to 7f as follows: 8f=3.5E-3, 8m=2.8E-5, and 8s=0.73. At 25 ms duration, Kruskal-Wallis test find *P* =3.4E-5 and pairwise Dunn’s multiple comparison test to 7f as follows: 8f=0.074, 8m=1.6E-3, and 8s=0.11. Unless otherwise stated, all statistics include the following numbers tested: 8f=14, 8m=11, 8s=11, and 7f=14.

**Supp. Fig. 9.**
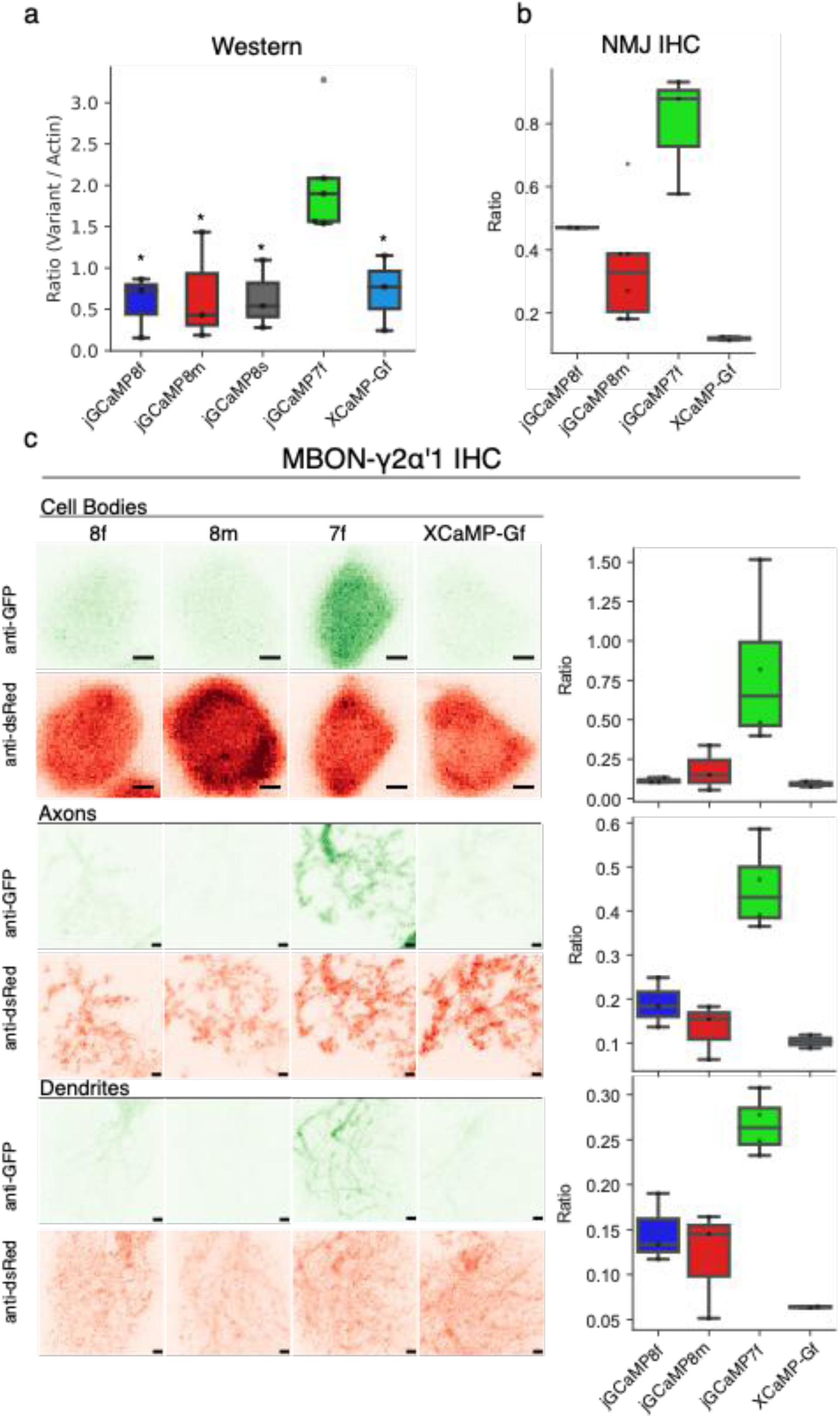
Expression of the GCaMP variants in adult fly visual system and larval neuromuscular junction. A. Western blot analysis comparing protein expression between GCaMP variants. Ratio is the band intensity levels from a variant divided by the band intensity from the actin loading control. Multi-comparison Kruskal-Wallis finds *P*=0.038 and pairwise Dunn’s multiple comparison test to 7f as follows: 8f=0.011, 8m=0.019, 8s=0.024, and XCaMP=0.038. Numbers tested are as follow: 8f=3, 8m=3, 8s=3, 7f=5, and XCaMP=3. The jGCaMP8 and XCaMP variants expressed ∼3x less protein than 7f in L2 neurons. B. Box plot comparing immunostaining at the NMJ. Ratio is the intensity from stain targeting variant divided by intensity from a myr::tdTomato co-expressed with the variant. Multi-comparison Kruskal-Wallis finds *P*=0.029 and pairwise Dunn’s multiple comparison test to 7f as follows: 8f=0.37, 8m=0.039, and XCaMP=4.2E-3. Numbers tested are as follow: 8f=2, 8m=6, 7f=3, and XCaMP=2. C. Immunostaining females expressing GCaMP variants and myr::tdTomato in MBON-γ2α’1. Left, images from cell bodies (top), axons (middle), and dendrites (bottom). Scale bar is 1 µm. Green images show variant expression while red images show myr::tdTomato expression. Right, box plots quantify the ratio between intensity from the variant to the myr::tdTomato. Multi-comparison Kruskal-Wallis for cell body finds *P*=0.05. Multi-comparison Kruskal-Wallis for axon finds *P* =0.032 and *P*-values from pairwise Dunn’s multiple comparison test as follows: 8f=0.13, 8m=0.018, and XCaMP=0.010. Multi-comparison Kruskal-Wallis for dendrite finds *P*=0.040 and p-values from pairwise Dunn’s multiple comparison test as follows: 8f=0.079, 8m=0.034, and XCaMP=0.010. Numbers tested are as follows: 8f=3, 8m=3, 7f=4, and XCaMP=2. The jGCaMP8 variants expressed ∼3x less protein than 7f in L2 neurons.

**Supp. Fig. 10.**
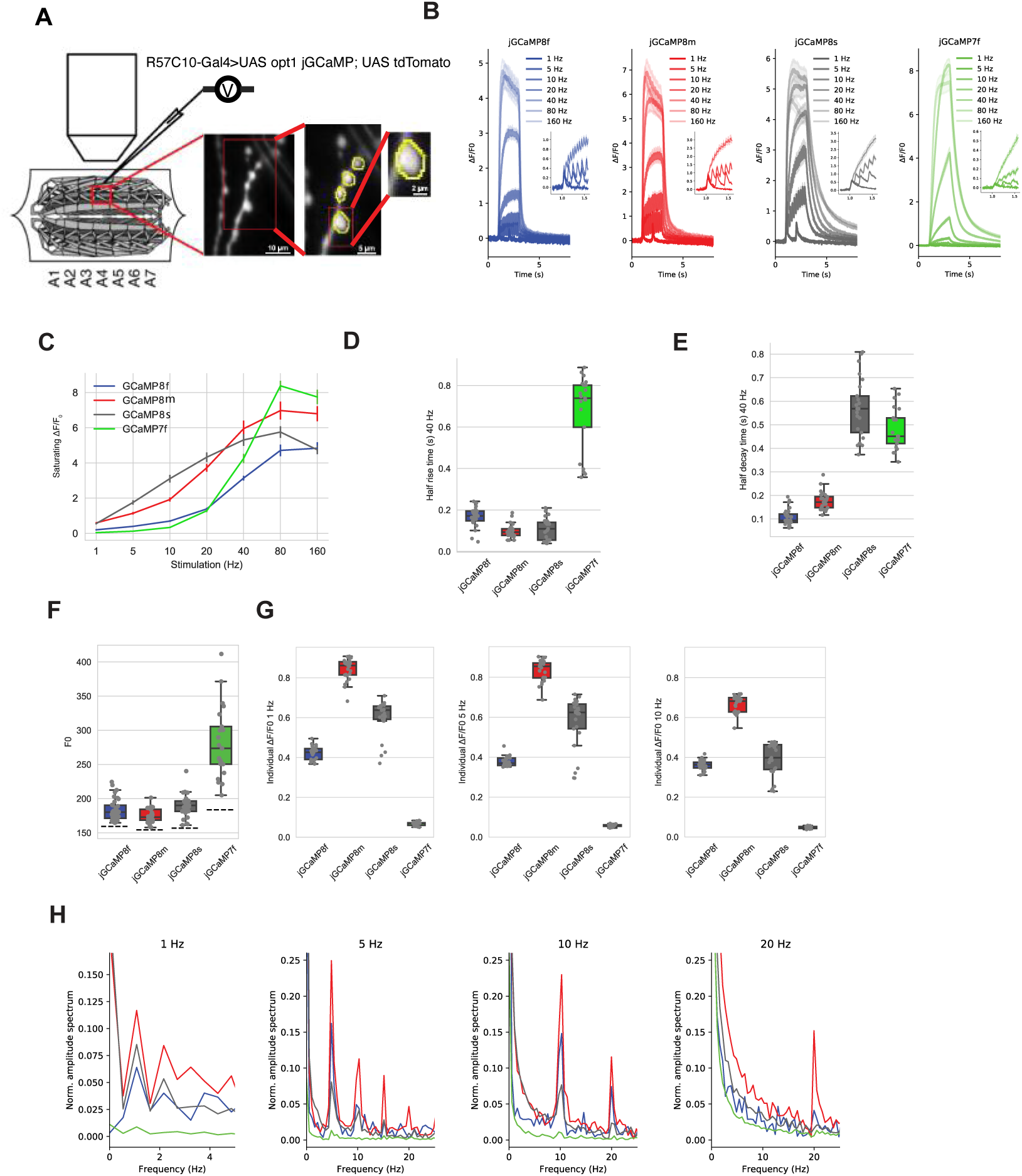
Characterization of GCaMP variants in larval neuromuscular junction. A. Design of larval NMJ experiments. B. Fluorescence response to 1, 5, 10, 20, 40, 80 and 160 Hz stimulation (2 s) of motor axons. Inset: zoomed response to 1, 5, 10 and 20 Hz. 8s showed superior response from 1-20 Hz and 7f above 80 Hz, where signals saturated. C. Saturating ΔF/F_0_ to 2 s motor axon stimulation at 1, 5, 10, 20, 40, 80 and 160 Hz. Mean and s.e.m. shown. D. Half-rise time from stimulus onset to saturated peak under 40 Hz stimulation. Half-rise time at 40 Hz stimulation was markedly shorter than 7f for all jGCaMP8 variants. E. Half decay-time from stimulus end to baseline under 40 Hz stimulation. Half-decay time was much shorter than 7f for 8f and 8m. F. F_0_ for each sensor. Dash line indicates the background fluorescence level. Resting fluorescence for the jGCaMP8 variants was lower than 7f. G. Individual responses to 1, 5, and 10 Hz stimulation. The jGCaMP8 series detect individual stimuli much better than 7f. H. Power spectral density normalized to 0 Hz for responses to 1, 5, 10, and 20 Hz stimulation. Colors as above. Power spectral analysis confirms the performance of the jGCaMP8 indicators, with 8m performing the best at all frequencies, particularly at the high end – 8m shows strong power at 20 Hz trains, whereas 7f is negligible. Panels B-G: Each data point represents a single bouton. # of boutons per line are: 8f, 27; 8m, 25; 8s, 25; 7f, 21. Boutons are from five individuals per line.

**Supp. Fig. 11.**
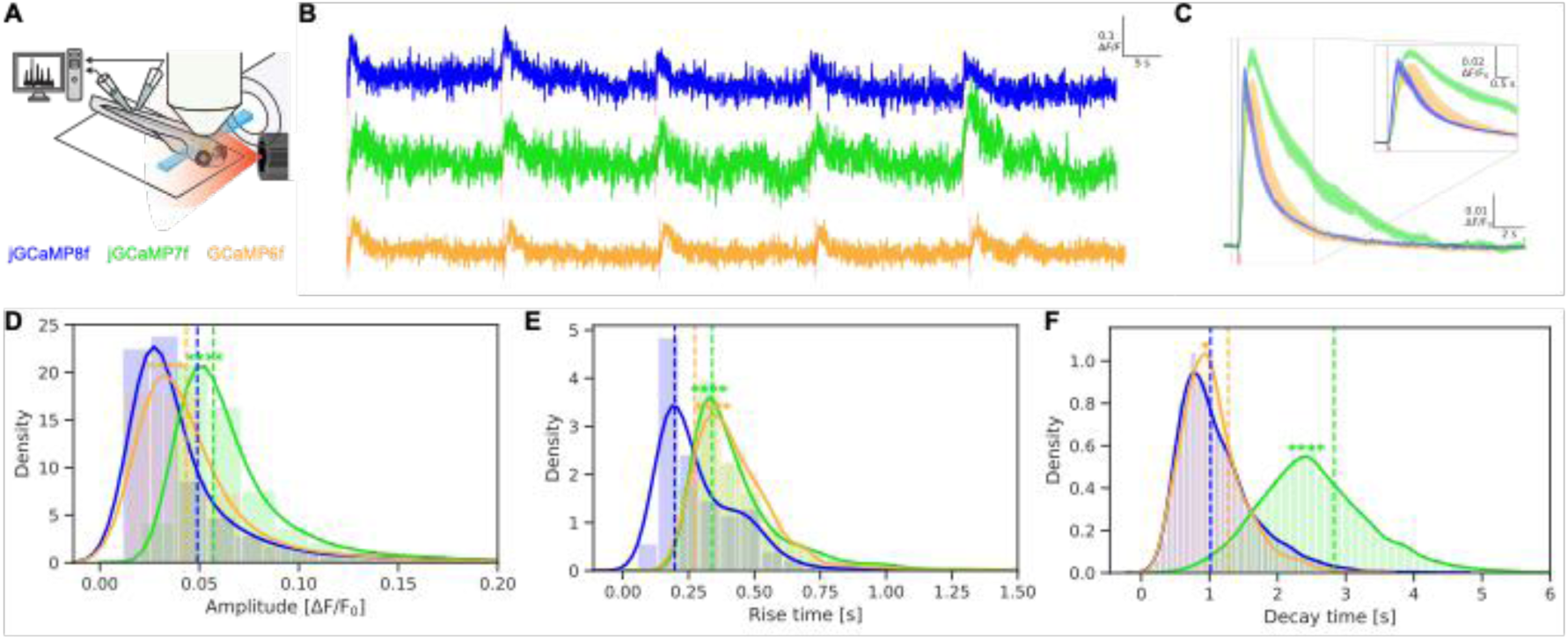
jGCaMP8f sensor characterization in larval zebrafish. A. Larval zebrafish were presented a series of short flashes while being subjected to fast multiplane light sheet calcium imaging and ventral root recording (proxy for behavior). Images were motion-corrected and segmented to give cell segments. Flash-responsive cells were extracted using a visual regressor (see Methods) and further analyzed. Total number of cells extracted – 8f: 5101; 7f: 7246; 6f: 2486. B. Representative single-cell responses of 8f, 7f and 6f to the visual stimulus. C. Variant mean. Single-cell responses were averaged to compute the mean response for each fish, which were then averaged again for the variant mean. Error bars indicate s.e.m. calculated with fish means. Number of fish tested – 8f: 5; 7f: 4; 6f: 4. D-F. Histogram (bars) and kernel density estimates (lines) of single-cell response amplitudes (D), half-rise time (E) and half-decay time (F) across all fish for each variant. The vertical dotted lines indicate the respective statistic for the variant means (C) for comparison. D. Amplitudes of the variant means – 8f: 0.0490; 7f: 0.0571; 6f: 0.0433. E. Half-rise times of the variant means – 8f: 0.197; 7f: 0.340, 6f: 0.0274. F. Half-decay times of the variant means – 8f: 1.02; 7f: 2.83; 6f: 1.28. Two-sample Kolmogorov–Smirnov tests (KS tests) were performed between 8f and the other densities to determine similarity of distributions. All 7f and 6f distributions were significantly different from their 8f counterparts in all three statistics as determined by two-sided KS tests. The results of one-sided KS tests asking if 8f is smaller than its counterparts is indicated in the figure according to the following n.s.: not significant (P>0.05); *P≤0.05; **P≤0.01; ***P≤0.001; ****P≤0.0001. 8f has significantly lower amplitudes, half-rise time and half-decay time than the other variants. One-sided KS tests for greater 8f values were also performed but was only significant for comparisons with 6f half-decay times (P≤0.0001).

**Supp. Fig. 12.**
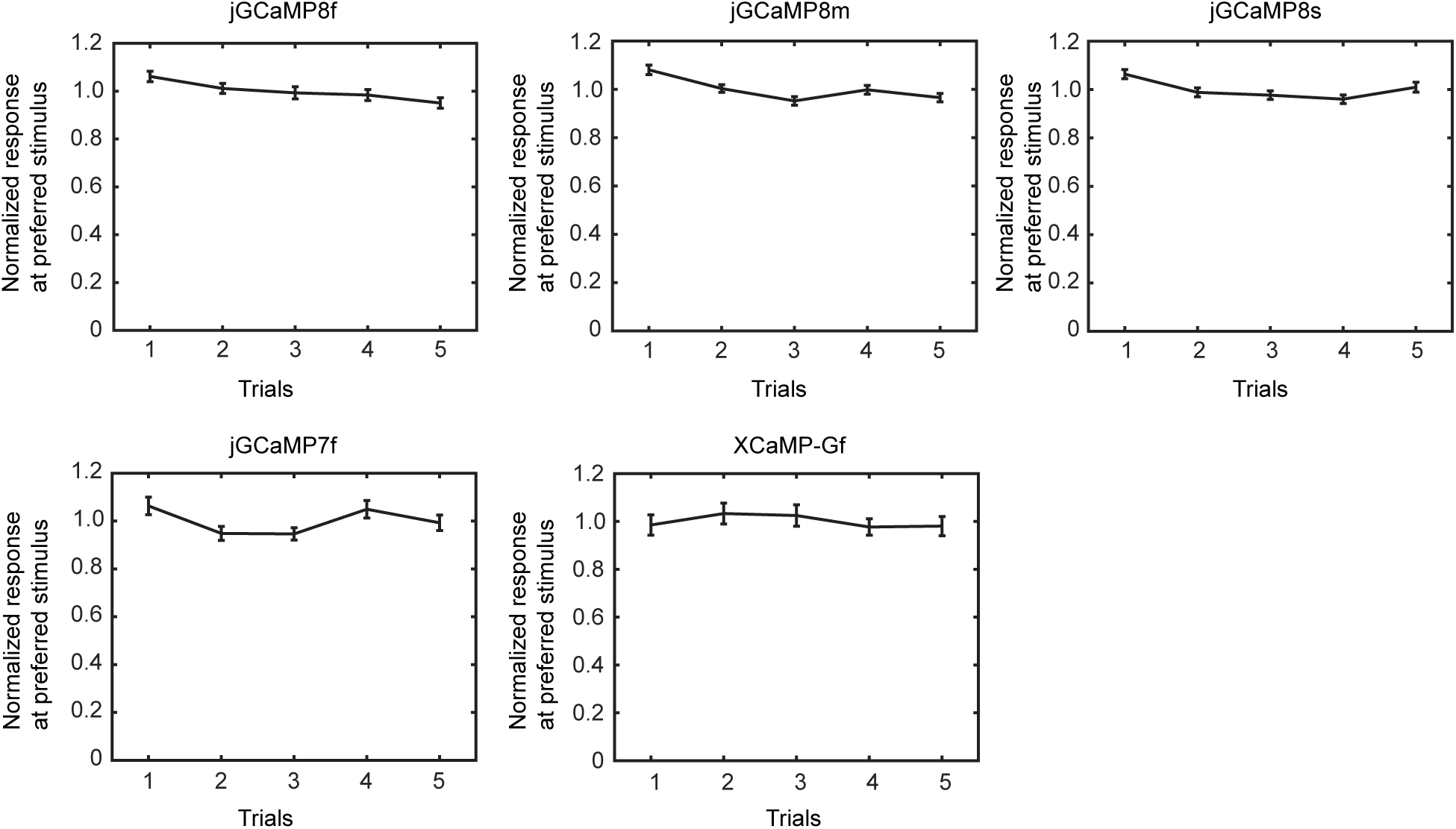
Reproducible responses across trials. The peak response amplitude of orientation selective neurons was averaged (8f, 288 neurons; 8m, 305 neurons; 8s, 420 neurons; 7f, 269 neurons; XCaMP-Gf, 121 cells) and plotted as a function of trial number.

**Supp. Fig. 13.**
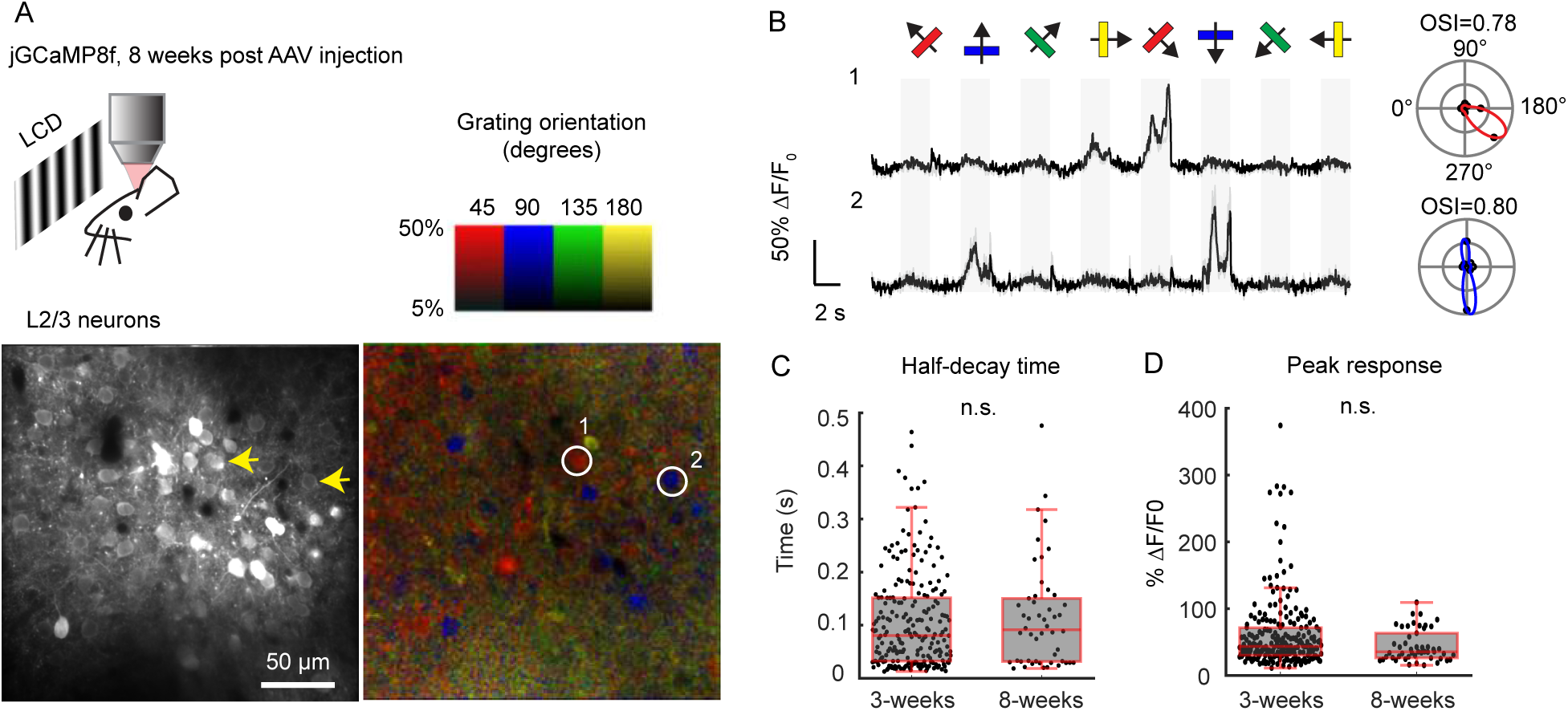
Response comparison between 3 weeks and 8 weeks post-AAV infection. A. Top, schematic of the experiment. Bottom, image of V1 L2/3 cells expressing 8f eight weeks post-AAV injection (left), and the same field of view color-coded according to the neurons’ preferred orientation (hue) and response amplitude (brightness). B. Example traces from two L2/3 neurons in A. Light traces: five individual trials; dark traces: mean. Eight grating motion directions are indicated by arrows and shown above traces. The preferred stimulus is the direction evoking the largest response. Polar plots indicate the preferred orientation or direction of the cells. OSI values displayed above each polar plot. C. Box-plot comparison of half-decay time for 8f between data acquired at 3 weeks and 8 weeks post-AAV injection. 225 cells from 6 mice for 3 weeks’ data; 50 cells from 2 mice for 8 weeks’ data. Two-sided Wilcoxon rank-sum test, *P* = 0.60. D. Comparison of peak response for 8f between data acquired at 3 weeks and 8 weeks post-AAV injection. 225 cells from 6 mice for 3 weeks’ data; 50 cells from 2 mice for 8 weeks’ data. Two-sided Wilcoxon rank-sum test, *P* = 0.053.

**Supp. Fig. 14.**
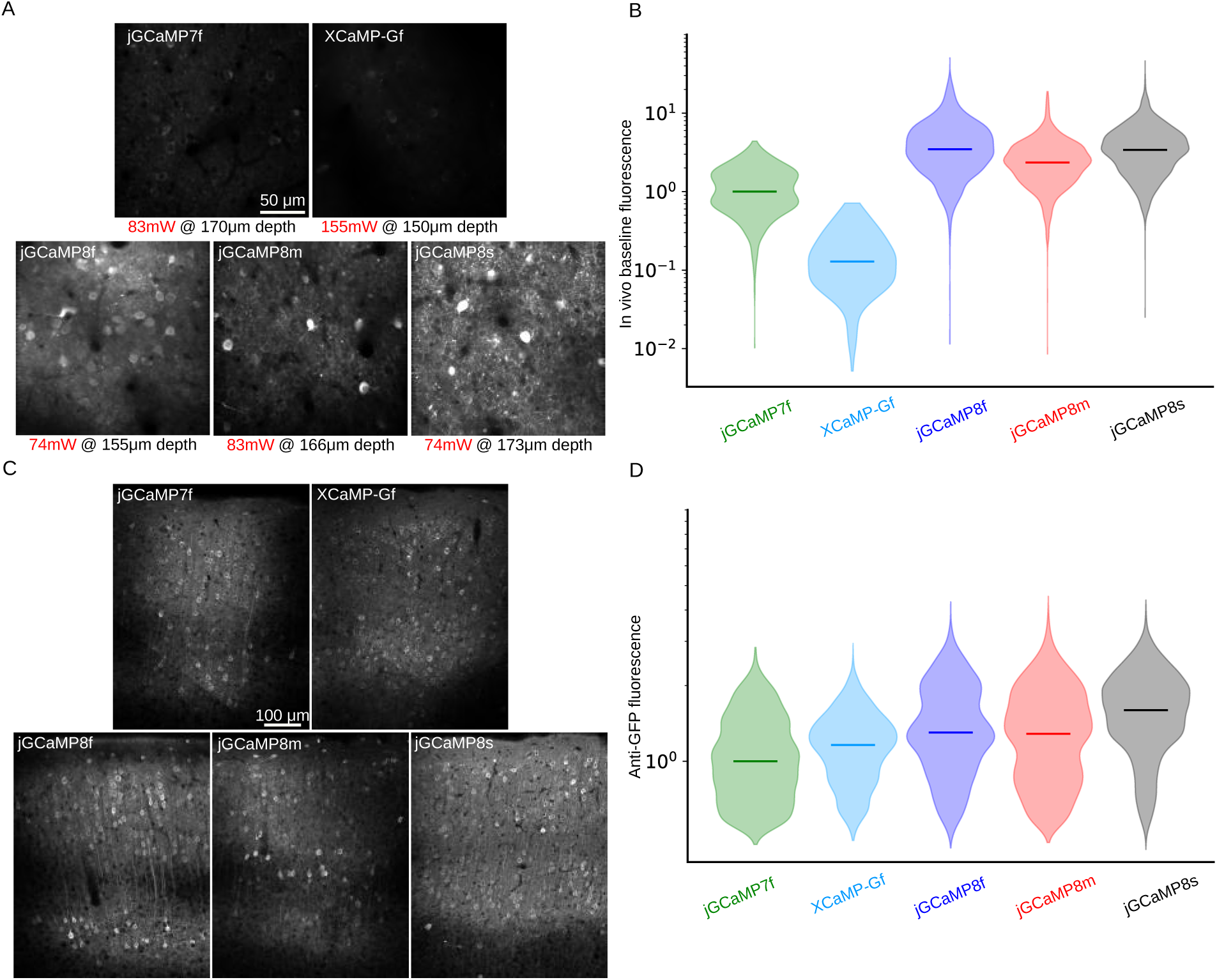
Sensor brightness *in vivo* and expression level. A. Representative *in vivo* movie averages for all GECIs. The post-objective illumination power and the depth of imaging is noted under each image. The color axis is the same for all images. B. *In vivo* distribution of excitation power-corrected baseline fluorescence values for segmented cellular ROIs. Horizontal bars represent the median of each distribution. Note the logarithmic scale. All data are normalized to the median of the 7f distribution. See panel A for representative motion corrected *in vivo* two-photon movie averages. C. Representative images of anti-GFP fluorescence for all GECIs in a coronal section across the center of an injection site, 20-22 days post injection. The color axis is the same for all images. D. Distribution of somatic fluorescence values of anti-GFP antibody labelling for all sensors, 20-22 days post injection. Horizontal bars represent the median of each distribution. All data is normalized to the median of the 7f values. Note that the expression levels are similar across sensors. The data is collected from two mice for each sensor. See panel C for representative images.

**Supp. Fig. 15.**
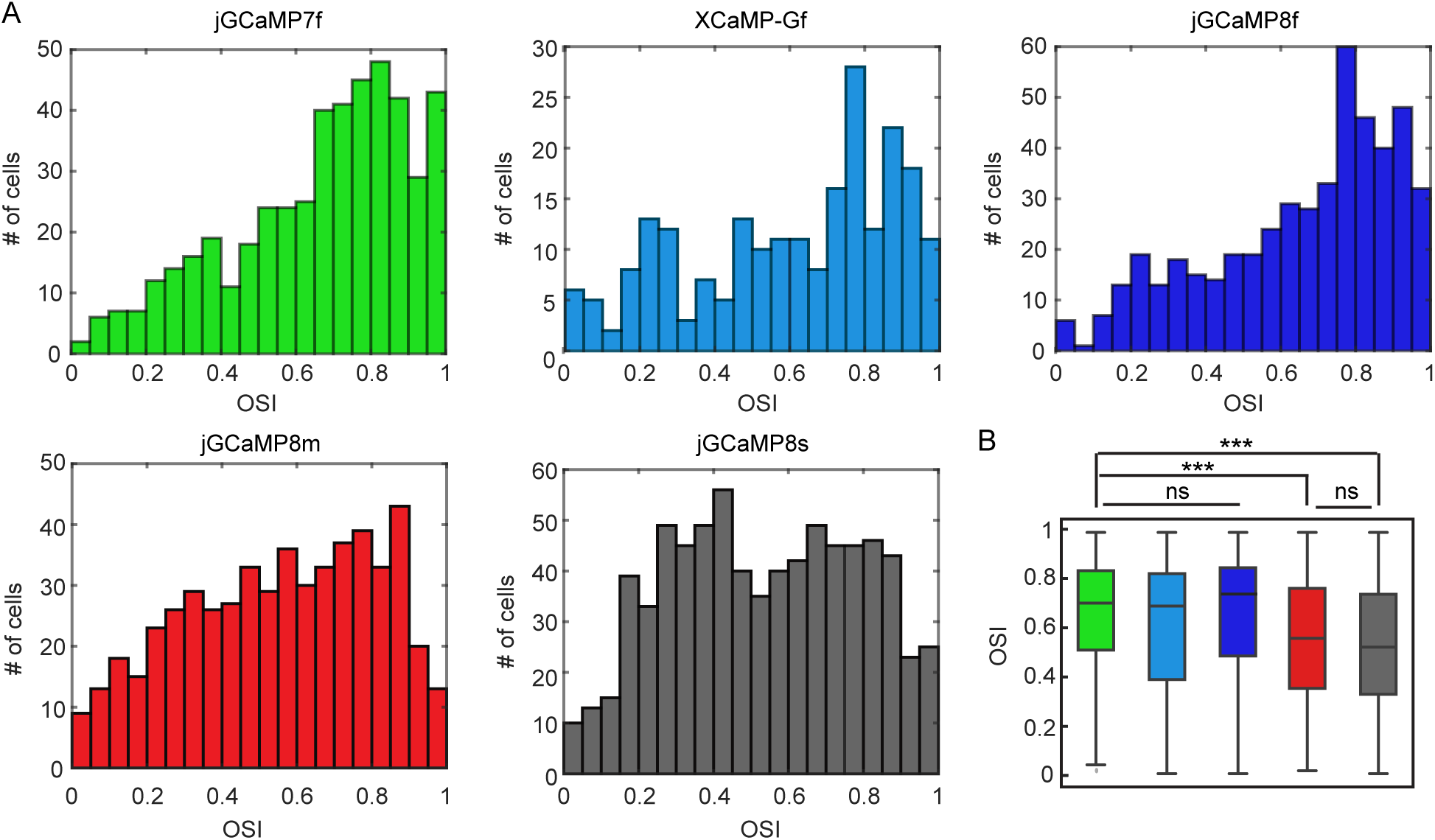
Orientation selectivity of the GCaMP-expressing mice. A. Distribution of orientation selectivity index (OSI) for visually responsive cells measured using different sensors (n = 473 cells, 7f; 221, XCaMP-Gf; 484, 8f; 532, 8m; 742, 8s). There is a noticeable left shift in the distributions of OSI for 8m and 8s. B. Comparison of OSI values across sensors (same data as in A). Kruskal-Wallis test (*P* < 0.001) with Dunn’s multiple comparison test was used for statistics. 7f *vs* XCaMP-Gf: *P* = 0.13; 7f *vs* 8f: *P* = 1.0; 7f *vs* 8m; *P* < 0.001; 7f *vs* 8s; *P* < 0.001; 8m *vs* 8s: *P* = 1.0. \*\*\**P* < 0.001. ns, not significant.

**Supp. Fig. 16.**
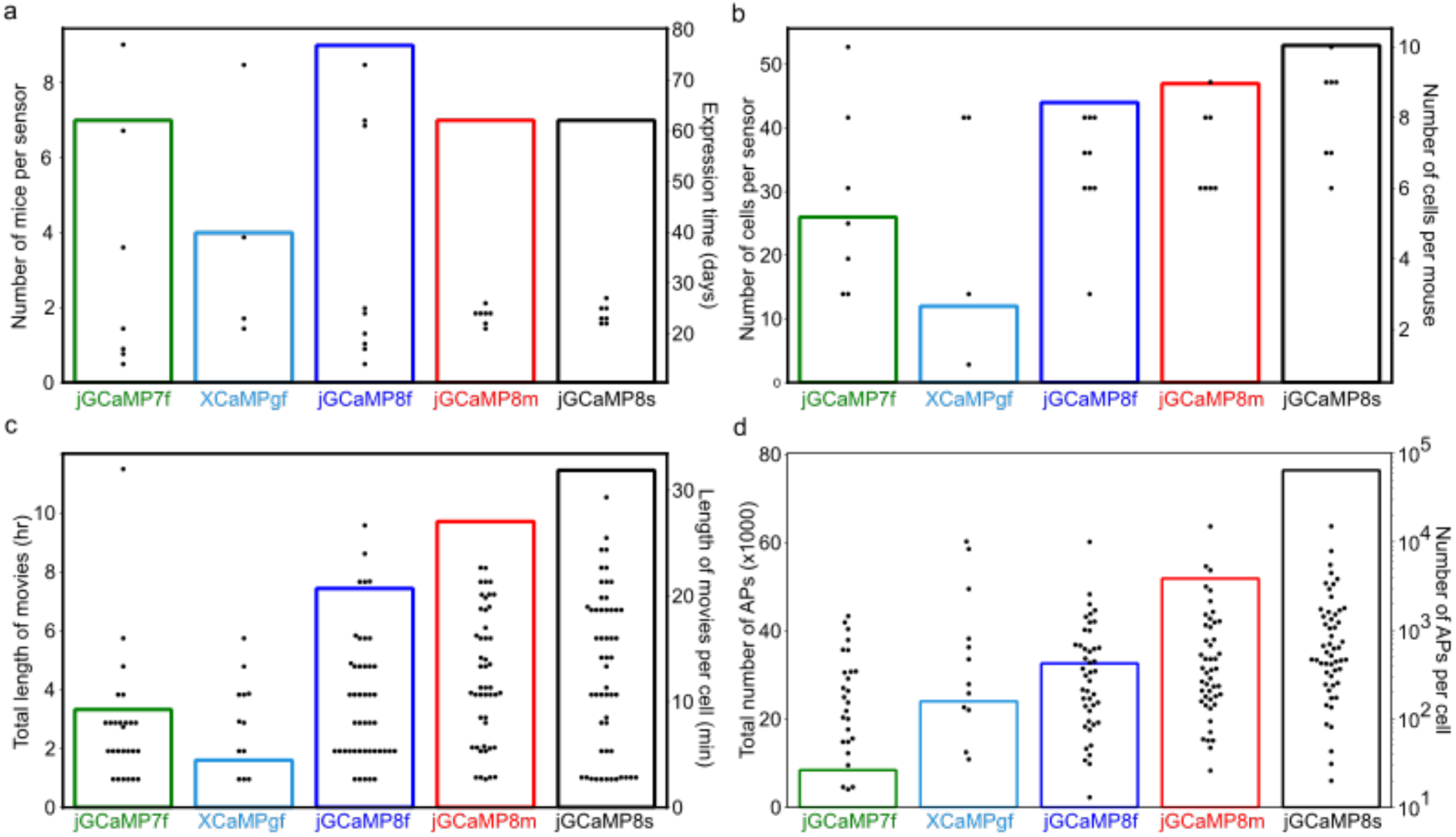
Descriptive statistics for loose-seal cell-attached recordings. A. Summary plot showing the number of mice used (bars, left y-axis) and the expression time at the time of the loose-seal recording in days (dots, right y-axis), for each sensor. B. Summary plot showing the total number of cells recorded (bars, left y-axis), and the number of cells recorded per mouse (dots, right y-axis) for each sensor. C. Summary plot showing the total length of simultaneous imaging and loose-seal recordings in hours (bars, left y-axis), and the length of simultaneous imaging and loose-seal recordings in minutes for each cell (dots, right y-axis). D. Summary plot showing the total number of action potentials (bars, left y-axis), and the number of recorded action potentials for each cell (dots, right y-axis – log scale), for each sensor.

**Supp. Fig. 17.**
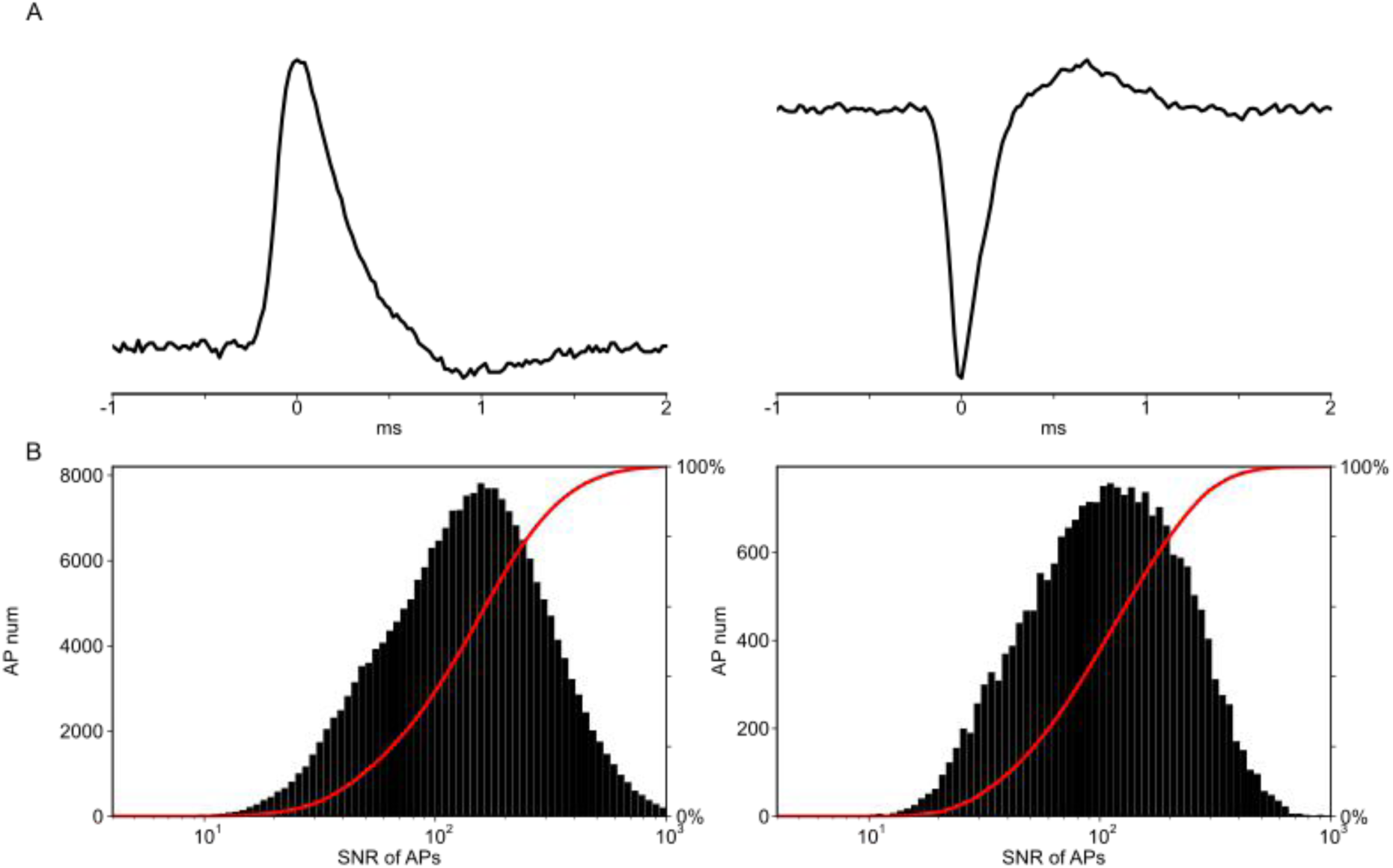
Signal-to-noise ratio of action potential recordings. A. Representative waveforms of loose-seal recorded action potentials in current-clamp (left) and voltage-clamp (right) recording mode. B. Signal-to-noise ratio distribution for all recorded action potentials in current-clamp (left) and voltage-clamp (right) recording mode.

**Supp. Fig. 18.**
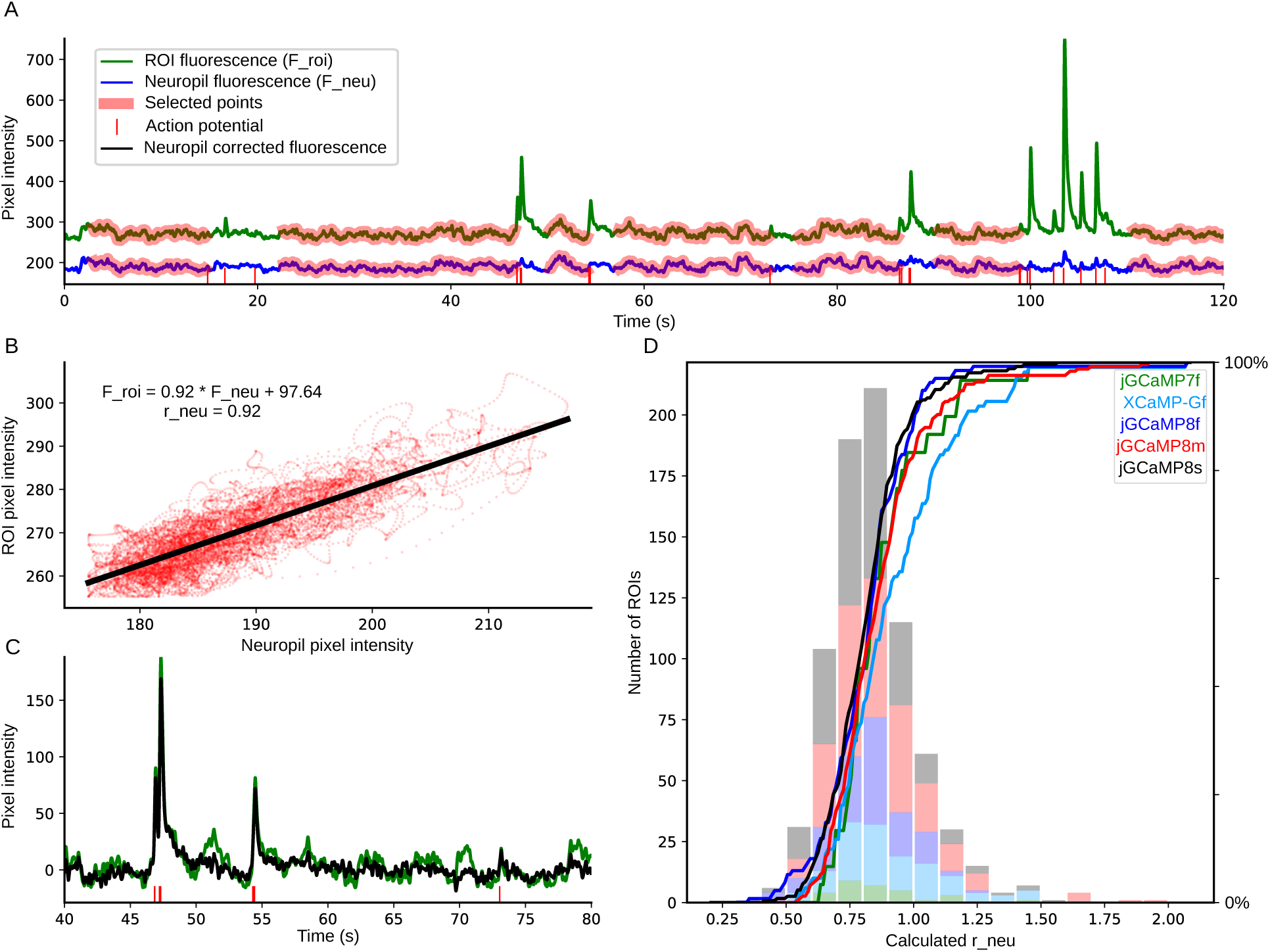
Sensor fluorescence across cell body ROIs and neuropil. A. A representative fluorescence trace for a cellular ROI (green) and its surrounding neuropil (blue) with simultaneous loose-seal recording. For calculating the distribution of neuropil contamination coefficients (r_neu), time points during the 3 seconds after an electrophysiologically recorded action potential (red vertical bars) were not included. Time points included in the analysis are highlighted in red. Note the correlation between cellular and neuropil ROI. Traces were high-pass filtered using a 10-second-long minimum filter and low-pass filtered with a Gaussian filter (σ = 10 ms). B. Cellular ROI pixel intensity values plotted against their corresponding neuropil pixel intensity values (time points highlighted with red on panel A), and their linear fit. The neuropil contamination coefficient is defined as the slope of this fitted function. C. Raw and neuropil corrected trace from panel A (40-80 sec), corrected with the neuropil contamination coefficient calculated in panel B (F_corr = F_roi -r_neu*F_neu). D. Distribution of r_neu values, each calculated on 3-minute-long simultaneous optical and electrophysiological recordings as shown in panels A-B. We included r_neu values only with a Pearson’s correlation coefficient > 0.7. Colors represent different GECIs. Calculated values of r_neu were similar between GECIs except for XCaMP-Gf, which was quite dim.

**Supp. Fig. 19.**
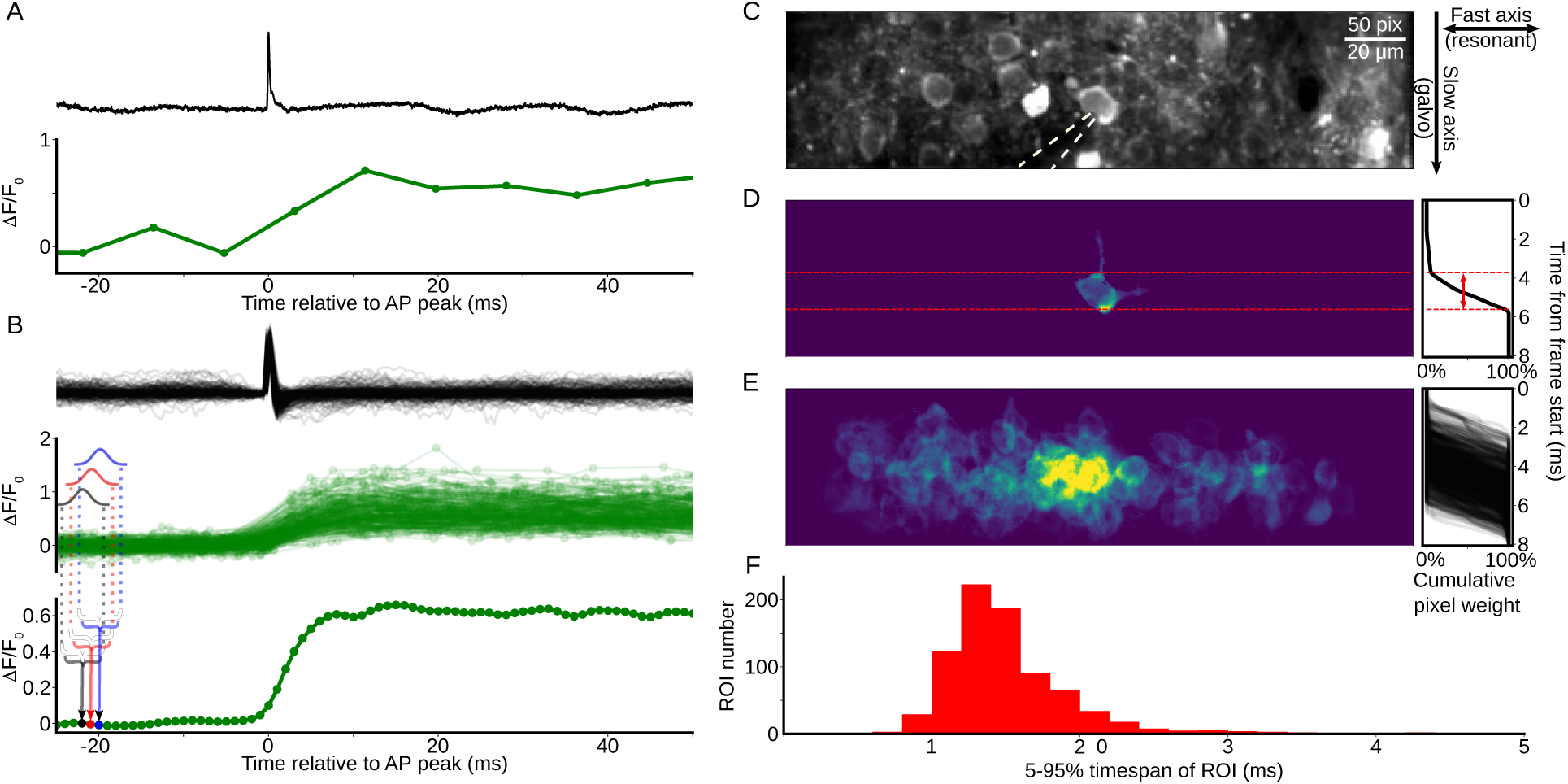
Effective ∼500 Hz imaging of fluorescent responses *in vivo*. A. Example isolated action potential during a simultaneous loose-seal recording at 50 kHz (top panel) and imaging at 122 Hz (bottom panel) of an 8s-expressing neuron. B. Same as in A but 250 isolated action potentials are aligned to the peak of the action potential and overlaid. Note that frame times (green dots in middle panel) are uniformly distributed in time. Bottom, construction of the high-resolution resampled trace. Each point in the resampled trace is generated by averaging the surrounding time points across the population of calcium transients with a Gaussian kernel. Three example points are highlighted with black, red and blue colors, together with the time span and weight used for the calculation of each point. C. Mean intensity projection of a representative field of view during cell-attached loose-seal recording. Recording pipette is highlighted with dashed white lines. The right panel shows how each frame is generated: the horizontal axis is scanned with a resonant scan mirror, the speed of which can be considered instantaneous relative to the vertical axis. The vertical axis is scanned with a slower galvanometer mirror, the speed of which determines the frame rate. D. Cellular ROI of the loose-seal recorded cell in panel C. Color scale shows pixel weights for ROI extraction. Right: cumulative pixel weight over the generation of a frame. We defined the timespan of the ROI as the 5-95% time of the cumulative pixel weight function. The timespan of the ROI is denoted with a red two-headed arrow. E. All loose-seal recorded ROIs weights overlaid as in panel D. An ROI was defined from three-minute-long movies, so a single recorded cell can have multiple overlapping ROIs in this image. F. Distribution of 5-95% timespans of all recorded ROIs. The timespans of most ROIs are under two milliseconds – thus the upper bound of the temporal resolution is ∼500 Hz.

**Supp. Fig. 20.**
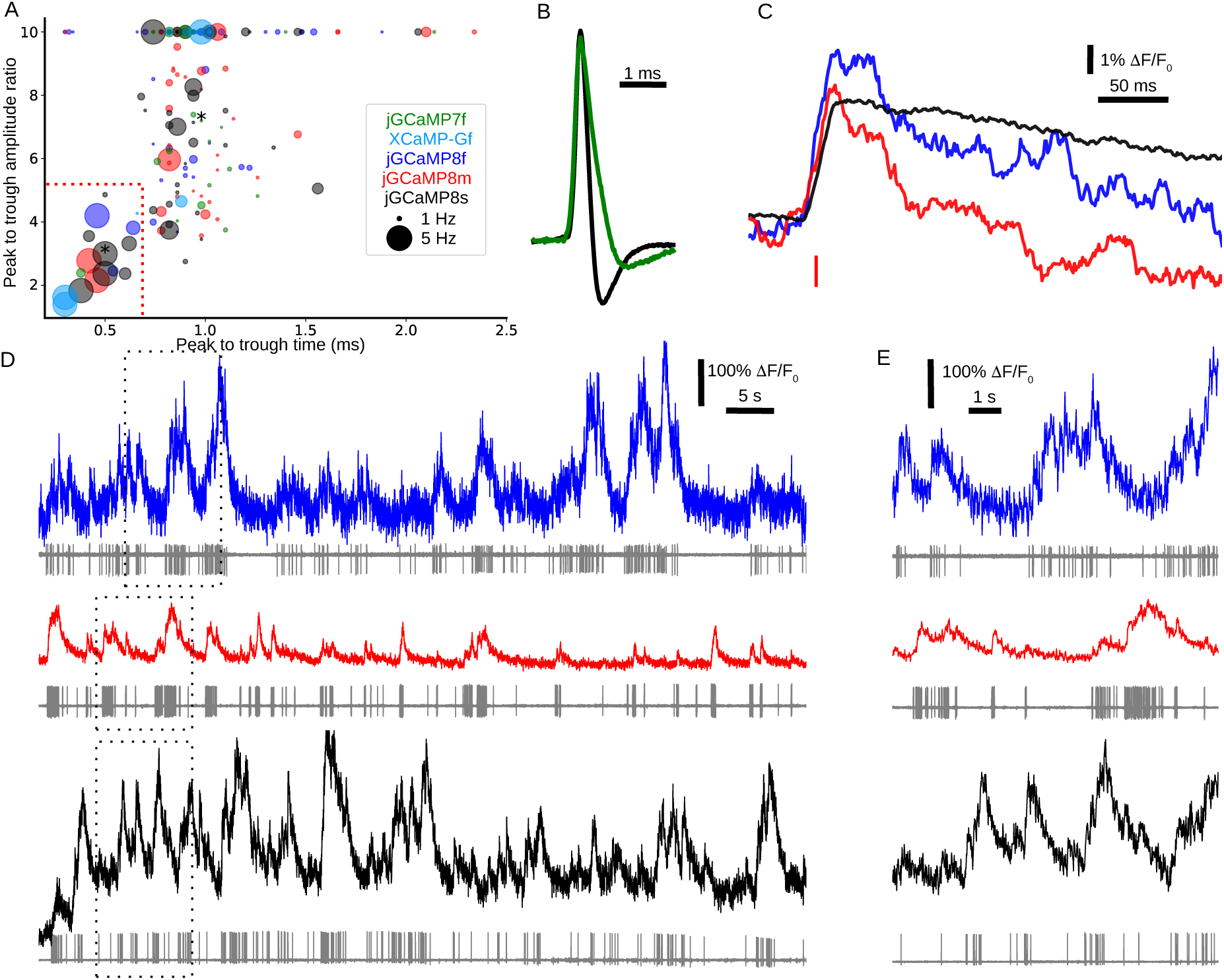
Responses in fast-spiking interneurons. A. Spike waveform parameters for each recorded cell; colors represent the expressed sensor, and the size of the circle represents average firing rate. Peak-to-trough ratios larger than 10 are plotted as 10. We defined putative interneurons as cells occupying the lower left quadrant (short peak-to-trough time and low peak-to-trough amplitude ratio), borders highlighted with red dotted lines. B. Example average action potential waveforms of a putative fast-spiking cell (black) and a putative pyramidal cell (green). The corresponding cells are marked with asterisks in panel A. C. Average calcium transient waveform for a single action potential in putative interneurons for 8f, 8m, and 8s. Resampling was done with a 20-ms-long mean filter. D. Simultaneous fluorescence dynamics and spikes in 8f (top), 8m (middle) and 8s (bottom) expressing putative interneurons. Fluorescence traces were filtered with a Gaussian filter (σ = 5 ms). E. Zoomed-in view of bursts of action potentials from panel C (top, 8f; middle, 8m; bottom, 8s).

**Supp. Fig. 21.**
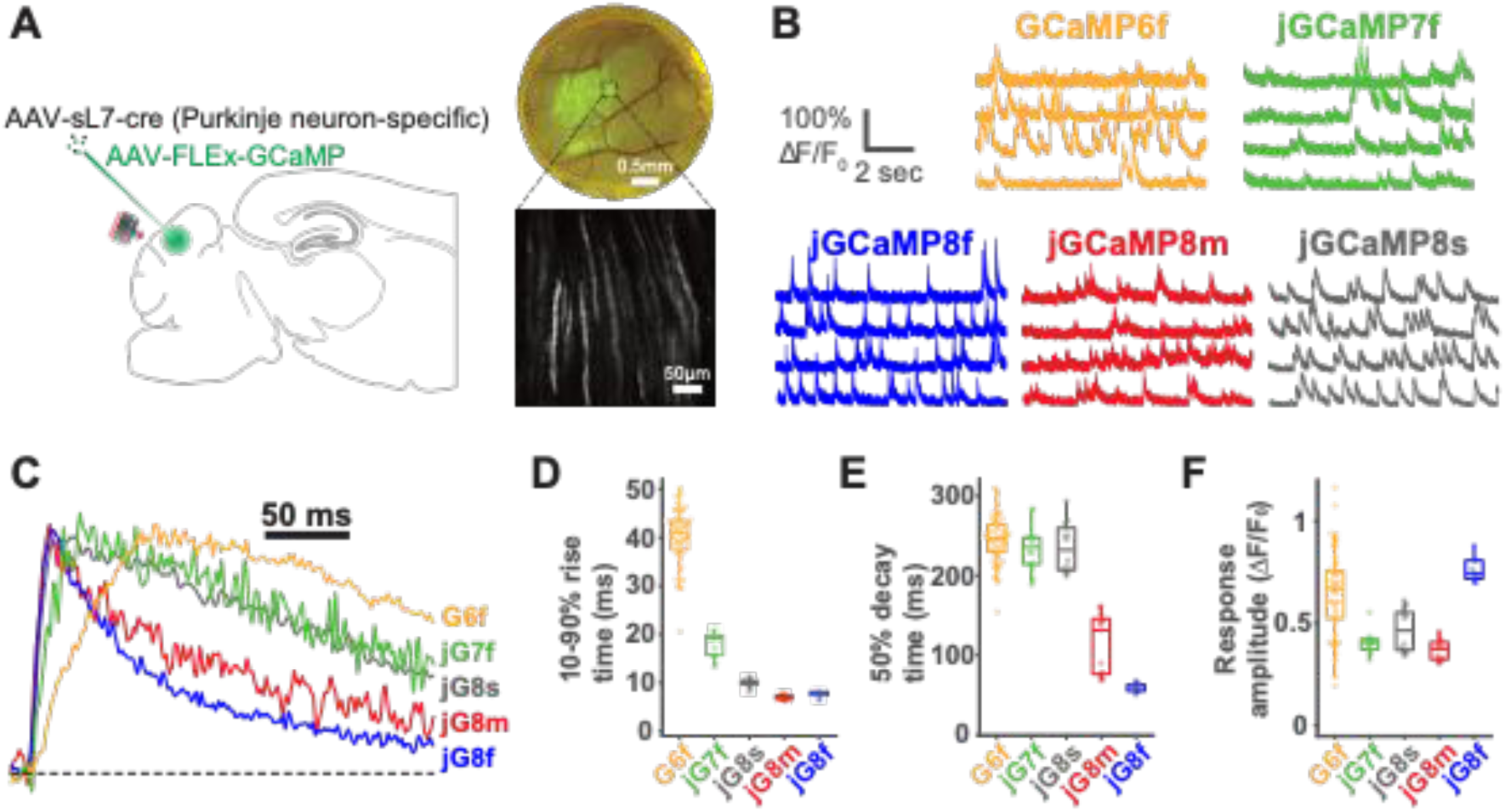
Imaging simple and complex spikes in cerebellar Purkinje neurons. A. Experimental design. Purkinje neurons in cerebellar lobule VI were transduced with a GCaMP variant as in the sample widefield (top right) and 2P (bottom right) images. Dendritic tufts were monitored for complex spike-related activity using 2P microscopy under free-locomotion conditions. B. Sample traces from adjacent dendrites for each variant. C. Normalized fluorescence traces from the average of 10 events nearest to the median values from each variant. D. Time to rise from 10-90% of baseline fluorescence. E. Half-decay time of the fluorescence signal following the peak fluorescence value. F. Distribution of ΔF/F_0_ response to complex spike. For each variant, 2 mice were imaged with number of dendrites per variant as: 6f, n = 51; 7f, n = 14; 8s, n = 14; 8m, n = 13; 8f, n = 9. Mean ± s.e.m. shown.

**Supp. Fig. 22.**
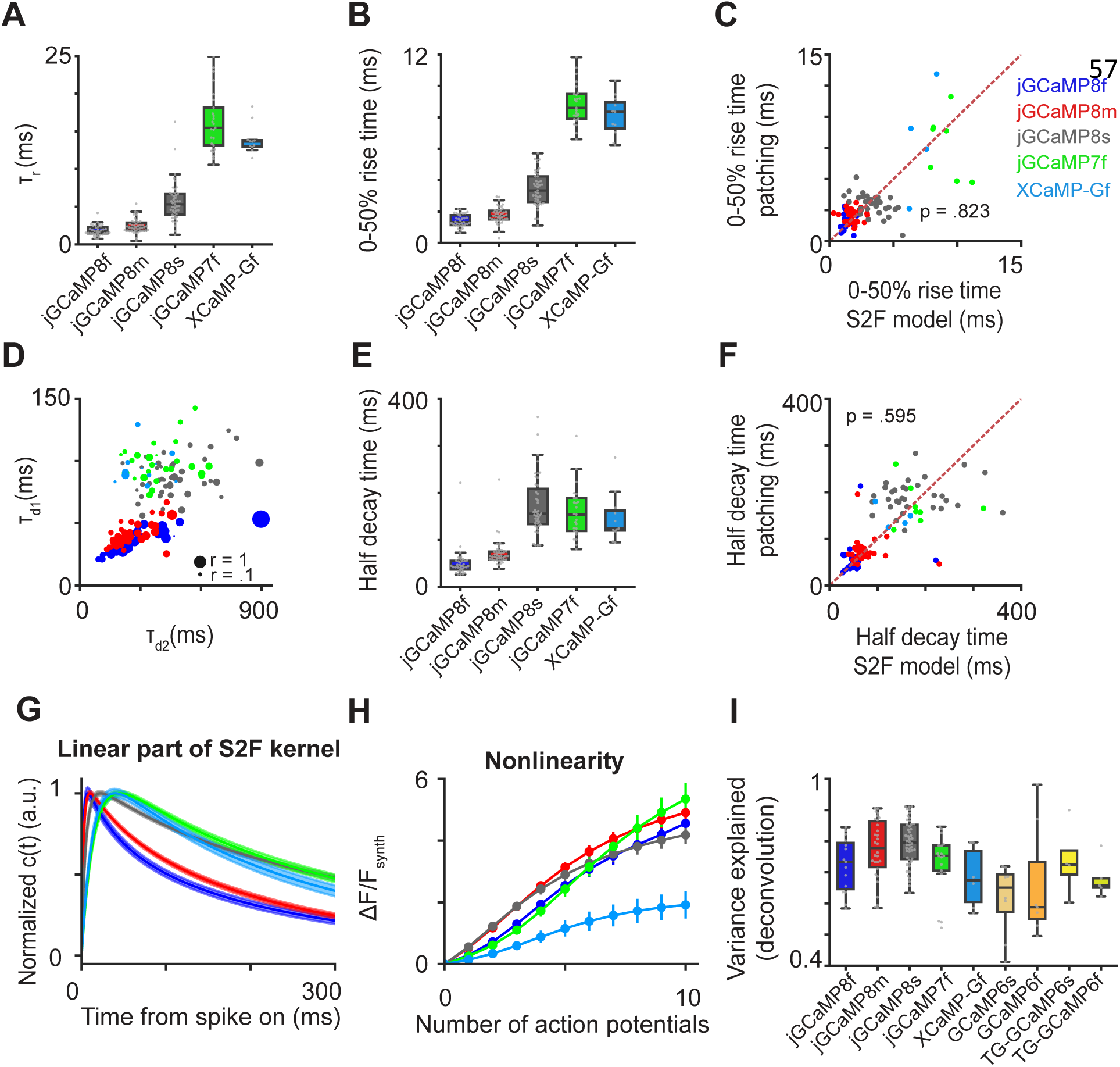
Statistics of S2F fits in the different imaging conditions. A-F. Statistics of S2F fits in the different imaging conditions (See **Supp. Table 7** for more details). Blue, 8f; red, 8m; dark gray, 8s; green, 7f; cyan, XCaMP-Gf. B. Boxplots of rise time, **τ**r. C. Boxplots of 0-50% peak rise-time derived from S2F fits. D. Comparison between 0-50% peak rise time derived from S2F fits (x-axis) with that measured by super-resolution patch data (y-axis). Red dashed line is the identity line. E. Scatter plots of decay times. X-axis, the slow decay time, **τ**_d2_; y-axis, the fast decay time, **τ**_d1_; size of dots, the ratio of the weight for fast decay time to that for the slow one, r. F. Box-plots of half-decay time derived from S2F fits. G. Comparison between half-decay time derived from S2F fits (x-axis) with that measured by super-resolution patch data (y-axis; see Fig. 4E for more details). Red dashed line is the identity line. G,H. ΔF/F_Synth_ simulated from the S2F models of different sensors. H. Normalized synthetic calcium latent dynamics, c(t); solid lines, mean; shaded area, s.e.m. I. Simulated peak nonlinearity, *i.e.*, synthetic fluorescence response to different numbers of action potentials. Error bars, s.e.m. across cells. J. Performance of the deconvolution algorithm (from fluorescence to spikes) on various indicators. x-axis, indicators; y-axis, variance explained, *i.e.*, the extent to which fluorescence dynamics can be fitted by the inferred spikes from deconvolution – the higher the variance explained, the better the deconvolution algorithm infers the spike timing and calcium kinetics from fluorescence. The deconvolution is based on a widely used algorithm from ^38^ with two decay constants.

