## Supplemental Table 1 for "Fast and sensitive GCaMP calcium indicators for imaging neural populations"

**Supplementary Table 1**

| PDB ID /peptide name | Peptide sequence | App. K_d_, nM | Satur. ΔF/F_0_ | Hill  coefficient | k_off1_(s^-1^) | k_off2_(s^-1^) | Normalized F_0_ |
| --- | --- | --- | --- | --- | --- | --- | --- |
| 1CDL | ARRKWQKTGHAVRAIGRLSS | 205 | 50 | 2.1 | 1.87 | N/A | 1.18 |
| 1CDM | LKKFNARRKLKGAILTTMLATRNFS | 728 | 14.5 | 2 | 6.03 | 5.76 | 1.80 |
| 1IQ5 | VRVIPRLDTLILVKAMGHRKRFGNPFR | N/A | N/A | N/A | N/A | N/A | 5.13 |
| 1IWQ | KKRFSFKKSFKLSGFSFKK | 3436 | 10.6 | 1.4 | N/A | N/A | 2.17 |
| 1NIW | RKKTFKEVANAVKISASLMG | 1062 | 14.8 | 2 | 23.01 | 0.12 | 1.42 |
| 1SY9 | GGFRRIARLVGVLREWAYR | 67 | 11.9 | 2.7 | 8.13 | 0.45 | 0.91 |
| 1YR5 | RKKWKQSVRLISLCQRLSR | 205 | 18 | 1.8 | 15.01 | 0.39 | 2.19 |
| 2BCX | KSKKAVWHKLLSKQRRRAVVACFRM | 3288 | 5.8 | 2 | N/A | N/A | 3.08 |
| 2F3Y | KFYATFLIQEYFRKFKK | N/A | N/A | N/A | N/A | N/A | 1.98 |
| 2FOT | ASASPWKSARLMVHTVATFNSIKER | 34.2 | 7.4 | 2.1 | 1.92 | 0.057 | 1.68 |
| 2HQW | KKKATFRAITSTLASSFKR | 682 | 29.9 | 1.7 | 5.01 | 0.43 | 1.65 |
| 2KNE | LRRGQILWFRGLNRIQTQIKVVKAFHS | 38.1 | 11 | 1.4 | 1.99 | N/A | 1.82 |
| 2LGF | AFIIWLARRLKKGKK | N/A | N/A | N/A | N/A | N/A | 1.87 |
| 2M55 | MDVFMKGLSKAKEGVVAAA | N/A | N/A | N/A | N/A | N/A | 0.97 |
| 2MES | MDCLCIVTTKKYRYQD | N/A | N/A | N/A | N/A | N/A | 0.29 |
| 2N6A | AAGSGWRKIKLAVRGAQAK | N/A | N/A | N/A | N/A | N/A | 1.52 |
| 2O60 | KRRAIGFKKLAEAVKFSAKLMG | 653 | 16.6 | 1.7 | 24.5 | 0.68 | 1.57 |
| 2VAY | KFYATFLIQEHFRKFMKRQEE | 1814 | 2.2 | 1.1 | N/A | N/A | 0.93 |
| 3BXX | KIYAAMMIMEYYRQSKAKKLQ | 615 | 3.5 | 1.9 | N/A | N/A | 1.66 |
| 3EWT | SKYITTIAGVMTLSQV | 5931 | 17.9 | 1 | N/A | N/A | 0.35 |
| 3GOF | RRREIRFRVLVKVVFFSS | 490 | 41.6 | 1.8 | 1.89 | N/A | 0.26 |
| 3GP2 | SFNARRKLKGAILTTMLATAS | 1523 | 31.4 | 1.6 | N/A | N/A | 0.66 |
| 3SUI | GRVSGRNWKNFALVPLLRDAS | N/A | N/A | N/A | N/A | N/A | 1.41 |
| 4AQRA | ERLQQWRKAALVLNASRRFRY | 420 | 22 | 1.8 | 2.82 | N/A | 1.52 |
| 4AQRB | REMRQKIRSHAHALLAANRFMDM | 865 | 16 | 1.9 | 6.04 | 1.15 | 0.92 |
| 4Q5U | ARKEVIRNKIRAIGKMARVFSVLR | 705 | 17.1 | 1.8 | 28.9 | 1.32 | 1.17 |
| 4UPU | NHWQKIRTMVNLPVISPFKSS | 13000 | N/A | N/A | N/A | N/A | 1.60 |
| 5DOW | KRNKALKKIRKLQKRGLIQMT | N/A | N/A | N/A | N/A | N/A | 0.55 |
| RS20^*^ | SSRRKWNKTGHAVRAIGRLSS | 131 | 56.5 | 2.2 | 1.09 | N/A | 1.00 |
| CKKAP^†^ | VKLIPSLTTVILVKSMLRKRSFGNPF | N/A | N/A | N/A | N/A | N/A | 1.91 |
| 6GGS^‡^ | GGSGGSGGSGGSGGSGGS | N/A | N/A | N/A | N/A | N/A | 1.48 |

Biophysical properties of initial sensors with different calmodulin-binding peptides used in this study.

Biophysical properties were measured in purified protein solutions. The positive control sensor has ^*^ RS20, similar to GCaMP6s with a shorter tag (**Methods**). ^†^ CKKAP (CaM-dependent kinase kinase peptide) was used in XCaMP ^23^. The negative control ^‡^ 6GGS has 6 sequential Gly-Gly-Ser in place of the CaM-binding peptide.
