## Supplemental Table 3 for "Fast and sensitive GCaMP calcium indicators for imaging neural populations"

**Supplementary Table 3:** **Data collection and refinement statistics of jGCaMP8.410.80**

|  | jGCaMP8.410.80 |
| --- | --- |
| **Data collection** |  |
| Space group | P4_1_2_1_2 |
| Cell dimensions |  |
| *a*, *b*, *c* (Å) | 120.6, 120.6, 97.9 |
| α, β, γ (°) | 90.0, 90.0, 90.0 |
| Resolution (Å) | 47.24-2.00 (2.05-2.00)* |
| *R*_sym_ or *R*_merge_ | 0.056 (0.496)* |
| *I* / σ*I* | 31.2(5.6)* |
| Completeness (%) | 100.0 (100.0)* |
| Redundancy | 14.3 (12.9)* |
| **Refinement** |  |
| Resolution (Å) | 45.49-2.00 |
| No. reflections all/free | 49229/2378 |
| *R*_work_ / *R*_free_ | 0.175/0.197 |
| No. atoms |  |
| Protein | 2980 |
| Ligand | 38 |
| Ion | 4 |
| Water | 243 |
| *B*-factors |  |
| Protein | 35.88 |
| Ligand | 40.40 |
| Ion | 34.72 |
| Water | 39.97 |
| R.m.s. deviations |  |
| Bond lengths (Å) | 0.0134 |
| Bond angles (°) | 1.732 |

*Values in parentheses are for highest-resolution shell.
