## Supplemental Table 4 for "Fast and sensitive GCaMP calcium indicators for imaging neural populations"

**Supp. Table 3-NS_2.** Comparison of sensitivity and kinetics of jGCaMP8 to XCaMP-G, -Gf, and -Gf0 sensors. Colors in each cell indicate whether the value was significantly higher for jGCaMP8 (yellow), XCaMP (blue), or not statistically different (no color), as evaluated with Dunn’s multiple comparisons test (*P*-values in cells).

| **ΔF/F**  (yellow: jGCaMP8 is higher, blue: XCaMP is higher) | | | |
| --- | --- | --- | --- |
| **Kruskal-Wallis test: p = 6.939649865339752e-30** | | | |
|  | **XCaMP-G** | **XCaMP-Gf** | **XCaMP-Gf0** |
| **jGCaMP8f** | 4.773183e-06 | 3.406466e-06 | 2.982465e-12 |
| **jGCaMP8m** | 3.749453e-06 | 3.496553e-06 | 1.475564e-09 |
| **jGCaMP8s** | 1.392821e-13 | 8.237702e-14 | 2.235685e-20 |

| **SNR**  (yellow: jGCaMP8 is higher, blue: XCaMP is higher) | | | |
| --- | --- | --- | --- |
| **Kruskal-Wallis test: p = 3.1161722160256525e-26** | | | |
|  | **XCaMP-G** | **XCaMP-Gf** | **XCaMP-Gf0** |
| **jGCaMP8f** | 1.000000e+00 | 4.914442e-03 | 1.029269e-07 |
| **jGCaMP8m** | 4.925522e-04 | 2.719804e-06 | 1.092976e-09 |
| **jGCaMP8s** | 1.189540e-09 | 1.824282e-14 | 3.741053e-21 |

| **Half-rise time**  (yellow: XCaMP faster, blue: jGCaMP8 faster) | | | |
| --- | --- | --- | --- |
| **Kruskal-Wallis test: p = 3.7785355253128484e-41** | | | |
|  | **XCaMP-G** | **XCaMP-Gf** | **XCaMP-Gf0** |
| **jGCaMP8f** | 2.247699e-35 | 1.171520e-15 | 3.193351e-17 |
| **jGCaMP8m** | 2.811176e-11 | 1.316382e-04 | 3.002614e-05 |
| **jGCaMP8s** | 1.439711e-12 | 1.328715e-03 | 2.222932e-04 |

| **Time to peak**  (yellow: XCaMP faster, blue: jGCaMP8 faster) | | | |
| --- | --- | --- | --- |
| **Kruskal-Wallis test: p = 1.5329660909865355e-32** | | | |
|  | **XCaMP-G** | **XCaMP-Gf** | **XCaMP-Gf0** |
| **jGCaMP8f** | 1.299251e-29 | 2.178685e-13 | 3.413239e-18 |
| **jGCaMP8m** | 2.075880e-08 | 2.739457e-03 | 7.322556e-05 |
| **jGCaMP8s** | 3.490259e-05 | 1.000000e+00 | 1.051670e-01 |

| **Half-decay time**  (yellow: XCaMP faster, blue: jGCaMP8 faster) | | | |
| --- | --- | --- | --- |
| **Kruskal-Wallis test: p = 1.4002014632305323e-38** | | | |
|  | **XCaMP-G** | **XCaMP-Gf** | **XCaMP-Gf0** |
| **jGCaMP8f** | 6.465057e-26 | 4.979016e-10 | 1.522713e-15 |
| **jGCaMP8m** | 3.159566e-06 | 1.035035e-01 | 2.784363e-03 |
| **jGCaMP8s** | 1.825883e-02 | 2.556897e-10 | 2.792187e-06 |
