## Supplemental Table 7 for "Fast and sensitive GCaMP calcium indicators for imaging neural populations"

**Table SX1. Statistics of the degree of nonlinearity of sensors measured by the difference of variance explained by S2F sigmoid from linear model (mean ± std. dev.)**

| Sensors | Variance explained  (sigmoid)(%) | Variance explained  (linear)(%) | Variance explained  (sigmoid - linear)(%) | Number of cells |
| --- | --- | --- | --- | --- |
| **jGCaMP8f** | 79.06 ± 12.43 | 73.24 ± 11.78 | 5.82 ± 3.48 | 38 |
| **jGCaMP8m** | 86.56 ± 12.17 | 84.99 ± 12.16 | 1.57 ± 1.46 | 44 |
| **jGCaMP8s** | 79.86 ± 19.00 | 78.33 ± 19.68 | 1.53 ± 1.83 | 51 |
| **jGCaMP7f** | 83.32 ± 14.72 | 76.30 ± 13.30 | 7.03 ± 5.37 | 26 |
| **XCaMP-Gf** | 80.66 ± 18.82 | 77.50 ± 17.60 | 3.16 ± 2.46 | 12 |
| **GCaMP6f** | 85.14 ± 15.89 | 62.99 ± 21.90 | 22.14 ± 11.63 | 11 |
| **GCaMP6s** | 92.38 ± 7.25 | 82.24 ± 8.52 | 10.14 ± 4.77 | 9 |
| **TG-GCaMP6f** | 77.31 ± 13.32 | 64.96 ± 13.05 | 12.99 ± 10.32 | 20 |
| **TG-GCaMP6s** | 71.68± 21.73 | 67.81 ± 14.53 | 7.96 ± 7.15 | 22 |
