## Supplemental Table 8 for "Fast and sensitive GCaMP calcium indicators for imaging neural populations"

**Table SX2. Statistics of parameter fits (mean ± std.dev.)**

| Sensors | Rise time 𝝉_r_ (ms) | Fast decay time 𝝉_d1_ (ms) | Slow decay time 𝝉_d2_ (ms) | Weight, r | 0-50% peak time (ms) | Half decay time (ms) |
| --- | --- | --- | --- | --- | --- | --- |
| jGCaMP8f | 1.85 ± 0.69 | 34.07 ± 9.21 | 263.7 ± 155.22 | 0.48 ± 0.3 | 1.41 ± 0.41 | 51.77 ± 32.48 |
| jGCaMP8m | 2.46 ± 0.94 | 41.64 ± 8.9 | 245.8 ± 86.57 | 0.28 ± 0.1 | 1.77 ± 0.53 | 72.76 ± 29.52 |
| jGCaMP8s | 5.65 ± 2.7 | 86.26 ± 15.22 | 465.45 ± 146.38 | 0.19 ± 0.08 | 3.44 ± 1.11 | 173.38 ± 61.74 |
| jGCaMP7f | 16.21 ± 3.98 | 95.27 ± 16.26 | 398.22 ± 127.82 | 0.24 ± 0.07 | 8.84 ± 1.29 | 159.22 ± 54.73 |
| XCaMP-Gf | 13.93 ± 1.95 | 99.38 ± 13.84 | 312.85 ± 96.92 | 0.2 ± 0.12 | 8.11 ± 1.29 | 147.71 ± 52.81 |
